# Recurrent Breast Cancer Cells Depend on *De novo* Pyrimidine Biosynthesis to Suppress Ferroptosis

**DOI:** 10.1101/2025.11.17.688927

**Authors:** Kelley R. McCutcheon, Josh Wu, Yasemin Ceyhan Ozdemir, Brock J. McKinney, Sharan Srinivasan, Ayaha Itokawa, Oliver J. Newsom, Anna Vigil, Douglas B. Fox, Lucas B. Sullivan, James V. Alvarez

**Affiliations:** Translational Research Program, Public Health Sciences Division, Fred Hutchinson Cancer Center.; Human Biology Division, Fred Hutchinson Cancer Center.; Department of Pharmacology and Cancer Biology, Duke University School of Medicine.

**Author notes:** Corresponding author: James V. Alvarez, Fred Hutchinson Cancer Center, 1100 Fairview Ave N, M5-C803, Seattle WA 98109.

**Keywords:** Recurrent breast cancer, Pyrimidine metabolism, DHODH (dihydroorotate dehydrogenase), Ferroptosis, Nucleotide salvage, Metabolic vulnerability

## Abstract

Breast cancer recurrence remains a major clinical challenge, often associated with therapy resistance and altered metabolic states. To define metabolic vulnerabilities of recurrent disease, we performed a CRISPR knockout screen targeting 421 metabolic genes in paired primary and recurrent HER2-driven breast cancer cell lines. While both primary and recurrent tumors shared dependencies on core metabolic pathways, recurrent tumors exhibited selective essentiality for the *de novo* pyrimidine synthesis pathway, including *Cad*, *Dhodh*, and *Ctps*. Pharmacologic inhibition of the rate-limiting enzyme DHODH with BAY-2402234 selectively impaired the growth of recurrent tumor cells, while primary tumor cells were relatively resistant. BAY treatment robustly inhibited pyrimidine synthesis in all lines, but only recurrent cells underwent iron-dependent lipid peroxidation and ferroptotic cell death. Lipidomic profiling revealed enrichment of polyunsaturated ether phospholipids in recurrent cells, which may predispose them to ferroptosis. A sensitizer CRISPR screen in primary cells further identified nucleotide salvage and lipid metabolic pathways as modifiers of DHODH inhibitor sensitivity. Stable isotope tracing and nutrient depletion experiments showed that primary cells can compensate for DHODH inhibition through nucleotide salvage, whereas recurrent cells exhibit impaired salvage capacity, likely due to reduced expression of *Slc28*/*Slc29* nucleoside transporters. Together, these findings reveal that breast cancer recurrence is associated with increased dependence on *de novo* pyrimidine synthesis to suppress ferroptosis, highlighting a therapeutically actionable metabolic vulnerability in recurrent disease.

## Introduction

Breast cancer recurrence remains a major clinical challenge, responsible for the majority of breast cancer–related deaths. Despite significant advances in early detection and targeted therapies, many patients ultimately experience metastatic relapse, often years or even decades after their initial diagnosis (1). These recurrent tumors are typically more aggressive and therapy resistant than their primary counterparts, reflecting fundamental biological differences between the two disease states.

A growing body of work demonstrates that tumor cells that survive initial therapy – whether as drug-tolerant persisters, dormant disseminated cells, or therapy-resistant populations – undergo extensive metabolic reprogramming to withstand treatment pressure and hostile microenvironments (2–6). These populations adapt to niches marked by oxidative stress, nutrient deprivation, hypoxia, and immune surveillance, enabling long-term survival despite continued therapeutic or microenvironmental constraints (2,7,8). Such adaptive metabolic states not only promote recurrence and metastasis but also reveal therapeutic vulnerabilities that are distinct from those of treatment-naïve primary tumors. Defining these context-specific metabolic dependencies is therefore essential for developing strategies to eliminate residual disease and prevent relapse.

One metabolic pathway of particular interest is *de novo* pyrimidine biosynthesis. Pyrimidine nucleotides are essential for DNA replication and repair, and their synthesis is tightly linked to mitochondrial function through the rate-limiting enzyme dihydroorotate dehydrogenase (DHODH). DHODH inhibitors, including brequinar, leflunomide, and BAY-2402234, have shown antitumor activity in multiple preclinical settings (9–12), yet the role of pyrimidine metabolism in recurrent breast cancer remains poorly understood.

Recent studies have also uncovered intriguing connections between nucleotide metabolism and ferroptosis, an iron-dependent form of cell death driven by lipid peroxidation. We and others have shown that recurrent breast tumors are particularly sensitive to ferroptosis (13,14), though the upstream metabolic pathways that govern this vulnerability remain undefined. DHODH has been implicated in ferroptosis regulation in certain contexts (15–17), raising the possibility that pyrimidine synthesis may intersect with lipid peroxidation programs during tumor recurrence.

Here, we show that recurrent breast cancer cells acquire a selective dependence on *de novo* pyrimidine biosynthesis to suppress ferroptosis and maintain survival. Through CRISPR-based metabolic screening, pharmacologic perturbation, and metabolite analysis, we reveal that recurrent tumors are acutely sensitive to DHODH inhibition, whereas primary tumor cells can compensate through nucleotide salvage. These findings identify pyrimidine metabolism as a therapeutically actionable vulnerability in recurrent breast cancer and provide new mechanistic insight into the metabolic rewiring that underlies recurrence.

## Results

### CRISPR screen identifies essential metabolic genes in primary and recurrent breast cancer cells

To identify metabolic dependencies specific to recurrent breast cancer, we performed a CRISPR knockout screen with a gRNA library targeting 421 metabolic genes (4 sgRNAs per gene; Table S1) in two primary tumor–derived cell lines (54074 and 99142, referred to as Primary 1 and Primary 2; HER2-on) and two recurrent tumor–derived cell lines (42929 and 48316, referred to as Recurrent 1 and Recurrent 2; HER2-off) (Fig. 1A-C). These lines were derived from an inducible HER2-driven genetically engineered mouse model (MMTV-rtTA;TetO-HER2/neu) in which doxycycline-dependent HER2 expression initiates primary tumor formation, and subsequent withdrawal of doxycycline induces complete tumor regression, leaving behind dormant residual cells that eventually give rise to spontaneous HER2-independent recurrences. Primary lines were isolated prior to oncogene downregulation, whereas recurrent lines were established from late-arising recurrent tumors (Fig. 1A). The library included sgRNAs targeting 10 pan-essential genes and 100 nontargeting sgRNAs. Cells were transduced with the sgRNA library at a multiplicity of infection (MOI) of 0.3 and selected in puromycin. Following selection, cells were collected at Day 0 or passaged for 7 or 14 population doublings (PD7 or PD14), and sgRNA abundance at each timepoint was measured using next-generation sequencing.

**Figure 1.**
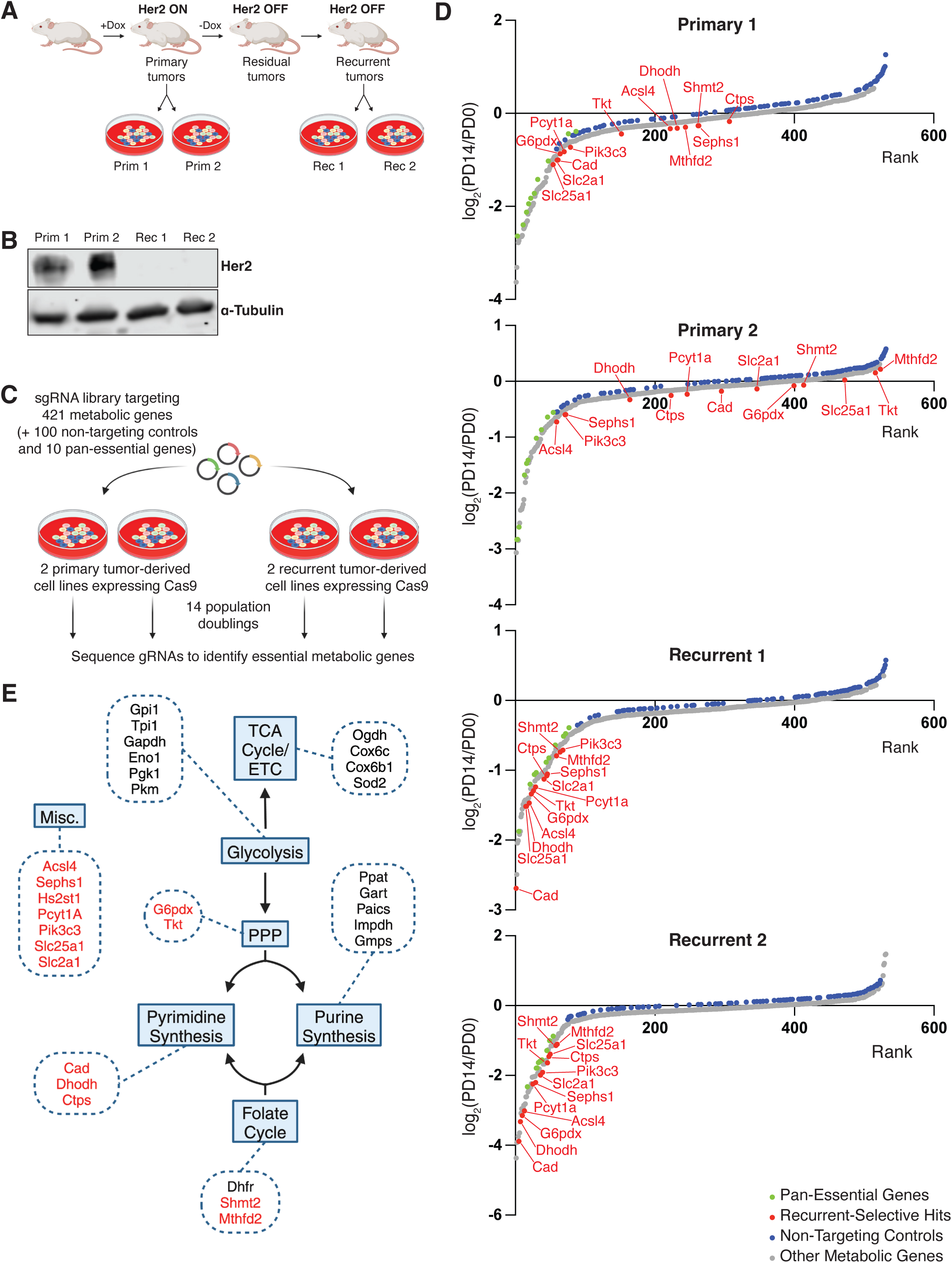
Identification of recurrent tumor-selective metabolic dependencies. **A.** Schematic of the inducible mouse model of HER2-driven primary mammary tumors and Her2-independent recurrent tumor. **B.** Western blot showing HER2 expression in cell lines derived from primary (P1 and P2) and recurrent (R1 and R2) tumors. **C.** Overview of the focused CRISPR screen used to identify essential metabolic genes in primary and recurrent tumor cells. **D.** Log fold-change (LFC) of gRNA abundance between Day 0 (PD0) and Population Doubling 14 (PD14) in two primary and two recurrent tumor-derived cell lines. LFC and FDR values were calculated using MAGeCK. **E.** Essential metabolic genes identified in the CRISPR screen, grouped by pathway. Common essential genes are shown in black; recurrent-selective essential genes are shown in red.

MAGeCK analysis identified 73 essential genes in Primary 1, 83 essential genes in Primary 2, 101 essential genes in Recurrent 1, and 84 essential genes in Recurrent 2 (false discovery rate (FDR) < 0.01; Table S2). sgRNAs targeting pan-essential genes were depleted in all cell lines, while nontargeting sgRNAs were not depleted, confirming the screen functioned as expected (Fig. 1D). Most essential genes identified were common to primary and recurrent tumor cells. For instance, genes in the glycolysis pathway, the citric acid cycle (TCA) and electron transport chain (ETC), and the *de novo* purine synthesis pathway, were identified as essential in both primary and recurrent tumor cells (Figure 1E).

To identify recurrent-selective essential genes, we considered genes that met the following criteria: FDR < 0.01 in both recurrent lines, an average log-fold change (LFC) difference ≥ log₂1.5 between recurrent and primary lines, and an absolute LFC ≤ 1 in both primary lines. Using these criteria, we identified 14 genes as recurrent-selective dependencies (Fig. 1D,E; Table 1). Recurrent-selective essential genes were primarily in three metabolic pathways: the *de novo* pyrimidine synthesis pathway (*Cad, Dhodh,* and *Ctps*), the pentose phosphate pathway (*G6pdx, Tkt*), and the folate cycle (*Shmt2, Mthfd2*). The pentose phosphate pathway generates ribose-5-phosphate, a precursor for pyrimidine synthesis, while the folate cycle supplies 5,10-methylene-THF, an essential cofactor for thymidylate synthase. Taken together, these findings suggest that recurrent breast cancer cells are broadly dependent on the metabolic network that supports *de novo* pyrimidine nucleotide production.

**Table 1.**
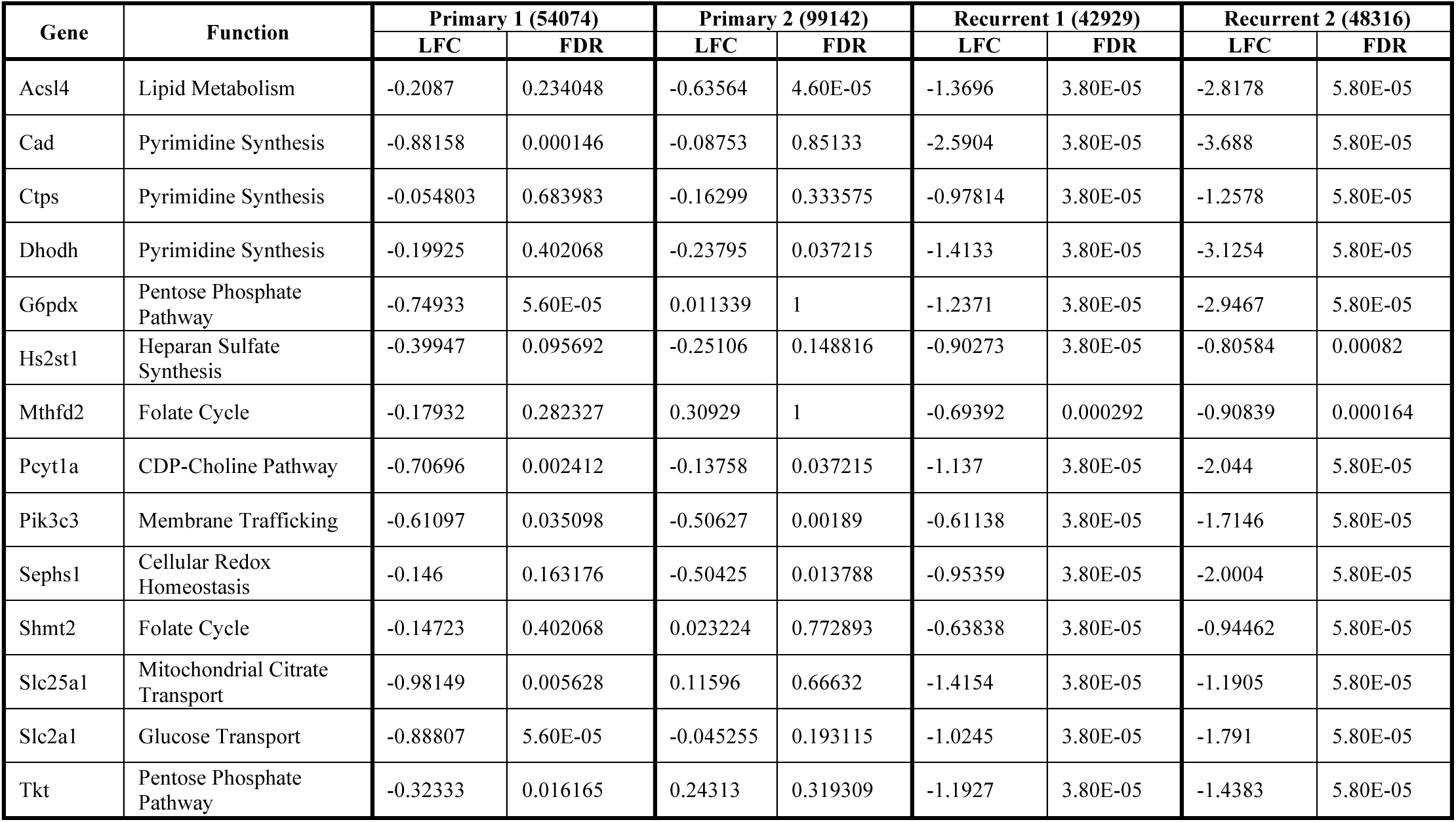
Recurrent-selective essential metabolic genes.

### DHODH activity is required for the growth of recurrent breast cancer cells

Among the top hits from our CRISPR screen was *Dhodh*, a rate-limiting enzyme in the *de novo* pyrimidine synthesis pathway that catalyzes the ubiquinone-mediated oxidation of dihydroorotate (DHO) to orotate (Fig. S1A). DHODH has been proposed as a therapeutic target in autoimmune and inflammatory disorders and cancer (9–12,18). Several small-molecule inhibitors of Dhodh, including brequinar, leflunomide, and BAY-2402234, are either FDA approved or in clinical development (9,19).

We first sought to validate the CRISPR screen result identifying *Dhodh* as a recurrence-selective vulnerability. To this end, we transduced primary and recurrent tumor cells with Cas9 and an sgRNA targeting *Dhodh*, resulting in approximately 75% reduction in DHODH expression (Fig. S1B). This depletion markedly impaired the growth of recurrent tumor cells but had minimal impact on the proliferation of primary tumor cells (Fig. 2A).

**Figure 2.**
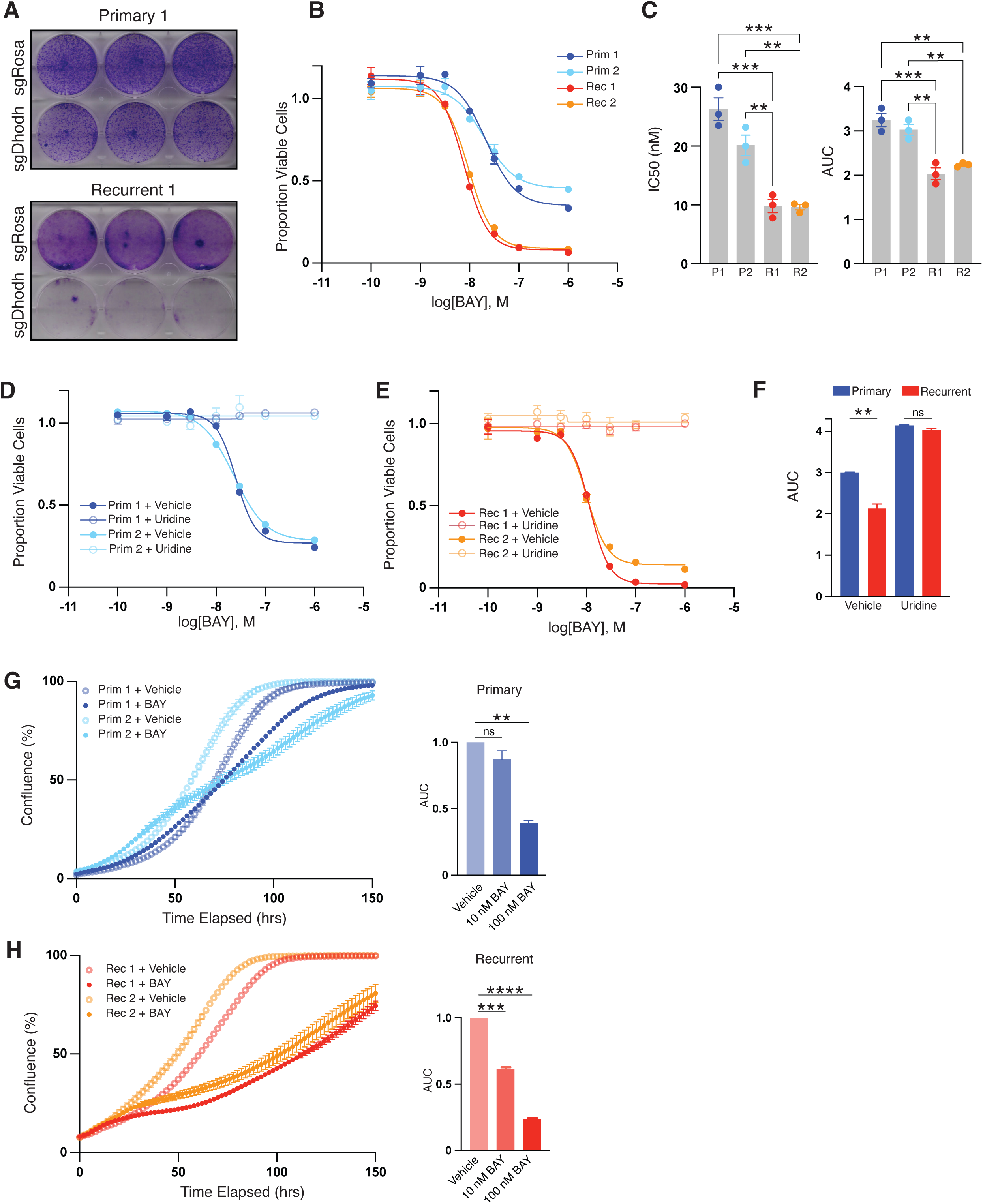
Dhodh activity is selectively required for the growth of recurrent tumor cells. **A.** Crystal violet staining of primary and recurrent tumor cells expressing either a control gRNA or a gRNA targeting Dhodh. **B.** Representative CellTiter-Glo (CTG) assay assessing viability of two independent primary and two independent recurrent tumor cell lines treated with BAY 2402234. **C.** Average IC₅₀ (left) and AUC (right) values for BAY 2402234 in primary and recurrent tumor cell lines, compiled from three independent CTG experiments. Statistical analysis was performed using ordinary one-way ANOVA; *** p ≤ 0.001, ** p ≤ 0.01. **D-F.** CTG assays demonstrating that exogenous uridine rescues BAY-induced loss of viability in both primary and recurrent tumor cells. **(F)** shows average AUC values of cells treated with BAY in the presence or absence of uridine. Statistical analysis was performed using ordinary one-way ANOVA, comparing primary versus recurrent cells; ** p ≤ 0.01. **G-H.** Cell growth curves obtained using the Incucyte Live-Cell Imaging System for two primary (G) and two recurrent (H) tumor cell lines treated with or without BAY 2402234. Average AUC values from a representative experiment are shown at right. To account for differences in baseline growth rates, AUC was calculated from 0 hours until the vehicle-treated condition reached 95% confluence. Statistical analysis was performed using ordinary one-way ANOVA, comparing primary versus recurrent cells; **** p ≤ 0.0001, *** p ≤ 0.001, ** p ≤ 0.01.

To further explore the dependence of recurrent tumor cells on DHODH activity, we tested the effects of pharmacologic inhibition of DHODH on primary and recurrent tumor cell growth. We used BAY-2402234 (BAY), which is a potent and selective inhibitor of DHODH (9). To confirm that BAY treatment inhibits pyrimidine synthesis, primary or recurrent tumor cells were treated with 10 nM BAY for 6 or 24 hours and metabolite levels in the pyrimidine synthesis pathway were measured using liquid chromatography-mass spectrometry (LC-MS). As expected, BAY treatment led to a robust accumulation of DHO and carbamoylaspartate, two metabolites upstream of DHODH, in both primary and recurrent cell lines (Fig. S1C). Consistent with these findings, we observed a significant reduction in downstream metabolites, including UMP, UDP, and UTP, following BAY treatment (Fig. S1D). Taken together, these changes in metabolite levels suggest that administration of 10 nM BAY leads to robust inhibition of DHODH in primary and recurrent tumor cells. Importantly, the extent of pathway inhibition was similar among all cell lines examined.

We next asked whether recurrent tumor cells are more sensitive to BAY treatment than primary tumor cells. Primary or recurrent tumor cells were treated with increasing doses of BAY for 3 days, and cell viability was determined using CellTiterGlo. Recurrent tumor cells were more sensitive to BAY treatment than primary tumor cells, with a 2-fold reduction in the GI50 dose (Primary 1: 26.3 nM; Primary 2: 20.13 nM; Recurrent 1: 9.84 nM; Recurrent 2: 9.64 nM; Fig. 2B-C). Even at the highest BAY concentration tested (1 µM), treatment only modestly reduced the growth of primary tumor cells but completely suppressed the growth of recurrent tumor cells (Fig. 2B). Consistent with this, the area-under-the-curve (AUC) values for BAY were significantly lower for recurrent tumor cells (Fig. 2C). Importantly, the growth-inhibitory effect of BAY was fully rescued by supplementation with exogenous uridine (Fig. 2D–F), confirming that BAY acts through inhibition of *de novo* pyrimidine synthesis.

We next wished to directly assess the effect of BAY on tumor cell growth using Incucyte live-cell imaging. Consistent with CellTiterGlo results, 10 nM BAY significantly slowed the growth of recurrent tumor cells but had only a modest effect on the growth of primary tumor cells (Fig. 2G-H). Taken together, these results indicate that DHODH inhibition selectively inhibits the growth of recurrent breast cancer cells and confirm the findings from the CRISPR screen that recurrent tumor cells are dependent upon *de novo* pyrimidine synthesis.

### DHODH inhibition induces DNA damage and S-phase arrest in both primary and recurrent tumor cells

We next investigated the cellular mechanisms underlying the selective dependence of recurrent tumor cells on DHODH. DHODH inhibition has previously been shown to induce DNA damage (12) and S-phase arrest (9), likely through depletion of pyrimidine dNTPs required for DNA synthesis. To determine whether DNA damage and cell cycle arrest were responsible for the growth defects following DHODH inhibition in recurrent tumor cells, we examined markers of DNA damage and cell cycle progression by western blot. We found that BAY treatment led to a dose-dependent increase in γH2AX levels and a decrease in phospho-histone H3 (pHH3) levels, confirming that DHODH inhibition induces DNA damage and cell cycle arrest (Fig. 3A). Interestingly, these responses were similar in primary and recurrent tumor cells (Fig. 3A).

**Figure 3.**
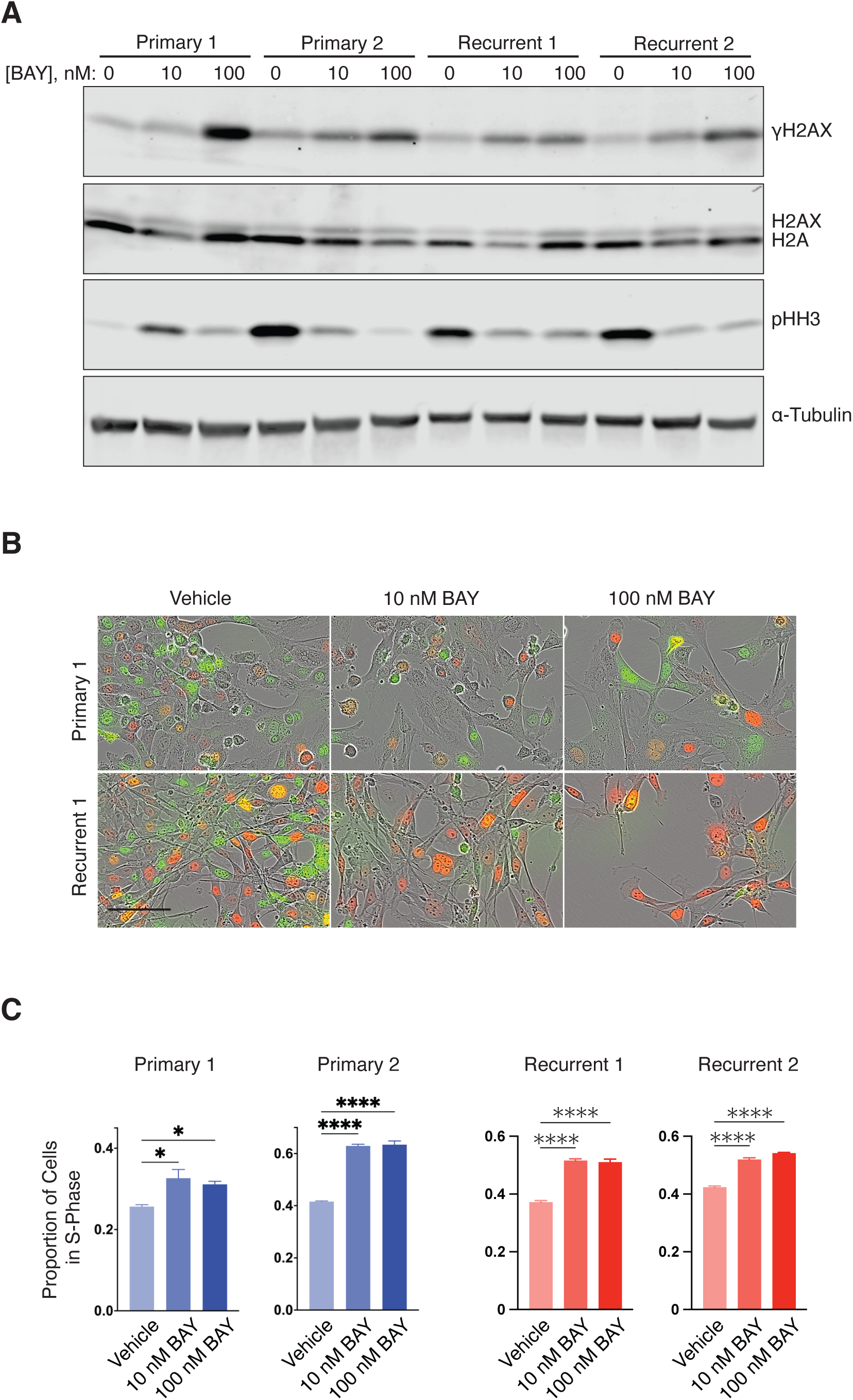
DHODH inhibition induces DNA damage and S-phase arrest in both primary and recurrent tumor cells. **A.** Western blot analysis of DNA damage and mitotic markers in two independent primary and two independent recurrent tumor cell lines treated with vehicle, 10 nM BAY 2402234, or 100 nM BAY 2402234 for 48 hours. Data are representative of three biologically independent experiments. **B.** Representative live-cell images of PIP-FUCCI–transduced cells from one primary and one recurrent tumor cell line. Cells with green nuclei are in G1 phase, those with red nuclei are in S phase, and those with overlapping green and red (yellow) are in G2/M. **C.** Quantification of the proportion of cells in S phase at 18 hours and 48 hours post-treatment, measured using the Incucyte system in two independent primary and two independent recurrent tumor cell lines. Statistical analysis was performed using ordinary one-way ANOVA, comparing each BAY treatment condition to its vehicle control; **** p ≤ 0.0001.

We next used the PIP-Fluorescent Ubiquitylation Cell Cycle Indicator (PIP-FUCCI) system to specifically delineate the cell cycle stage at which cells arrest following DHODH inhibition. Cells expressing the PIP-FUCCI construct exhibit green fluorescence when in the G1 phase of the cell cycle, red fluorescence in S-phase, and both green and red fluorescence in G2/M. We found that treatment with both 10 nM and 100 nM BAY for 48 hours induces accumulation of cells in S-phase (Fig. 3B, C). The increase in the proportion of cells in S-phase following DHODH inhibition was observed in both primary and recurrent cells (Fig. 3B, C).

Taken together, these data suggest that DHODH inhibition causes DNA damage, likely through dNTP depletion, and S-phase arrest. However, the similar responses observed in primary and recurrent tumor cells indicate that differential induction of DNA damage or cell cycle arrest does not account for their distinct sensitivities to DHODH inhibition, nor for their differential dependence on de novo pyrimidine synthesis.

### DHODH inhibition induces lipid peroxidation and ferroptotic cell death selectively in recurrent tumor cells

Recent work suggests that DHODH inhibition can induce ferroptosis (15,17), an iron-dependent form of cell death characterized by elevated lipid peroxidation(20), although the mechanism by which DHODH regulates ferroptosis remains controversial (16,21). We have shown that recurrent tumors exhibit heightened sensitivity to ferroptosis (13). We therefore tested whether DHODH inhibition induces ferroptosis, and whether this effect is selective for recurrent tumor cells.

Primary and recurrent tumor cells were treated with 10 nM BAY for 24 hours, and lipid peroxidation was quantified using the fluorescent probe C11-BODIPY 581/591 (22). BAY treatment caused robust lipid peroxidation in both recurrent tumor cell lines but not in primary tumor cells (Fig. 4A–B). Importantly, this lipid oxidation was iron-dependent, as it was partially suppressed by the iron chelator deferoxamine (DFO) (Fig. 4C–D). To directly assess cell death, we performed Incucyte live-cell imaging using Cytotox Red, a viability dye that detects the loss of membrane integrity characteristic of ferroptosis. BAY treatment (10 nM) increased cell death in both recurrent tumor cell lines but not in primary tumor cells (Fig. 4E), and DFO partially rescued this effect (Fig. 4F). Together, these results indicate that DHODH inhibition selectively induces lipid peroxidation and ferroptotic death in recurrent tumor cells.

**Figure 4.**
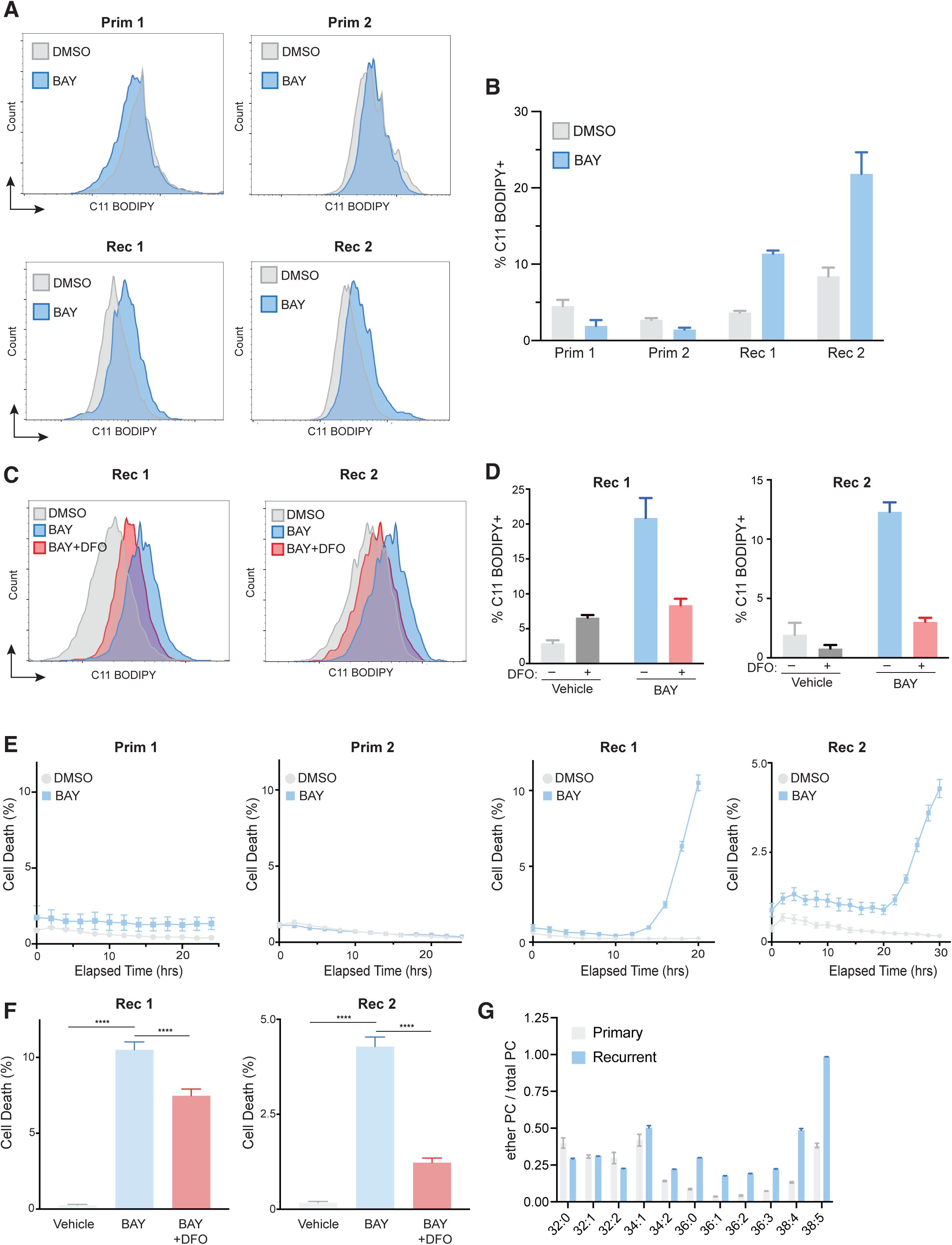
DHODH inhibition induces lipid peroxidation and ferroptosis selectively in recurrent tumor cells. **A.** C11 BODIPY staining of primary and recurrent tumor cells treated with 10 nM BAY 2402234 for 24 hours to assess lipid peroxidation. **B.** Quantification of C11 BODIPY-positive cells shown in (A). **C.** C11 BODIPY staining of recurrent tumor cells pre-treated with the iron chelator deferoxamine (DFO) for 2 hours prior to BAY treatment, demonstrating iron-dependent lipid peroxidation following DHODH inhibition. **D.** Quantification of lipid peroxidation data shown in (C). **E.** Cell death measured using the Incucyte Live-Cell Imaging System with Cytotox Red viability dye in two primary and two recurrent tumor cell lines treated with or without 10 nM BAY 2402234. **F.** Cell death quantified as in E, but with cells pre-treated with DFO prior to BAY exposure. The percentage of dead cells was assessed at the end of the imaging assay. **G.** Mass spectrometry-based quantification of ether phospholipids in primary and recurrent tumor cells.

Given that phospholipid composition strongly influences ferroptosis sensitivity (23), we next performed lipidomic profiling to determine whether distinct lipid compositions between primary and recurrent tumor cells underlie the ferroptosis susceptibility of recurrent tumors. Prior studies have demonstrated that peroxisome-derived ether phospholipids, particularly polyunsaturated ether-linked species, are potent drivers of ferroptosis by supplying highly peroxidizable lipid substrates (24,25). Consistent with this model, we found that recurrent tumor cells were enriched for polyunsaturated ether phospholipids, particularly phosphatidylcholine species (Fig. 4G). These findings suggest that the ether lipid–enriched membranes of recurrent tumor cells heighten their susceptibility to lipid peroxidation and thus sensitize them to ferroptosis following DHODH inhibition.

Both the Kennedy and CDP–DAG pathways generate phospholipids using CDP-activated intermediates (Fig. S2A). Alternatively, ether phospholipids are synthesized via a peroxisomal pathway (26). Inhibition of *de novo* pyrimidine synthesis has recently been reported to trigger a compensatory increase in peroxisome-derived ether lipids(27). Because recurrent tumor cells are more sensitive to ferroptosis and have elevated levels of ether phospholipids, we hypothesized that these cells suppress the Kennedy and CDP–DAG pathways and instead increase the synthesis of peroxisome-derived ether phospholipids.

To test this, we compared the expression of phospholipid synthesis genes – including Pcyt1a, Pcyt1b, Pcyt2, and Cds1 – in primary and recurrent tumor cells. Recurrent cells exhibited marked downregulation of Pcyt2 and Cds1, with a modest decrease in Pcyt1b (Fig. S2B–F), whereas expression of Ctps and Pcyt1a was unchanged. Consistent with this, analysis of previously published metabolomic data from primary and recurrent tumors (5) revealed that CDP-choline and CDP-ethanolamine levels were dramatically reduced in recurrent tumor cells (Fig. S2G–I).

Together, these results suggest that recurrent tumor cells downregulate canonical phospholipid synthesis via the Kennedy and CDP–DAG pathways, and exhibit a compensatory increase in ether phospholipid production, thereby predisposing them to ferroptosis upon DHODH inhibition.

### CRISPR screen identifies nucleotide salvage and lipid metabolism pathways as modifiers of sensitivity to DHODH inhibition in primary breast cancer cells

To understand the molecular mechanisms underlying the differential dependence of primary and recurrent tumor cells on *de novo* pyrimidine synthesis pathway we performed a sensitizer CRISPR screen in primary tumor cells. Primary tumor cell line 2 was transduced with the metabolism-focused sgRNA library, and treated with 10 nM BAY or vehicle for 14 population doublings (Fig. 5A). Importantly, this dose of BAY only modestly slows the growth of primary tumor cells (Fig. 5B and see Fig. 2G). In this screen, sgRNAs that are selectively depleted in the BAY-treated cells target genes whose knockdown sensitizes cells to pyrimidine synthesis inhibition – that is, they are synthetically lethal with DHODH inhibition. MAGeCK analysis identified 5 genes that were significantly depleted (FDR < 0.01) in BAY-treated cells as compared to vehicle-treated cells (Table S3). One of these 5 genes is *Dhodh*, confirming that total loss of DHODH activity – in this case achieved through low-dose BAY treatment (10 nM) together with CRISPR knockout – inhibits the growth of primary tumor cells. This is consistent with our findings that high-dose BAY (100 nM) effectively inhibits primary tumor cell growth. Three of the top 5 hits – *Tk1*, *Tyms*, and *Dhfr* – are involved in nucleotide synthesis. Of note, TK1 is an important enzyme in the pyrimidine salvage pathway, phosphorylating thymidine to form dTMP. The remaining hit, *Plpp1*, is involved in regulating signaling from bioactive lipids and may indirectly regulate lipid uptake (28).

**Figure 5.**
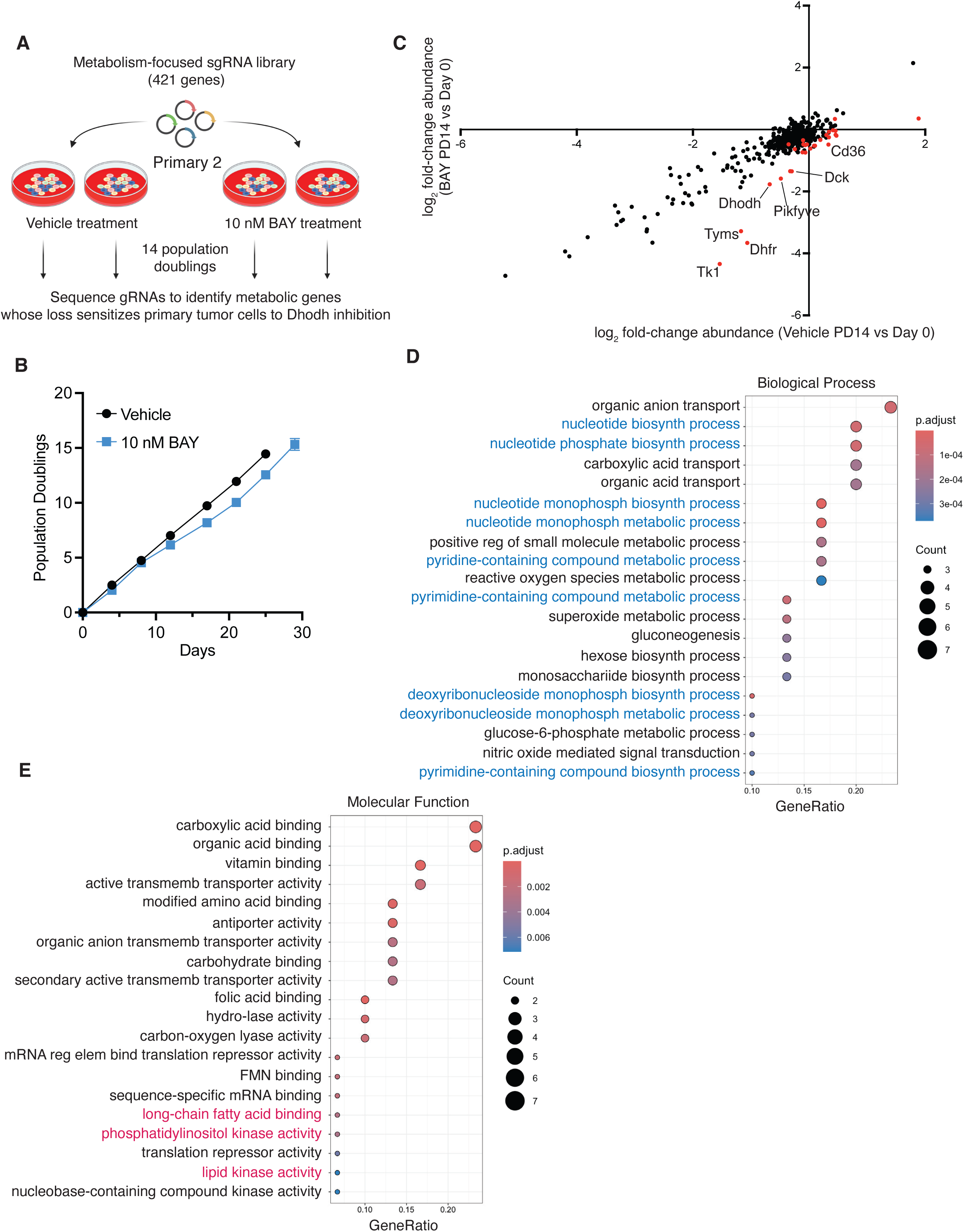
CRISPR screen to identify genes that sensitize primary tumor cells to Dhodh inhibition. **A.** Overview of the focused CRISPR screen used to identify metabolic genes whose knockout sensitizes primary tumor cells to Dhodh inhibition. **B.** Proliferation of primary tumor cells (Primary #1) following Dhodh inhibition. Population doublings were measured over time in cells treated with vehicle or 10 nM BAY 2402234, and cumulative population doublings were calculated at each passage. **C.** Scatter plot showing LFC of gRNAs at the endpoint (PD14) relative to Day 0 in vehicle-treated (x-axis) versus BAY-treated (y-axis) cells. gRNAs preferentially depleted in BAY-treated cells (red) correspond to genes whose knockout sensitizes primary tumor cells to Dhodh inhibition. **D-E.** Gene Ontology enrichment analysis of Biological Processes (D) and Molecular Functions (E) among BAY-sensitizer genes.

Using a less stringent cutoff (uncorrected p-value < 0.05), we identified 30 significantly depleted genes (Table S3). Gene Ontology enrichment analysis of these hits revealed genes involved in pyrimidine synthesis, including nucleotide salvage genes such as *Dck* (Fig. 5D). Interestingly, these hits were also enriched for genes involved in lipid metabolism (“phospholipid metabolic process”, p-adjusted = 0.0049; “lipid modification”, p-adjusted = 0.0091; “long-chain fatty acid binding”, p-adjusted = 0.0029; “phosphatidylinositol kinase activity”, p-adjusted = 0.0033; “lipid kinase activity”, p-adjusted = 0.0079; Fig. 5D-E and Table S4). These included the lipid translocase *Cd36*; the lipid kinase *Pikfyve*; *Ppard*, a transcription factor that regulates lipid metabolism genes; and *Hadh*, a mitochondrial enzyme involved in fatty acid β-oxidation (Table S4).

Taken together, these results indicate that inhibition of both nucleotide salvage and lipid metabolism sensitizes primary tumor cells to DHODH inhibition. These results suggest that differences in nucleotide salvage and lipid metabolism pathways between primary and recurrent tumor cells may mediate their differential dependence on *de novo* pyrimidine synthesis.

### Recurrent tumors are defective in nucleotide salvage

To explore the hypothesis that differences in nucleotide salvage between primary and recurrent tumor cells underlies their differential dependence on *de novo* pyrimidine synthesis, we first determined the proportion of pyrimidine nucleotides derived from *de novo* synthesis in each cell line. During *de novo* synthesis, glutamine contributes its amide nitrogen to carbamoyl-phosphate in the first reaction catalyzed by Cad. Following subsequent reactions catalyzed by CAD and DHODH, the glutamine-derived nitrogen gets incorporated at the N3 position of the pyrimidine ring. Labeling cells with ^15^N-amide glutamine allows for determination of the fraction of each nucleotide that is generated from *de novo* synthesis. We labeled primary or recurrent tumor cells with ^15^N-glutamine for 6, 24, or 48 hours and measured incorporation of ^15^N into UMP or UTP using LC/MS. There was a time-dependent increase of ^15^N into both UMP and UTP in all cell lines (Fig. 6A-B). After 48 hours of labeling, nearly 100% of UMP and UTP were derived from ^15^N -glutamine in all cell lines (Fig.6A-B), indicating that, at baseline, both primary and recurrent tumor cells generate nearly all of their UMP and UTP from *de novo* synthesis rather than salvage.

**Figure 6.**
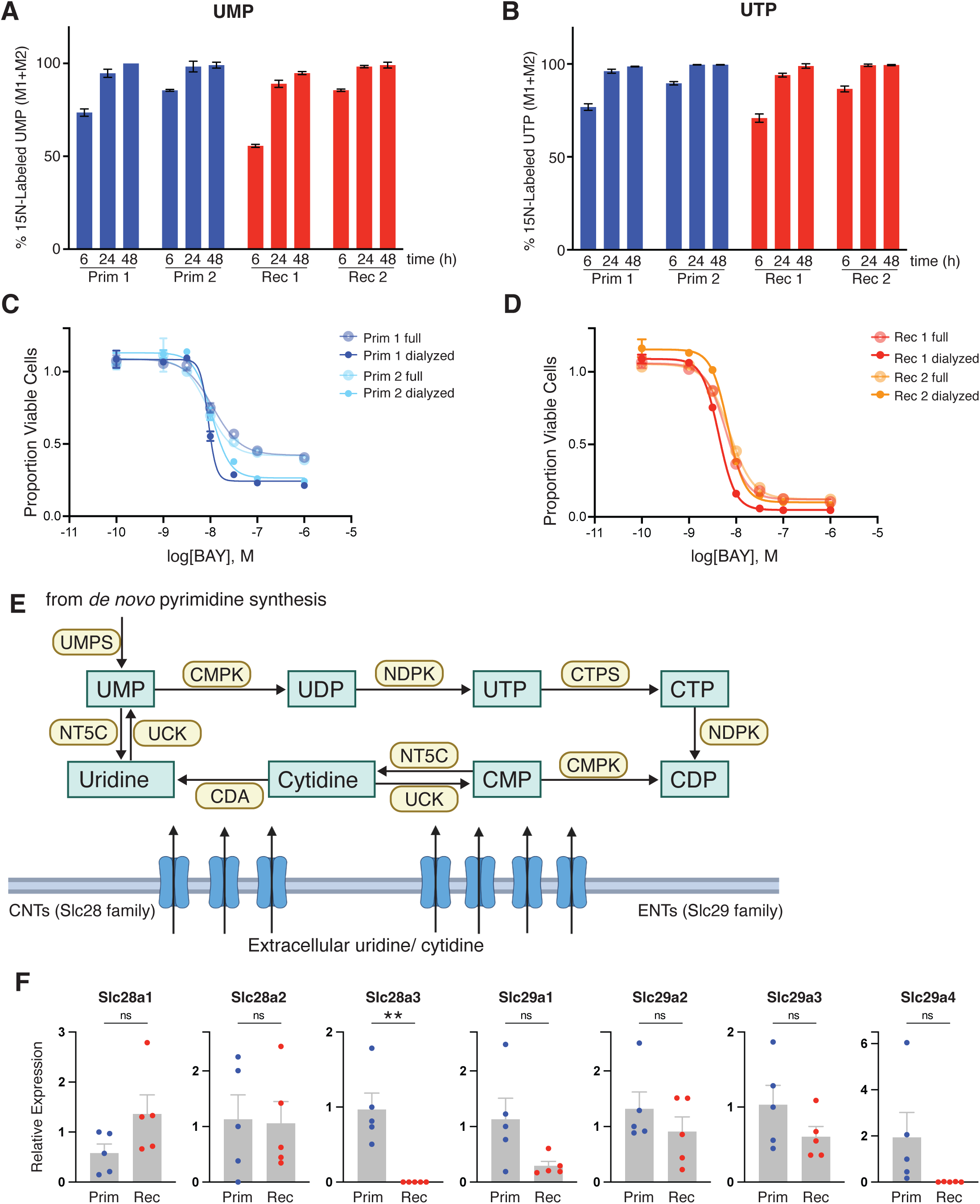
Recurrent tumors are defective in nucleotide salvage. **A-B.** Incorporation of ^15^N-glutamine into UMP (A) and UTP (B) in primary and recurrent tumor cells after 6, 24, and 48 hours of labeling. **C-D.** CellTiter-Glo viability assays of two primary (C) and two recurrent (D) tumor cell lines treated with increasing concentrations of BAY in normal serum or dialyzed serum. **E.** Schematic of the nucleotide salvage pathway. **F.** qPCR analysis of Slc28a and Slc29a gene expression in five independent primary tumor cell lines and five independent recurrent tumor cell lines. Statistics were performed using an unpaired t-test comparing primary versus recurrent cell lines. * *p* ≤ 0.05. **G.** qPCR analysis of Slc28a and Slc29a gene expression in five independent primary tumors and five independent recurrent tumors from MTB;TAN mice. Statistics were performed using an unpaired t-test comparing primary versus recurrent tumors. **** *p* ≤ 0.0001, ** *p* ≤ 0.01, * *p* ≤ 0.05.

While these data suggest that primary and recurrent tumor cells preferentially utilize *de novo* synthesis at baseline, we hypothesized that DHODH inhibition may induce cells to switch to nucleotide salvage pathway to compensate for loss of pyrimidine synthesis. To address this, we asked whether removing nucleotide precursors from cell culture media would affect sensitivity to DHODH inhibition. We cultured primary or recurrent tumor cells in media containing dialyzed calf serum, which removes low molecular weight metabolites including nucleosides and nucleobases. Culturing primary tumor cells in dialyzed media increased their sensitivity to DHODH inhibition (Fig. 6C). In contrast, dialyzed media had no effect on the response of recurrent tumor cells to DHODH inhibition (Fig. 6D). This suggests that primary tumor cells, but not recurrent tumor cells, are able to use nucleotide salvage to promote cell viability following inhibition of *de novo* pyrimidine synthesis.

To explore the mechanistic basis for differences between primary and recurrent tumor cells’ ability to perform nucleotide salvage, we measured the expression of key genes in the pyrimidine salvage pathway. Nucleosides and nucleobases are transported into cells through two families of solute carrier (SLC) transporters, SLC28 and SLC29 (Fig. 6E). We found that expression of several of these transporters, including *Slc28a3* and *Slc29a4*, was significantly reduced in recurrent tumor cells (Fig. 6F).

Taken together, these results suggest that primary cells can switch from *de novo* pyrimidine synthesis to salvage pathways following DHODH inhibition. In contrast, recurrent tumor cells are unable to perform nucleotide salvage, likely due to reduced expression of SLC28 and SLC29 family of transporters. This difference in the capacity to perform nucleotide salvage may underlie the increased dependence of recurrent tumor cells on *de novo* pyrimidine synthesis.

### Dependence on *de novo* pyrimidine synthesis in human breast cancer cells

We next asked whether differential dependence on *de novo* pyrimidine synthesis is also observed in human breast cancer cells. Using publicly available DepMap data, we examined breast cancer cell lines for dependency on *CAD* and *DHODH*, two key enzymes in the *de novo* pyrimidine synthesis pathway. Dependence scores for these genes were highly concordant across cell lines, indicating that each reflects overall pathway reliance (Fig. S3A). Based on these data, we selected three cell lines with low predicted pathway dependence (HCC1419, BT549, T47D) and three with high dependence (HCC1954, MDA-MB-231, ZR-75-1) for further testing. Cells were treated with increasing concentrations of BAY for three days, and viability was measured by CellTiter-Glo (Fig. S3B). Consistent with DepMap predictions, sensitivity to BAY correlated with pathway dependence: the low-dependence lines showed minimal response (AUC = 4.20, 4.10, and 3.91), whereas the high-dependence lines were inhibited in the low-nanomolar range (AUC = 3.31, 3.27, and 2.90). These results suggest that, as in our HER2-driven primary and recurrent tumor models, only a subset of breast cancer cells exhibit a strong requirement for *de novo* pyrimidine synthesis.

We next tested whether resistance to BAY in the less-dependent lines was mediated by nucleoside salvage from serum. Cells were cultured in dialyzed serum to remove exogenous nucleosides and retested for BAY sensitivity. Two resistant lines (T47D and BT549) became sensitized under dialyzed conditions (Fig. S3C). Among the sensitive lines, HCC1954 and MDA-MB-231 were further sensitized, while ZR-75-1 was unaffected. These findings suggest that the spectrum of responses seen in primary versus recurrent HER2-driven tumors is recapitulated, at least in part, in human breast cancer cell lines. T47D and BT549 cells resemble primary tumor cells – relatively resistant to BAY due in part to nucleotide salvage – whereas ZR-75-1 cells resemble recurrent tumor cells, displaying intrinsic sensitivity to BAY and limited ability to salvage extracellular nucleotides.

### Inhibition of HER2 signaling downregulates expression of nucleotide salvage and phospholipid synthesis genes

Our findings indicate that recurrent tumor cells exhibit reduced expression of several solute carriers and phospholipid synthesis genes. Because primary tumor cell growth is driven by HER2 signaling, which promotes broad metabolic activity, we hypothesized that the loss of these metabolic genes in recurrent tumor cells may result from diminished HER2 signaling. To test this, we examined whether acute inhibition of HER2 downregulates these genes in vivo. Doxycycline was administered to MMTV-rtTA;TetO-HER2/neu mice to induce primary HER2-driven tumors. When tumors reached ∼15 mm in diameter, one cohort of mice was euthanized to harvest primary tumors, while the remaining mice were removed from doxycycline to downregulate HER2 and initiate regression (Fig. 7A). A second cohort was euthanized two days after dox withdrawal to assess the immediate effects of HER2 inhibition, and a third cohort was monitored for recurrent tumor formation and euthanized when recurrent tumors reached 10–15 mm. qPCR analysis confirmed that HER2 expression decreased within two days of dox withdrawal and was fully suppressed in recurrent tumors (Fig. 7B). Notably, *Slc28a3* and *Slc29a4* exhibited the same temporal pattern – an initial reduction following acute HER2 inhibition and a further decrease in recurrent tumors – suggesting that their expression is regulated by HER2 signaling in mammary tumors (Fig. 7C). Expression of phospholipid synthesis pathway genes (*Pcyt1b*, *Pcyt2*, *Cds1*) similarly declined shortly after HER2 withdrawal and remained low in recurrent tumors (Fig. 7C). Importantly, for both SLC genes and phospholipid genes, this pattern mirrors the downregulation observed in recurrent tumor cell lines (cf. Fig. 6F and S2C with Fig. 7B-C). Taken together, these results demonstrate that HER2 signaling maintains expression of key solute carrier and phospholipid synthesis genes in primary tumors, and that their reduced expression in recurrent tumors is a direct consequence of loss of HER2 signaling.

**Figure 7.**
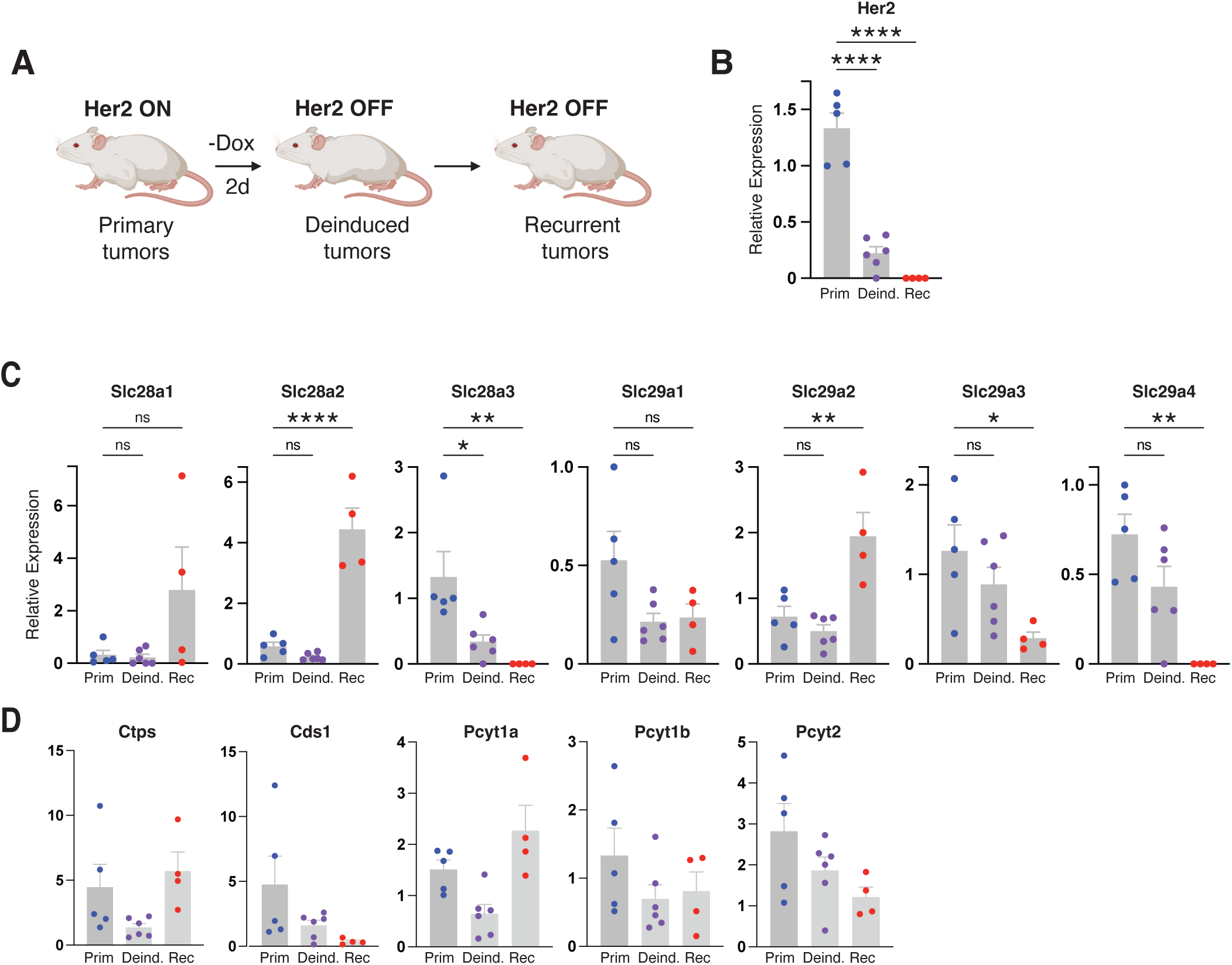
Inhibition of Her2 signaling downregulates expression of nucleotide salvage and phospholipid synthesis genes. **A.** Schematic of the experimental workflow used to generate primary HER2-driven tumors, tumors following acute HER2 downregulation (“deinduced” tumors), and recurrent tumors. **B–D.** qPCR analysis of Her2 (B), Slc genes (C), and phospholipid synthesis genes (D) in primary, deinduced, and recurrent tumors. Statistical analyses were performed using an ordinary one-way ANOVA comparing deinduced and recurrent tumor chunks to primary tumors (****p ≤ 0.0001, **p ≤ 0.01, *p ≤ 0.05).

## Discussion

Our study identifies *de novo* pyrimidine biosynthesis as a selective and targetable metabolic vulnerability in recurrent breast cancer, linking it to ferroptosis regulation and nucleotide salvage deficiency. Using paired primary and recurrent HER2-driven breast cancer models, we found that recurrent tumor cells exhibit marked dependence on DHODH and other core enzymes in the pyrimidine biosynthetic pathway. Pharmacologic inhibition of DHODH with BAY-2402234 suppressed the growth of recurrent tumor cells, whereas primary tumor cells were relatively resistant, despite equivalent blockade of pyrimidine synthesis. These findings reveal that recurrent tumor cells rely on pyrimidine metabolism in ways that extend beyond simple nucleotide provision.

Mechanistically, our results connect DHODH activity to ferroptosis resistance, consistent with recent work establishing DHODH as a mitochondrial anti-ferroptotic enzyme. Mao et al. (15) demonstrated that DHODH, localized to the inner mitochondrial membrane, reduces ubiquinone to ubiquinol and thereby suppresses lipid peroxidation in parallel to GPX4, though subsequent work suggested that the effects of Dhodh inhibition on ferroptosis are more complex (16). We find that recurrent breast tumor cells occupy a ferroptosis-prone metabolic state characterized by an enrichment of polyunsaturated ether phospholipids, a lipid class previously linked to ferroptosis sensitivity (24,25). In this context, DHODH inhibition tips the redox balance toward catastrophic lipid peroxidation and cell death. These results are reminiscent of recent findings that DHODH inhibition decreases CDP-choline and phosphatidylcholine synthesis, sensitizing tumor cells to CD8⁺ T-cell-mediated ferroptosis and enhancing responses to PD-1 blockade (17). Collectively, our data and recent studies converge on a model in which DHODH maintains ferroptosis resistance by sustaining CDP-driven phosphatidylcholine biosynthesis and buffering mitochondrial redox stress. Loss of DHODH activity lowers CDP levels, diminishes phosphatidylcholine synthesis, promotes compensatory ether-lipid accumulation, and renders tumor cells vulnerable to unrestricted lipid peroxidation—thereby creating a metabolic state that is permissive for ferroptosis and potentially exploitable therapeutically.

Our results also highlight a distinct loss of nucleotide salvage capacity in recurrent tumors, further amplifying their dependence on *de novo* synthesis. Primary tumor cells compensate for DHODH inhibition by salvaging extracellular nucleosides, whereas recurrent cells fail to do so, likely due to reduced expression of SLC28 and SLC29 nucleoside transporters. This mirrors findings in diffuse midline glioma (11), where elevated uridine degradation rendered cells dependent on *de novo* pyrimidine synthesis, and in IDH1-mutant glioma (12), where the oncogenic metabolic background created a synthetic-lethal requirement for this pathway. Similarly, small-cell lung cancers exhibit an intrinsic addiction to DHODH (10). Together, these studies support a broader model in which specific oncogenic or adaptive states curtail nucleotide salvage and channel survival through DHODH-driven pyrimidine metabolism.

From a translational perspective, DHODH inhibitors such as BAY-2402234 and brequinar have demonstrated preclinical efficacy in brain and lung cancer models and show favorable brain penetration and pharmacodynamics. Our data extend this therapeutic rationale to recurrent, therapy-resistant breast cancer, a disease setting in which metabolic rewiring may create selective liabilities not present in primary tumors.

Furthermore, our sensitizer CRISPR screen indicates that combining DHODH inhibition with perturbations in nucleotide salvage or lipid metabolism could potentiate ferroptosis and achieve greater selectivity for recurrent disease.

Finally, our findings reinforce a broader conceptual framework: recurrent and residual tumor cells inhabit a distinct metabolic landscape defined by altered redox state, impaired salvage capacity, and an increased need to suppress ferroptosis. These adaptations, while supporting survival under therapeutic stress, generate new dependencies that can be exploited therapeutically. Future studies should determine how these pathways operate in vivo, how they influence immune composition and response, and whether similar DHODH-ferroptosis couplings are conserved across metastatic or therapy-resistant cancers.

In summary, our work establishes *de novo* pyrimidine biosynthesis as a central metabolic vulnerability in recurrent breast cancer, mechanistically linking DHODH-driven nucleotide metabolism to ferroptosis suppression. We propose that targeting DHODH could represent a unifying therapeutic strategy for eliminating metabolically adapted, therapy-resistant tumor populations before they seed relapse.

## Materials and Methods

### Cell lines and cell culture

Primary and recurrent mammary tumor cell lines were generated as described previously(5,29,30) from bitransgenic MMTV-rtTA;TetO-Her/neu (MTB;TAN) mice on an FVB/N background. All mouse cell lines were cultured in high glucose Dubecco’s modified Eagle’s medium (DMEM) (Gibco #11965092) supplemented with 10% iron-supplemented calf serum (Cytiva #SH30072.03), 1% L-glutamine (Gibco #25030-081), 1% Pen Strep (Gibco $15140-122), 10 ng ml^-1^ mouse EGF (Sigma #E4127), and 5 µg ml^-1^ bovine insulin (GeminiBio # 700-112P) at 37°C with 5% CO_2_. Cell culture medium for primary cells was also supplemented with 2 µg ml^-1^ doxycycline (RPI #D43020) to maintain HER2/neu expression, 1 µM progesterone (Sigma #P7556), and 1 µg ml^-1^ hydrocortisone (Sigma #H0396). For live-cell fluorescent imaging experiments in Fig. 4c and supplementary fig. 1c, FluoroBrite DMEM (Gibco #A1896701) was used. Human cell lines were obtained from American Type Culture Collection (ATCC) and subcultured according to ATCC protocols.

Cell viability assays were performed using CellTiter-Glo (Promega #G7571) according to the manufacturer’s instructions. 1 x 10^3^ cells were plated per well on an opaque 96-well plate. The next day, the medium was replaced with the indicated treatments (see Table S5 for drug supplier and catalog numbers). CellTiter-Glo reagent was added and luminescence values were read using a BioTek Synergy HTX plate reader 3 days after treatment.

### Incucyte live-cell imaging

For IncuCyte-based assays, cells were plated in six-well plates at densities ranging from 2.5×10^4^ to 1×10^5^ cells per well, yielding an initial confluence of approximately 30–70%. After allowing cells to adhere for 24 hours, the medium was replaced with complete growth medium containing either DMSO (Sigma, D2650) or BAY 2402234 (Selleck, S9947). For cell-death assays, 250 nM Incucyte® Cytotox Red Reagent (Sartorius, 4632) was added at the time of treatment.

Plates were immediately transferred to an IncuCyte S3 or SX5 Live-Cell Analysis System (Sartorius). For proliferation assays, phase-contrast images were acquired every 3 hours using a 10× objective, and percent confluence was quantified using IncuCyte analysis software. For cell-death assays, images were acquired every 2 hours using appropriate phase-contrast and red fluorescence channels. Nine technical replicates (images) per well were collected at each time point.

Image analysis was performed in IncuCyte 2023A (Sartorius). Red fluorescent objects (dead cells) were quantified using threshold (RCU) and area parameters optimized for each cell line. Live-cell counts were quantified by phase-contrast segmentation using optimized settings for segmentation adjustment, detection sensitivity, contrast, morphology, and area. Data were exported as .csv files for downstream analysis.

Cell-death percentage was calculated for each image as:

Cell death (%)=(Red count) / (Red count + Live cell count) × 100.

Values were averaged across replicates and plotted as a function of time after treatment using Prism (GraphPad).

### CRISPR screening

The metabolism-focused sgRNA library was prepared essentially as described previously(5). Briefly, a custom CRISPR library targeting 421 mouse genes and 10 pan-essential genes with four sgRNAs each and 100 non-targeting sgRNAs (see Supplemental Table 1) was designed using sequences from the Brie CRISPR knockout library(31). Pooled oligos (Twist Biosciences) were PCR amplified and cloned into lentiguide-puro by Gibson Assembly (Addgene #52963), assembled sgRNA vectors were transformed into competent cells, and library DNA was isolated as described(5). Lentivirus was prepared by co-transfecting HEK293T cells with lentivector, psPAX2 and pMD2.g and harvesting supernatant 48 hours post-transfection.

Two independent Cas9-expressing primary tumor cell lines and two independent Cas9-expressing recurrent tumor cell lines were transduced with library lentivirus and after 2 days, cells were treated with puromycin (Sigma #P8833) to select for transduced cells. After selection, 3 x 10^6^ cells were plated onto 2 15cm plates per replicate. Leftover cells were collected for “PD0” samples. Cell pellets were then collected after 14 population doublings. Cell pellets were stored at -80°C degrees before gDNA was extracted using the DNeasy Blood & Tissue Kit (Qiagen #69504). A library was prepared for sequencing, sequenced, and analyzed as described previously(5).

### Metabolite extraction and LC-MS analysis

For metabolite extraction, 1×10^5^ cells were plated per well in six-well plates and allowed to adhere overnight. The following day, vehicle- and 24-hour treatment groups were washed twice with PBS, after which the medium was replaced with complete growth medium containing either DMSO (Sigma, D2650) or BAY 2402234 (Selleck, S9947). Eighteen hours after the initial treatment, cells designated for the 6-hour treatment condition were washed twice with PBS and the medium was replaced with BAY 2402234-containing medium.

After the appropriate treatment duration, wells were washed once with ice-cold blood bank saline (Fisher, 23062125), and metabolites were extracted by adding 800 µL of ice-cold 80% HPLC-grade methanol (Sigma, 34860) prepared in HPLC-grade water (Sigma, 270733). Cells were scraped thoroughly, and 600 µL of extract was collected and clarified by centrifugation at 21,300 × g for 10 minutes at 4°C. A total of 400 µL of the supernatant was transferred to fresh tubes and dried in a CentriVap concentrator. Dried extracts were stored at −80°C until LC–MS analysis.

### Western blots, RT-qPCR, and virus production

Western blotting was performed as described previously(32). Antibodies and working concentrations are listed in Table S5. Membranes were imaged using the Odyssey Infrared Imaging System (LI-COR).

Total RNA was isolated using the RNeasy Mini Kit (Qiagen, 74104), and cDNA was synthesized using the ImProm-II Reverse Transcription System (Promega, A3800). TaqMan probes used for quantitative PCR are listed in Table S5. qPCR was performed on a ViiA 7 Real-Time PCR System (Applied Biosystems), and expression values were normalized to Tbp.

Lentivirus production was carried out in HEK293T cells by co-transfection of the lentiviral expression construct (Table S5) with psPAX2 and pMD2.G packaging plasmids (Addgene 12260 and 12259). Viral supernatants were collected at 48 h and 72 h post-transfection, clarified by filtration, and used to transduce target cells in the presence of 4–8 µg/mL polybrene (Sigma, 107689).

### Cell cycle analysis by PIP-FUCCI

Two independent primary mammary tumor cell lines and two independent recurrent mammary tumor cell lines were transduced with pLenti-PGK-Neo-PIP-FUCCI (Addgene, 118616). Forty-eight hours after transduction, cells were selected with G418 disulfate salt (Sigma, A1720) until non-transduced control cells were fully eliminated. Following selection, 2.5×10^4^ to 5×10^4^ cells were plated per well in six-well plates. The following day, the medium was replaced with complete growth medium containing either DMSO (Sigma, D2650) or BAY 2402234 (Selleck, S9947). Live-cell imaging was performed every 2 hours using a 20× objective with 500-ms exposure for the green channel and 400-ms exposure for the red channel. FUCCI green-, red-, and double-positive objects were quantified using Incucyte software.

### Flow cytometry

A total of 1 × 10^!^ adherent cells were plated per well in six-well plates and allowed to adhere for 24 hours. Cells were then treated with 10 nM BAY 2402234 or an equivalent volume of DMSO for 24 hours. For deferoxamine (DFO) conditions, cells were treated with 10 nM BAY 2402234 in combination with 100 µM DFO.

To assess lipid peroxidation, cells were incubated for 1 hour with 10 µM BODIPY™ 581/591 C11 (ThermoFisher, D3861); one well was left unstained as a negative control. After staining, cells were trypsinized, resuspended in HBSS, and passed through 70-µm cell strainers into flow cytometry tubes. Samples were analyzed immediately on a BD FACSymphony™ A3 cell analyzer.

Cells were first gated by forward and side scatter (FSC/SSC) to exclude debris, followed by FSC-H × FSC-A gating to remove doublets. To quantify lipid peroxidation, a gate capturing approximately 2.5% of the highest FITC-positive population in DMSO-treated cells was established using FlowJo software and applied uniformly across all samples. Relative BODIPY oxidation was compared across treatment groups. Data were graphed using Prism, and statistical significance was assessed using two-way ANOVA.

### Statistics and reproducibility

Plots and statistical analyses were generated using Prism 10 software (GraphPad). For CRISPR screening experiment, false discovery rate values were generated using MaGeCK (33) to determine statistical significance for depleted genes.

## ACKNOWLEDGEMENTS

We thank members of the Alvarez Lab for critical reading of the manuscript. This work was funded by the National Cancer Institute under award R01CA285322 and R01CA292658 (to J.V.A.), and by startup funds from the Fred Hutchinson Cancer Center (to J.V.A.). This research was supported by the following shared resources of the Fred Hutch/University of Washington/Seattle Children’s Cancer Consortium (P30 CA015704): Genomics & Bioinformatics Shared Resource, RRID:SCR_022606; Cellular Imaging Shared Resource, RRID:SCR_022609; Comparative Medicine Shared Resource, RRID:SCR_022610; and Flow Cytometry Shared Resource, RRID:SCR_022613.

## Supplemental Figure Legends

**Figure S1.**
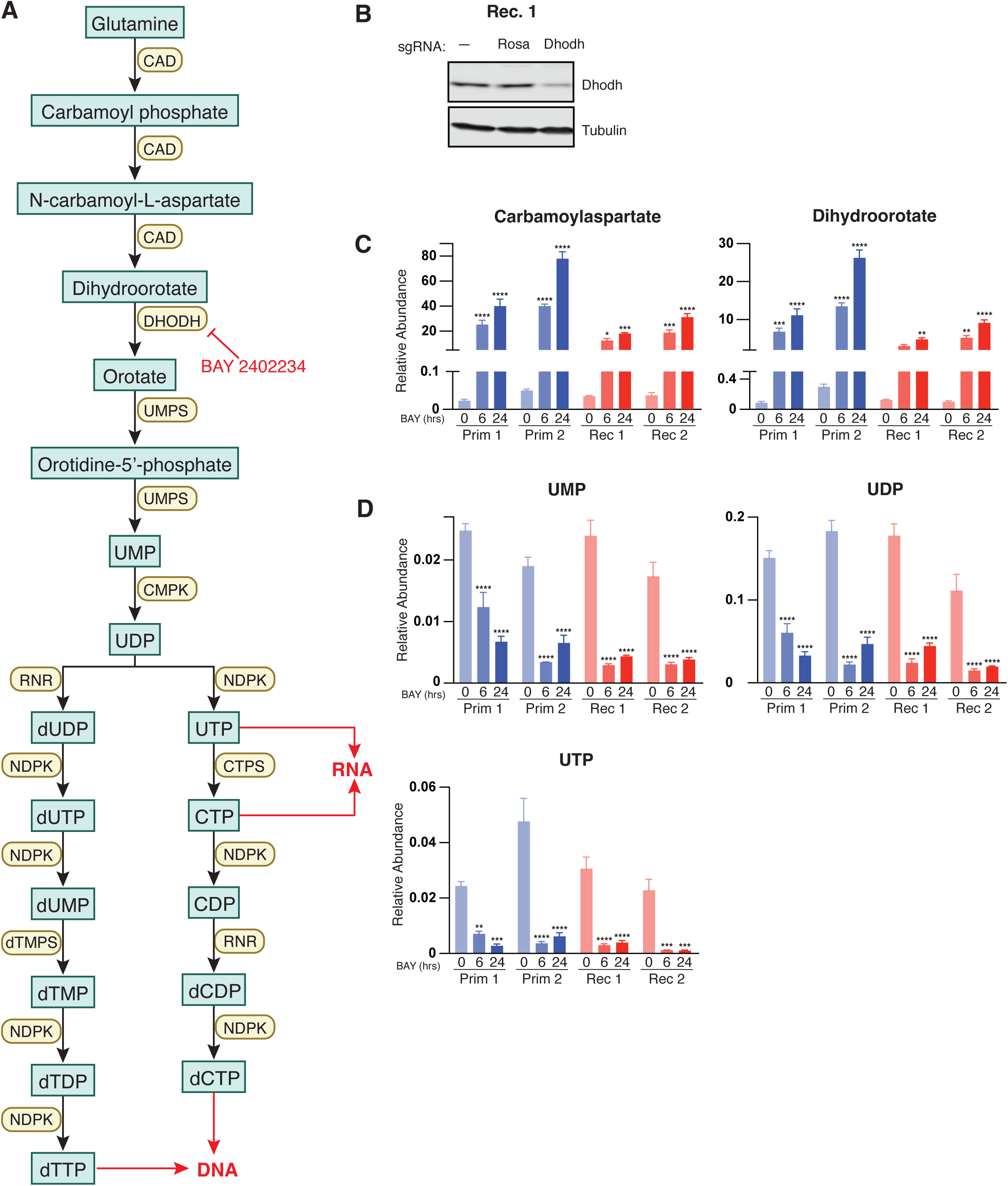
BAY 2402234 inhibits the *de novo* pyrimidine synthesis pathway in primary and recurrent tumor cells. **A.** Schematic of the *de novo* pyrimidine synthesis pathway. **B.** Western blot confirming loss of Dhodh protein following CRISPR-mediated knockout. **C-D.** Mass spectrometry-based quantification of metabolites upstream **(C)** or downstream **(D)** of Dhodh in the *de novo* pyrimidine synthesis pathway primary and recurrent tumor cells treated with 10 nM BAY for 6 or 24 hours. Statistical significance was assessed by ordinary one-way ANOVA, comparing each BAY-treated group (6 hr, 24 hr) to its matched 0 hr control. **** *p* ≤ 0.0001, *** *p* ≤ 0.001, ** *p* ≤ 0.01, * *p* ≤ 0.05.

**Figure S2.**
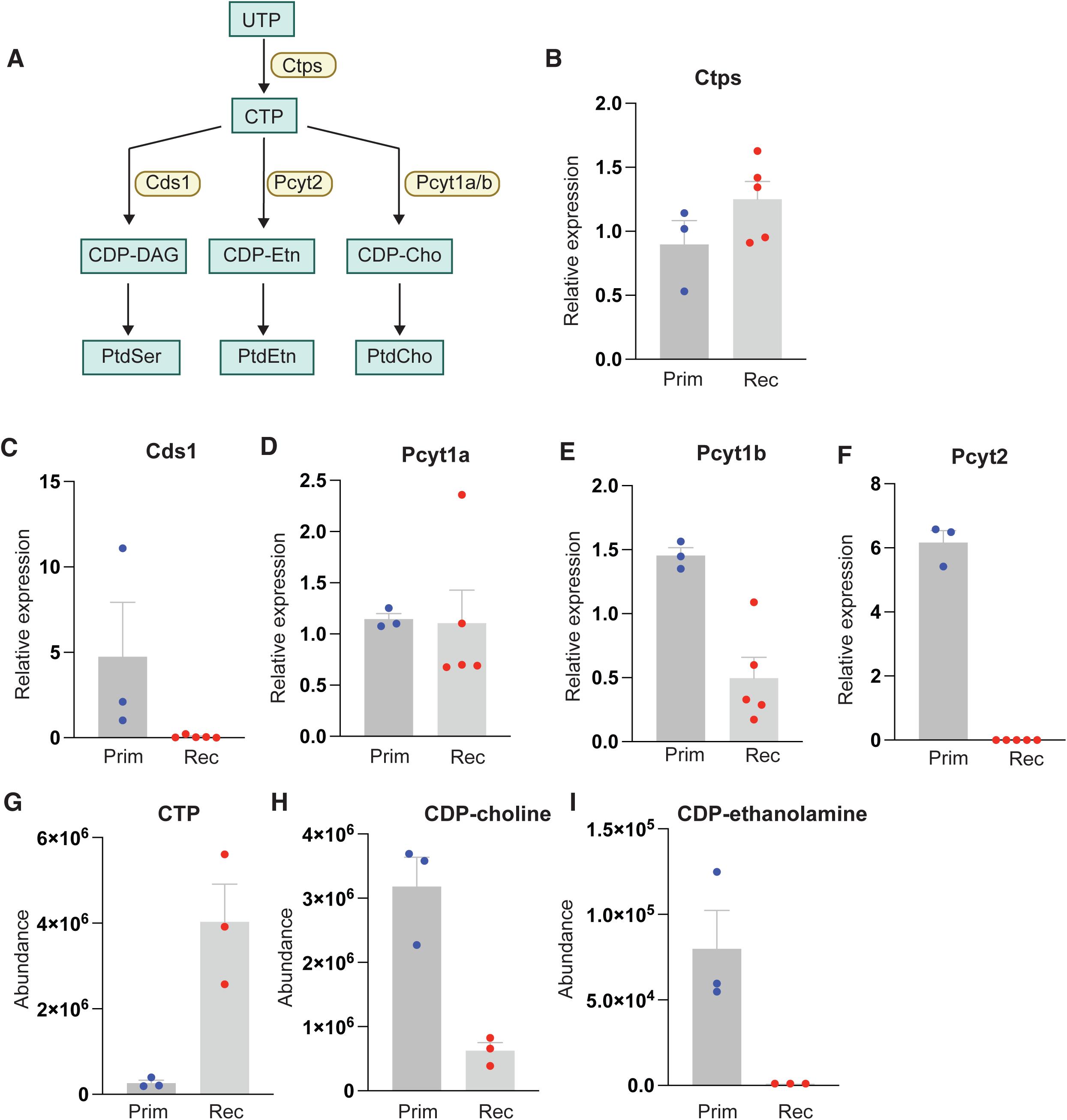
Recurrent tumors downregulate the phospholipid synthesis pathway. **A.** Schematic of the phospholipid synthesis pathway. **B-F.** qPCR analysis of the indicated phospholipid genes in three independent primary tumor cell lines and five independent recurrent tumor cell lines. **G-I.** Mass spectrometry-based quantification of the indicated phospholipid pathway metabolites in one primary and one recurrent tumor cell lines (three technical replicates per condition).

**Figure S3.**
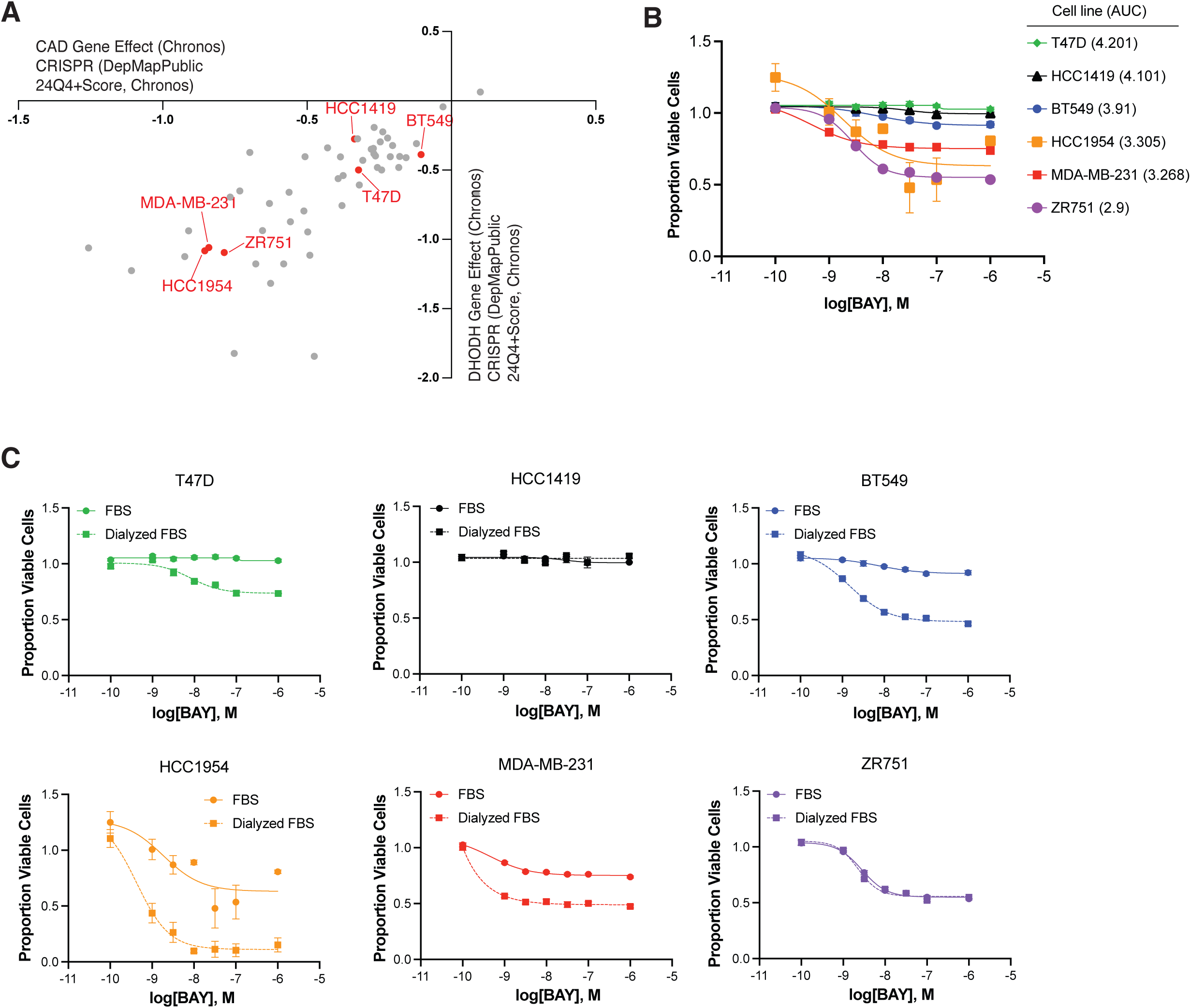
Dependence on de novo pyrimidine synthesis in human breast cancer cell lines. **A.** DepMap co-dependency analysis showing coordinated essentiality of CAD and DHODH, two key enzymes in the de novo pyrimidine synthesis pathway, across human breast cancer cell lines. Selected cell lines categorized as having low predicted dependence (HCC1419, BT549, T47D) or high predicted dependence (HCC1954, MDA-MB-231, ZR751) are highlighted in red. **B-C.** CellTiter-Glo viability assays of the selected human breast cancer cell lines treated with increasing concentrations of BAY 2402234 in full serum (FBS) or in dialyzed serum (dialyzed FBS). In B., the area under the curve (AUC) for each line is shown in the legend.

