## Supplemental Table 1 for "Recurrent Breast Cancer Cells Depend on *De novo* Pyrimidine Biosynthesis to Suppress Ferroptosis"

Table S1. Metabolism-focused gRNA library.

| **CRISPR Library** | | |
| --- | --- | --- |
| **Primer Name** | **Sequence** | **Gene Target** |
| control_0 | TTAGTTCGACCACTCCTGAT | non-targeting |
| control_1 | GTCCTAGATCCTATCGGGAG | non-targeting |
| control_2 | AGGACTATCCGCGGGATTAG | non-targeting |
| control_3 | GTAGCTGACCGCTTAACTAA | non-targeting |
| control_4 | CTCGGACGGCATACGACAAT | non-targeting |
| control_5 | ACGTGTAAGGCGAACGCCTT | non-targeting |
| control_6 | GGCGCTACTGTAATGACGGT | non-targeting |
| control_7 | ACTTTACATCATGTCGTCGT | non-targeting |
| control_8 | GGCACCGTTCGGAAACCGAC | non-targeting |
| control_9 | CATCGTAACACACGTACGAG | non-targeting |
| control_10 | GCCGCTCTTGATAACGACGC | non-targeting |
| control_11 | AAGCACAAGAACGGTCCGCC | non-targeting |
| control_12 | GTAGACATATTGCGTAATCG | non-targeting |
| control_13 | ATAGATGTCTACGCGCCGTT | non-targeting |
| control_14 | GTGATGGCCACGTCCGAACC | non-targeting |
| control_15 | AACGGTAGCGTACCCGTGAA | non-targeting |
| control_16 | TGCGGCAATGTTAACCCTTA | non-targeting |
| control_17 | AGTTTACGTGGTCCGATGTC | non-targeting |
| control_18 | TAAGCGGCGTCATGTCGCCC | non-targeting |
| control_19 | CATACTGTTATCGACCCGCA | non-targeting |
| control_20 | CAAGGGGTGGTCATTGCGAC | non-targeting |
| control_21 | ATTCGGACCGTTATCTCACC | non-targeting |
| control_22 | TGCAGTAGTCGCGGATAGGC | non-targeting |
| control_23 | CTACGAGATTCGGCTAAGCT | non-targeting |
| control_24 | GTGCCTGATAGTGTGAAGCG | non-targeting |
| control_25 | ATGACACTTACGGTACTCGT | non-targeting |
| control_26 | CGAGGATGTACATACGTAAA | non-targeting |
| control_27 | ACTGCGCGTATAGGACGCAA | non-targeting |
| control_28 | GGGTGCACACGCCGGCCTAT | non-targeting |
| control_29 | CAACGATCAGGCGTGTTATC | non-targeting |
| control_30 | CTACGAGGGCCGCGAGCGGT | non-targeting |
| control_31 | TAACACGCACTCACGTCCGG | non-targeting |
| control_32 | GTGTCTTTCGGTCTTACGAG | non-targeting |
| control_33 | CCGTTCTGACGACGCTAAAG | non-targeting |
| control_34 | AGCAGTTCGGGTAACGCCCA | non-targeting |
| control_35 | CGTCATGGCAAGATCGATAA | non-targeting |
| control_36 | ATTGTTCGACCGTCTACGGG | non-targeting |
| control_37 | CCGACTGCCGAGCTAGGCGT | non-targeting |
| control_38 | GTGCGCGCGTATAGAAAAAA | non-targeting |
| control_39 | TCTTCTCGAGCGTGCGGGCC | non-targeting |
| control_40 | TTCAACGACGGAAGACGCGC | non-targeting |
| control_41 | TTTACGTATGCTGTGCGCGA | non-targeting |
| control_42 | TGATGGACGCGATACGTTTA | non-targeting |
| control_43 | GCCCTCGAGCTCACGATGAG | non-targeting |
| control_44 | TACATCGAACAACCAACCGA | non-targeting |
| control_45 | GAGCTAATCTGTAGACGTCG | non-targeting |
| control_46 | AGCGTATCTTATGTACCGCT | non-targeting |
| control_47 | TTGTCAACTTCGGCCAACGC | non-targeting |
| control_48 | TTGGCGGCTCGCTCCTGCGT | non-targeting |
| control_49 | TGATGCCGGTACCCGTAACT | non-targeting |
| control_50 | CCTCCTTCAGTTCGATCTGG | non-targeting |
| control_51 | GCCGCGCCATAATATGCCAT | non-targeting |
| control_52 | TGATAAACGATGCGAACTCG | non-targeting |
| control_53 | CTCGTGTCACTCCTCGGTTC | non-targeting |
| control_54 | GACCGACGGTATACCCTACT | non-targeting |
| control_55 | TATGAACCTCCGGATCGGTG | non-targeting |
| control_56 | AGTCATCCTCTATGCGCGTA | non-targeting |
| control_57 | AACCCCGGCTGTCATCGCCG | non-targeting |
| control_58 | TCCTATAATTGAGCGAACGG | non-targeting |
| control_59 | CCGGACTTGTTATACTTGAT | non-targeting |
| control_60 | CAACCTGCCTAGCGACCCGC | non-targeting |
| control_61 | ACGAGGAGAAGTCGAGTATT | non-targeting |
| control_62 | AAGGCCTTAACACGTCGACC | non-targeting |
| control_63 | GTACCATGATAACCGTACTA | non-targeting |
| control_64 | GCCGAGAGGCGTAAGCGCGA | non-targeting |
| control_65 | GATAGAACAACAGCGGTGCA | non-targeting |
| control_66 | TTCATCGCAGATCGATTTCG | non-targeting |
| control_67 | CTCGATGCCACGGAAGGCGG | non-targeting |
| control_68 | GGATAGCCCGGTTGGTGCGT | non-targeting |
| control_69 | TTCCAGTATACCGAATTCGC | non-targeting |
| control_70 | TTGGACGTACACTTTCGTTC | non-targeting |
| control_71 | GCGATCGCCGGTATAGCTTT | non-targeting |
| control_72 | GGAAACTCTTGCTCGACACG | non-targeting |
| control_73 | TAGGGGTGCGCATTAGACTA | non-targeting |
| control_74 | CCGTCGAGCAATCCCGCCAA | non-targeting |
| control_75 | CTAATTCCAATATCCGTCTG | non-targeting |
| control_76 | AATGAAGCACCGATTGCGGA | non-targeting |
| control_77 | GCCACGACCTGCGAATAATT | non-targeting |
| control_78 | ACGACGAGAGATACTTAGGC | non-targeting |
| control_79 | CGTCGCATGTTGTTGCTCAT | non-targeting |
| control_80 | AGCGCCATAGGACGCCAAAC | non-targeting |
| control_81 | TCACAACCCCCGACTATCGC | non-targeting |
| control_82 | GGGGTAGGCCTAATTACGGA | non-targeting |
| control_83 | GGACGTAGATTAGGGCGTAA | non-targeting |
| control_84 | CGACTCCGGGAATATCCGTT | non-targeting |
| control_85 | TCAGCAGAATTGGGCCCGTA | non-targeting |
| control_86 | TTAACTGGGGGGACCGGACG | non-targeting |
| control_87 | CATTACGTGTCGAGCTCCGG | non-targeting |
| control_88 | TCTAGGATACTCTTAACGGG | non-targeting |
| control_89 | CTTAAGTTCGCGACGGAATG | non-targeting |
| control_90 | GTAGCTTCATGTCCGGTCGG | non-targeting |
| control_91 | GGGGATGGCCTTACGTCGCG | non-targeting |
| control_92 | TAACAGCTTCCGCGTAATAT | non-targeting |
| control_93 | TCGTAAAGTCGCAGCGACGT | non-targeting |
| control_94 | CGGATCCGGTACGGTCAGTT | non-targeting |
| control_95 | ACCGGATGTGGGCGCCTCTC | non-targeting |
| control_96 | GATGCGGGTGGAAAACGTTA | non-targeting |
| control_97 | ATCGTAAATTACACGTACAG | non-targeting |
| control_98 | AACCAGCATTTGACCGCGCT | non-targeting |
| control_99 | CGAATCGGGAAGGCGCGTGT | non-targeting |
| Aanat_1 | CGGATCTCATCCAAGTAGAG | AANAT |
| Aanat_2 | CACCGCCAGTACGTGCAGGT | AANAT |
| Aanat_3 | AGGAACTCTGAGGTCCCAAG | AANAT |
| Aanat_4 | CTCAATCTCAAACGCACTGG | AANAT |
| Abca1_1 | ACATGTCATCAACATAACAG | ABCA1 |
| Abca1_2 | GTGGACCCGTACTCTCGCAG | ABCA1 |
| Abca1_3 | CAAGCTGTCAAGCAACACTG | ABCA1 |
| Abca1_4 | GGTATACACAGAGCCATTTG | ABCA1 |
| Abcc1_1 | GGTGAAGTACAGATAGAACG | ABCC1 |
| Abcc1_2 | GGCTAGGATGACTTGCAGAG | ABCC1 |
| Abcc1_3 | AGCTGTAGGCACATTCACGT | ABCC1 |
| Abcc1_4 | GAGCGGAGGTCGATCAAGAG | ABCC1 |
| Abcc4_1 | AAGCGGTGAAATCTTGCACG | ABCC4 |
| Abcc4_2 | TGGTACTTAGGAATTTACGC | ABCC4 |
| Abcc4_3 | CAGACCCTCGTTGAAAGACG | ABCC4 |
| Abcc4_4 | GGTTAAGTAACTCGGCCATG | ABCC4 |
| Abcc5_1 | GGAGCTGATGAACTTAAATG | ABCC5 |
| Abcc5_2 | AAATCCGAGAGGAGGAACGT | ABCC5 |
| Abcc5_3 | CATTAAAGAAAAGTCCCTAG | ABCC5 |
| Abcc5_4 | AAAGACAAGAGGGCTACCAG | ABCC5 |
| Abce1_1 | AGGTTGCCTATCCCTCGTCC | ABCE1 |
| Abce1_2 | AGTTGCCCTGTGGTTCGGAT | ABCE1 |
| Abce1_3 | TATTAGAGATCGAATCGTAA | ABCE1 |
| Abce1_4 | ATGTTCCAACAGAGAACCTG | ABCE1 |
| Abcg1_1 | CAAGGACAATGCGTATACAG | ABCG1 |
| Abcg1_2 | GACTATAAGAGAGACCTCGG | ABCG1 |
| Abcg1_3 | CAGGCCGATCCCAATGTGCG | ABCG1 |
| Abcg1_4 | TTTCCTACTCTGTACCCGAG | ABCG1 |
| Acaca_1 | GCCATTCATTATCACTACGT | ACACA |
| Acaca_2 | AATGCATGCGATCTATCCGT | ACACA |
| Acaca_3 | TTGATTCATAGGTACCGAAG | ACACA |
| Acaca_4 | AAGCCCTTCGAACATACACC | ACACA |
| Acacb_1 | AGAACAACGATATCGACACG | ACACB |
| Acacb_2 | CCAGTGTGCCGACCACATGG | ACACB |
| Acacb_3 | GTGGCCGTACATGTCGATGG | ACACB |
| Acacb_4 | TTCATTACGGAACATCTCGT | ACACB |
| Acadl_1 | AGGCGATCGAGCTTCACGGT | ACADL |
| Acadl_2 | AGAGCGTACTCCAATTGCAC | ACADL |
| Acadl_3 | AACATCGCAGAGAAACATGG | ACADL |
| Acadl_4 | CCGGAAAATGTCATGCTCCG | ACADL |
| Acadm_1 | AAGATGTGGATAACCAACGG | ACADM |
| Acadm_2 | CCCGGAATATGACAAAAGCG | ACADM |
| Acadm_3 | TCGAACACAACACTCGAAAG | ACADM |
| Acadm_4 | AGCAGGTTTCAAGATCGCAA | ACADM |
| Acads_1 | GGTGACTCATGGGTCCTCAA | ACADS |
| Acads_2 | GCTCATGATAACTCCCGTGG | ACADS |
| Acads_3 | GATGGGCTTCAAAATAGCCA | ACADS |
| Acads_4 | GCTCACCCATCTTCTTAACC | ACADS |
| Ache_1 | CAGCAGACCTACATTGCCAG | ACHE |
| Ache_2 | AAATGTCTGCTACCAGTACG | ACHE |
| Ache_3 | GACCCCGTAAACCAGAAAGT | ACHE |
| Ache_4 | CAGTATGTGCATGCCCACGG | ACHE |
| Acly_1 | GAGAGAGATTGACCCCGACG | ACLY |
| Acly_2 | AGAGCGATTCGAGATTACCA | ACLY |
| Acly_3 | TTGTCACCTGTACACGACGG | ACLY |
| Acly_4 | GGACGAAAAGCTGAATACCG | ACLY |
| Aco1_1 | CCACCGCCATTTAGGCCGCG | ACO1 |
| Aco1_2 | CATATGCTATTACCAGAGGG | ACO1 |
| Aco1_3 | TAGCCCACCACATCAAACCT | ACO1 |
| Aco1_4 | GTGTAGCACCTCCGACAAGT | ACO1 |
| Aco2_1 | AGATCCGTGCCACTATTGAG | ACO2 |
| Aco2_2 | GCGTTTACGGCCCGACCGGG | ACO2 |
| Aco2_3 | TGGGTGAGGTCCAACCAGAG | ACO2 |
| Aco2_4 | GATTGAAATTAACCTCAATG | ACO2 |
| Acot9_1 | GAACCCTTACACATCCATCA | ACOT9 |
| Acot9_2 | CTGGGTTGGAAATACATCCA | ACOT9 |
| Acot9_3 | ACATCTTGGTAGAATGATTG | ACOT9 |
| Acot9_4 | GAAGAAGAGCTCTTTAAACA | ACOT9 |
| Acox1_1 | CGATCCAGACTTCCAACATG | ACOX1 |
| Acox1_2 | AATGTCGGATGGCTTGCGGT | ACOX1 |
| Acox1_3 | CCTCACAGCACTGTATCGAA | ACOX1 |
| Acox1_4 | TCTCTTCATAACCAAACTTG | ACOX1 |
| Acox2_1 | TAGGCCGAAGACGAGACTGG | ACOX2 |
| Acox2_2 | GATGAGCAGATTGCTAAATG | ACOX2 |
| Acox2_3 | CTTGAGTGACCAGTGTCGGT | ACOX2 |
| Acox2_4 | ATGCCAAGATAATTGCTCTG | ACOX2 |
| Acox3_1 | CAAGACGGCAACTCATGCGG | ACOX3 |
| Acox3_2 | GCCTCATGGACAGCCCAGCG | ACOX3 |
| Acox3_3 | TCAGCACTAGAATCTTCAGG | ACOX3 |
| Acox3_4 | GCGCATACTTCCGTACCTGG | ACOX3 |
| Acsl1_1 | GATGTCAGAACCATGTACGA | ACSL1 |
| Acsl1_2 | ATCACCTACATAGTGAACAA | ACSL1 |
| Acsl1_3 | TTCCACCAGATCACTGCCGT | ACSL1 |
| Acsl1_4 | TATGTTTGAGACCGTTGTAG | ACSL1 |
| Acsl3_1 | TGATCACAACGTACCCACGT | ACSL3 |
| Acsl3_2 | ACATCATTGCTTCTATAACG | ACSL3 |
| Acsl3_3 | TAATGATATGCCGCAGACGT | ACSL3 |
| Acsl3_4 | TTATCAAGTGTATCGCAGCC | ACSL3 |
| Acsl4_1 | GTCCAGGGATACGTTCACAC | ACSL4 |
| Acsl4_2 | GCCCATATCCCTGACCAATG | ACSL4 |
| Acsl4_3 | CAATAGAGCAGAGTACCCTG | ACSL4 |
| Acsl4_4 | GGAACAGCGGCCATAAGTGT | ACSL4 |
| Acsl5_1 | CTCTGCTCGATCAGACACCT | ACSL5 |
| Acsl5_2 | TAGACCCTGTTAAGGAGTCG | ACSL5 |
| Acsl5_3 | CGTCATCAAAAGGGTCCATG | ACSL5 |
| Acsl5_4 | GTACCCAGCGTGTCATACAG | ACSL5 |
| Acsl6_1 | AGAGAATCATCATGCCCCCG | ACSL6 |
| Acsl6_2 | CTGCTACACGTATTCCATGG | ACSL6 |
| Acsl6_3 | TGGCAGTAGACAACAGACTG | ACSL6 |
| Acsl6_4 | GCCCGGCGATCTGTGATTGG | ACSL6 |
| Acy1_1 | AATCTGACCAAGCTTGAAGG | ACY1 |
| Acy1_2 | TGTATCCTCAATGAAGCGTG | ACY1 |
| Acy1_3 | GACAGGTACCACATCTGTGT | ACY1 |
| Acy1_4 | AAAGACAGTAAAGGCATCCG | ACY1 |
| Adh1_1 | CCTTCGCGCAAAGTCCCCCG | Adh1 |
| Adh1_2 | CTGCCGCTCAGACGATCACG | Adh1 |
| Adh1_3 | TCACAAACCCTTCACCATCG | Adh1 |
| Adh1_4 | GGGGTGACTTGTGTGAAACC | Adh1 |
| Adh4_1 | GGAGTGACCAACTTCAAACC | ADH4 |
| Adh4_2 | TTCATTCTCTCAATACACTG | ADH4 |
| Adh4_3 | TGAGCTCCTGATCAATAGTG | ADH4 |
| Adh4_4 | TCAATTTCTTCAATGCAAAG | ADH4 |
| Adh5_1 | AAGACGGCACAAGTAGAACC | ADH5 |
| Adh5_2 | CCCCATCTGTCATCTCAACG | ADH5 |
| Adh5_3 | CAATAAAGATAAATTCGCAA | ADH5 |
| Adh5_4 | ACTTTATCCAAAGGGGCCGA | ADH5 |
| Adh7_1 | ACCGTCAACCTTCGCTACGG | ADH7 |
| Adh7_2 | AACAGCAGCTCCATAACCCG | ADH7 |
| Adh7_3 | TTTCTCCATTGAGGAAATCG | ADH7 |
| Adh7_4 | CACAGAGTGTATCAGTCCCA | ADH7 |
| Adk_1 | TGTGCTGCGTGTATCACTGG | ADK |
| Adk_2 | CACTTTCAATACCGACTCTG | ADK |
| Adk_3 | ATAAGGCATGACGTCCATCA | ADK |
| Adk_4 | CATTGGGATAGATAAGTTTG | ADK |
| Ahcy_1 | GCGCACCTGACAGAAGCTGT | AHCY |
| Ahcy_2 | TGTCAACGATTCTGTCACCA | AHCY |
| Ahcy_3 | TGACCCTATCATACCCTCCA | AHCY |
| Ahcy_4 | TGTGATGATTGCGGGCAAGG | AHCY |
| Ak2_1 | TGAAGGCGACAATGGATGCA | AK2 |
| Ak2_2 | GGTAGGACCGGCCACTCTTG | AK2 |
| Ak2_3 | TCCGAACCGGAGATTCCGAA | AK2 |
| Ak2_4 | TCAGCCAGTTTGGGTGCCTG | AK2 |
| Akr1b3_1 | TGAAAGTTGCTATTGACTTG | Akr1b3 |
| Akr1b3_2 | GGATCTCTTCATTGTCAGCA | Akr1b3 |
| Akr1b3_3 | ACCTTATTCACTGGCCAACG | Akr1b3 |
| Akr1b3_4 | TCTCCTGAGTTAGGTACGGG | Akr1b3 |
| Alad_1 | AAATGGAGCATTCCTAGCAG | ALAD |
| Alad_2 | ACGGACAGCCTCAATAGTTG | ALAD |
| Alad_3 | GCTCCGTCAGACATGATGGA | ALAD |
| Alad_4 | AAAACATGGACTTGGCAACA | ALAD |
| Alas1_1 | TGGATGAGGTCCATGCAGTG | ALAS1 |
| Alas1_2 | TGAGATGGTTAACGTCATTG | ALAS1 |
| Alas1_3 | TGCAGGAAATGCATGCTGTG | ALAS1 |
| Alas1_4 | AGAGCGCCGCATCTTTGCCG | ALAS1 |
| Alas2_1 | ATCCAAGGCATTCGCAACAG | ALAS2 |
| Alas2_2 | ACATCACACAATTCCTCCAG | ALAS2 |
| Alas2_3 | AATACTAAATAGGAACTGGT | ALAS2 |
| Alas2_4 | AGGTACCAGCAAGTTTCATG | ALAS2 |
| Aldh18a1_1 | CAAGTCTAGAGTGGGCCTAG | ALDH18A1 |
| Aldh18a1_2 | CGATGGGGACGATGTTCATG | ALDH18A1 |
| Aldh18a1_3 | CGTCCGAGAGGACAATCAAG | ALDH18A1 |
| Aldh18a1_4 | TGAGGGGTACCGTGATAAAG | ALDH18A1 |
| Aldh1a1_1 | CTTGTCGACATCCATGTGAG | ALDH1A1 |
| Aldh1a1_2 | CGTGAACCTATTGGAGTGTG | ALDH1A1 |
| Aldh1a1_3 | TAAATCCGACAAGTATGCAT | ALDH1A1 |
| Aldh1a1_4 | AGCAGAGCAAACTCCTCTCA | ALDH1A1 |
| Aldh1a2_1 | AGGATATTGACGACTCCGGG | ALDH1A2 |
| Aldh1a2_2 | ATTGTGCTGTGGTAACACCG | ALDH1A2 |
| Aldh1a2_3 | CCAGCCTGCATAATACCTCA | ALDH1A2 |
| Aldh1a2_4 | CTTGGTTGAAGAACACACCC | ALDH1A2 |
| Aldh1a3_1 | CACCAGGCATGAGCCCATCG | ALDH1A3 |
| Aldh1a3_2 | TGCATCCAGCCGGCGCCACG | ALDH1A3 |
| Aldh1a3_3 | CCACCCGGCAAAATATCTGA | ALDH1A3 |
| Aldh1a3_4 | AACACTAGAGAAAATATGTG | ALDH1A3 |
| Aldh1b1_1 | TTCACCCGACATGAGCCAGT | ALDH1B1 |
| Aldh1b1_2 | GTTCCACAAGATCAGCTAGG | ALDH1B1 |
| Aldh1b1_3 | CAACAACGAGTGGCATGATG | ALDH1B1 |
| Aldh1b1_4 | CATCACTGGCTACGGCCCCA | ALDH1B1 |
| Aldh2_1 | CACCGTCAACCCTTCCACAG | ALDH2 |
| Aldh2_2 | ACGGGACCGGACCTACCTAG | ALDH2 |
| Aldh2_3 | AGCCCCCAACCATCAGCCTG | ALDH2 |
| Aldh2_4 | CAAGTTGGCCACGTAGAGCG | ALDH2 |
| Aldh3a1_1 | TGAATGGACCTCCTACTACG | ALDH3A1 |
| Aldh3a1_2 | AGTCCTTGTCTACGTAACAA | ALDH3A1 |
| Aldh3a1_3 | CTGTACCCAGTGATCAAAGG | ALDH3A1 |
| Aldh3a1_4 | ATGTGATCACTCACTTCCGA | ALDH3A1 |
| Aldh3a2_1 | CTGTATGCGATTGTTAATGG | ALDH3A2 |
| Aldh3a2_2 | GGAAATCTTAGCAGCCATCG | ALDH3A2 |
| Aldh3a2_3 | AGTCTCTGTCAATGTAACAA | ALDH3A2 |
| Aldh3a2_4 | TCTGGAGGGAGGCTTCGCAC | ALDH3A2 |
| Aldh7a1_1 | GGGCTCCGCGAGGATAACGA | ALDH7A1 |
| Aldh7a1_2 | TGGTGGAGCAGATATCGGGT | ALDH7A1 |
| Aldh7a1_3 | ACACTAACGAGGGACGTAGT | ALDH7A1 |
| Aldh7a1_4 | TAGATTCCTGCCCCAAAACG | ALDH7A1 |
| Alox5_1 | CGTACTTTGAATCCGTAGGG | ALOX5 |
| Alox5_2 | GCACAGACGTAAAGAACTGG | ALOX5 |
| Alox5_3 | GCTGGTTGTACCTTGAAAAG | ALOX5 |
| Alox5_4 | GTTGATTACGAGCTACTGGA | ALOX5 |
| Amd1_1 | TCTTCGTACCATCCCAAGGT | AMD1 |
| Amd1_2 | TCTTCCTGGAAATTCCGGTG | AMD1 |
| Amd1_3 | AGCAGGAAGCTTATGTACTC | AMD1 |
| Amd1_4 | AAGCTTGCTAGGGATTACAG | AMD1 |
| Aprt_1 | GAGTCCGGGTCTTTCAAGAG | APRT |
| Aprt_2 | CTAACAGGTCTAGACTCCAG | APRT |
| Aprt_3 | TGTGTGCTCATCCGGAAACA | APRT |
| Aprt_4 | CGCACCTGAACAGCACGCCC | APRT |
| Aqp9_1 | ACATTGCAAAAGACACCGCT | AQP9 |
| Aqp9_2 | GTCTTCTATAGACGGACTCA | AQP9 |
| Aqp9_3 | TCAGTCGAGAAAAGGCTGGT | AQP9 |
| Aqp9_4 | GCTGAGCTCCCACATAAAAG | AQP9 |
| Asns_1 | GCATGCCATCTATGACAGCG | ASNS |
| Asns_2 | GAATGCAGCCGATAAGAGTG | ASNS |
| Asns_3 | GCTGTGTGTTCAGAAGCTAA | ASNS |
| Asns_4 | ATTTGAATATCAGACCAATG | ASNS |
| Ass1_1 | TAACATCTTACCTTAATCTG | ASS1 |
| Ass1_2 | TGGGATCCTGGAAAACCCCA | ASS1 |
| Ass1_3 | AAGGCACTTCCTCACCAGGT | ASS1 |
| Ass1_4 | TGCCTTCACCTGTAGCAACA | ASS1 |
| Atp11a_1 | ACCTGAACGACCCTGTAGTG | ATP11A |
| Atp11a_2 | CAGTGCAACCTCATCAGGCG | ATP11A |
| Atp11a_3 | ACTTTGCCTTCTATCACTCG | ATP11A |
| Atp11a_4 | ACACGGTGCTGAAGTACGTG | ATP11A |
| Atp2a2_1 | TGTCTTGATTAACAGCTCGG | ATP2A2 |
| Atp2a2_2 | GGACCTTGGTACTTCCAACT | ATP2A2 |
| Atp2a2_3 | GATAACAGAAGTACAACCAA | ATP2A2 |
| Atp2a2_4 | GAGGCTACTTCAATCCTGGG | ATP2A2 |
| Atp2c1_1 | TGATGCCGTCAGTATCACTG | ATP2C1 |
| Atp2c1_2 | GAGGGTCAATGATTCCGACC | ATP2C1 |
| Atp2c1_3 | GAAGTAACATCGCCTTCATG | ATP2C1 |
| Atp2c1_4 | CATGCAGGCCATCGGAAGTG | ATP2C1 |
| Atp5g3_1 | GCCCGGGTTACGCACCAGAG | ATP5G3 |
| Atp5g3_2 | AATGTCTCTGCTGATTACAC | ATP5G3 |
| Atp5g3_3 | TCTCGGCCAGAGACTAGGAC | ATP5G3 |
| Atp5g3_4 | GTGTGTGTCAGCTGATCCGA | ATP5G3 |
| B4galt1_1 | GGCCAGAGAGGTAATAGACG | B4GALT1 |
| B4galt1_2 | CAGGGCTGGAGTCGAGACCC | B4GALT1 |
| B4galt1_3 | GCTCAATATTGGCTTTCAAG | B4GALT1 |
| B4galt1_4 | TGATGTGGACCTCATTCCGA | B4GALT1 |
| B4galt2_1 | GATAGTGGAGCCAATAGCGT | B4GALT2 |
| B4galt2_2 | GGGCATAGACGTCAAAGTAG | B4GALT2 |
| B4galt2_3 | CGGGTGCAGAGGGAAAATCC | B4GALT2 |
| B4galt2_4 | GGCGGCACGTCAGGACAGGG | B4GALT2 |
| Bckdha_1 | CATGACCAACTATGGCGAGG | BCKDHA |
| Bckdha_2 | CAGCGAAATTGAAACCGGCG | BCKDHA |
| Bckdha_3 | CTTGTCTACAAACTCTGCAG | BCKDHA |
| Bckdha_4 | CTGGCCACGCAGATCCCTCA | BCKDHA |
| Bckdk_1 | ACAGCACGAGCTATACATCC | BCKDK |
| Bckdk_2 | AGTCCGCATCAACGGACATG | BCKDK |
| Bckdk_3 | GCGACCGGAATAGAGCATCA | BCKDK |
| Bckdk_4 | TGTCTTCATGTAGCGCCAAG | BCKDK |
| Cad_1 | CGCAGGGGTACCCGACCGTG | CAD |
| Cad_2 | AGGATTAGAACCTTTCGTGG | CAD |
| Cad_3 | ATGGTGAGTGCCCACCACAA | CAD |
| Cad_4 | CTCAGAAACTCTGTTACGGG | CAD |
| Cbs_1 | ACACTACGATGACACCGCCG | CBS |
| Cbs_2 | TCGCCATGCCACTCCCACGT | CBS |
| Cbs_3 | GTGAGTTCTTCAATGCGGGT | CBS |
| Cbs_4 | ATAATGTGGGGAGTCCGCCA | CBS |
| Cel_1 | ACCCTCTCCCCATACAACAA | CEL |
| Cel_2 | GTGGACACACTGTCGCCAAG | CEL |
| Cel_3 | CAAAAACTACCTGTATGACG | CEL |
| Cel_4 | AGGCCAGGTAGTGCACAACA | CEL |
| Chat_1 | AGTCAAAATGGCGTCCAACG | CHAT |
| Chat_2 | CAGGACGATGCCATCAAAAG | CHAT |
| Chat_3 | TGAGCCCGAGCATGTCATCG | CHAT |
| Chat_4 | GGCCCCAAACCGCTTCACAA | CHAT |
| Chka_1 | AGTGCTCTTGCGGCTCTATG | CHKA |
| Chka_2 | GGTTGTTCGTCCGCTAGCGG | CHKA |
| Chka_3 | AAGACTCAAATTCAGCAGGG | CHKA |
| Chka_4 | AGCCTGGCGAGGCCTTCGCG | CHKA |
| Chkb_1 | ATGGTCCCAAACAACCATCG | CHKB |
| Chkb_2 | GGTGCTGCTACGACTCTACG | CHKB |
| Chkb_3 | CGTGGTTCGGTAGTGAGCAT | CHKB |
| Chkb_4 | CAGAGCCAATCCCCAATCGG | CHKB |
| Cox4i1_1 | TGCTCGCGACCGGAAGAACG | COX4I1 |
| Cox4i1_2 | CACGCCGATCAGCGTAAGTG | COX4I1 |
| Cox4i1_3 | AGCTTCGCCGAGATGAACAG | COX4I1 |
| Cox4i1_4 | GGCAGACAGCATCGTGACAT | COX4I1 |
| Cox4i2_1 | GGGCGTAGCAGTCAACGTAG | COX4I2 |
| Cox4i2_2 | CCAGCGCTCCTATCCCATGC | COX4I2 |
| Cox4i2_3 | GGACAGGACTCAGAACTAGA | COX4I2 |
| Cox4i2_4 | CTTCTGCACAGAGCTCAGCG | COX4I2 |
| Cox5a_1 | CCGTCGCTGTACCGCAGCCG | COX5A |
| Cox5a_2 | CCCATGAGAATAGCAGCGAA | COX5A |
| Cox5a_3 | TGAGGAGTTTGATGCTCGCT | COX5A |
| Cox5a_4 | CTGCATTGCGAGCATGTAGA | COX5A |
| Cox5b_1 | TAGCCTGCTCCTCATCAGTG | COX5B |
| Cox5b_2 | AAAGGCAGCTTCAGGCACCA | COX5B |
| Cox5b_3 | TCACGTACCTCCAGAAGCCA | COX5B |
| Cox5b_4 | ATCATGATAGCAGCACAGAA | COX5B |
| Cox6a1_1 | TGAGGGTAGGCAACGAACGG | COX6A1 |
| Cox6a1_2 | CCGTGGGCGCCACTCGACAT | COX6A1 |
| Cox6a1_3 | CGCCGCAGCTCGGATGTGGA | COX6A1 |
| Cox6a1_4 | TCAACGTGTTCCTCAAGTCG | COX6A1 |
| Cox6a2_1 | CTTAACTGCTGGATGCACGC | COX6A2 |
| Cox6a2_2 | GCCGGGAAGAGCCAGCACAA | COX6A2 |
| Cox6a2_3 | GGTGATACGGGATGAACTCT | COX6A2 |
| Cox6a2_4 | AGGGAGCAGAGGGCTACGCC | COX6A2 |
| Cox6b1_1 | AAGGCAATGACGGCCAAGGG | COX6B1 |
| Cox6b1_2 | AACTGTTGGCAGAACTACCT | COX6B1 |
| Cox6b1_3 | AGAACCAGACTAAGAACTGT | COX6B1 |
| Cox6b1_4 | ACGCCGGTACCACTCACACA | COX6B1 |
| Cox6b2_1 | GTGAAGACCATGAATCGCCG | COX6B2 |
| Cox6b2_2 | ACACGGAAATAGTACTCGCA | COX6B2 |
| Cox6b2_3 | GGGAAGCGCGGATCAAAGGG | COX6B2 |
| Cox6b2_4 | CGTTGTCCATTGGCCCGGAG | COX6B2 |
| Cox6c_1 | TATTGCTGGCGCATTCATTG | COX6C |
| Cox6c_2 | AATATGAACCCGCAGACGCT | COX6C |
| Cox6c_3 | AGCACATACCTTATAGGCAG | COX6C |
| Cox6c_4 | TGGTTTGGGCAACAGCGCAC | COX6C |
| Cox7a1_1 | GACCTCCCAGTACACTTGAA | COX7A1 |
| Cox7a1_2 | AAGCCACTTAGAAAACCGTG | COX7A1 |
| Cox7a1_3 | TGGTAGATGAGCTAAAAGAC | COX7A1 |
| Cox7a1_4 | AAAAGACCGGACCAGAGCCT | COX7A1 |
| Cox7a2_1 | GGGATGCCAGTTCATCTGAA | COX7A2 |
| Cox7a2_2 | GAGCCATTGTGGCTCTGTAG | COX7A2 |
| Cox7a2_3 | ATGTTGCGGAATCTGCTGGT | COX7A2 |
| Cox7a2_4 | CTGGCATCCCATTATCCTCC | COX7A2 |
| Cox7b_1 | TTATCTGCTCACCTTGGAGA | COX7B |
| Cox7b_2 | GGGAATGCTATATTAGCAGG | COX7B |
| Cox7b_3 | AACTAGGTGCCCTCTTCTGG | COX7B |
| Cox7b_4 | TAGCATTCCCATATTTGTCA | COX7B |
| Cox7b2_1 | GCAATATGATGCTAATTAGT | COX7B2 |
| Cox7b2_2 | GTTTAGTGCATATCTGGCCA | COX7B2 |
| Cox7b2_3 | CCAAGCATTCTGAAAATTGT | COX7B2 |
| Cox7b2_4 | GAAGAAACTCATGACAAATA | COX7B2 |
| Cox7c_1 | CCGGAGGTTCACGACCTCCG | COX7C |
| Cox7c_2 | GCCACTATGAGGAGGGTCCG | COX7C |
| Cox7c_3 | TTTCAGTGGAAAACAAGTGG | COX7C |
| Cox7c_4 | TCCCCGGACCCTCCTCATAG | COX7C |
| Cox8a_1 | GCCTGACCGGCTCGGCCCGG | COX8A |
| Cox8a_2 | TTCGAGTGGACCTGAGCCCG | COX8A |
| Cox8a_3 | TCTGTGTAGGATATCACCAT | COX8A |
| Cox8a_4 | GCTCCCGCGCCGGCTTCGAG | COX8A |
| Cox8c_1 | GCTTCGAGAACAGGACTGCA | COX8C |
| Cox8c_2 | AAAAAAAATCCTGTCGCCCA | COX8C |
| Cox8c_3 | TTCTCGAAGCCTGGCCACCC | COX8C |
| Cox8c_4 | GGCTTTCTGAGTGGCTGAGG | COX8C |
| Cp_1 | CATATAAGCATCAATTAGGG | CP |
| Cp_2 | GCTGTGAGGAGCGACCTGGT | CP |
| Cp_3 | ATGAAAAGTGTAGATCCTAG | CP |
| Cp_4 | GCTGAACAAATACCACACGA | CP |
| Cpox_1 | TGGGCGCATAAGGATTCTTG | CPOX |
| Cpox_2 | GAAGCAGCGAACCAAATGAG | CPOX |
| Cpox_3 | ACCGACATACTTGAACCAAG | CPOX |
| Cpox_4 | CGGACCCGCCTCTAGCCCAG | CPOX |
| Cps1_1 | TGAGCCTCACAATTTCGTCG | CPS1 |
| Cps1_2 | ATGCAGACCGAATCATCACA | CPS1 |
| Cps1_3 | TACAGTATTCCATGGAAGTG | CPS1 |
| Cps1_4 | GTTGGTGGCATCTCGTGTCG | CPS1 |
| Cpt1a_1 | CACATTGTCGTGTACCACAG | CPT1A |
| Cpt1a_2 | CATACTGCTGTATCGTCGCA | CPT1A |
| Cpt1a_3 | ACCTTGGACCCAAATTGCAG | CPT1A |
| Cpt1a_4 | ACGTTGGACGAATCGGAACA | CPT1A |
| Csad_1 | ATTCACTACAGTGTCAAGAC | CSAD |
| Csad_2 | ACACGGACTGTGGTTCCATG | CSAD |
| Csad_3 | CTTGACCACTCGGACACTGT | CSAD |
| Csad_4 | TCTCCTCCTAGGTCTGCGAA | CSAD |
| Cth_1 | TGCATGGATGAAGTGTATGG | CTH |
| Cth_2 | GCAATGGAATTCTCGTGCCG | CTH |
| Cth_3 | TTCCAAAACCAAATTGCTAG | CTH |
| Cth_4 | TTTGTGGACAATTTGTGCGC | CTH |
| Ctps_1 | ATTGGCCATTAACCACAAGC | CTPS |
| Ctps_2 | ATACCAGTACGTCATTAACA | CTPS |
| Ctps_3 | GCCCACAAGAGCGATCGAGC | CTPS |
| Ctps_4 | TTAATACCCGTAGACGAAGA | CTPS |
| Cubn_1 | GTGTATCTGGAACATTCGCG | CUBN |
| Cubn_2 | AGTTATCAACTTCACCCACG | CUBN |
| Cubn_3 | ACAGCTCCGAATGCTACTGG | CUBN |
| Cubn_4 | CCACCTGTGTGAACACTATG | CUBN |
| Dbt_1 | TAGATGATATCGCTTATGTG | DBT |
| Dbt_2 | TCATCACAATACCCGAAGTG | DBT |
| Dbt_3 | CTGTGAAGTTCAAAGTGACA | DBT |
| Dbt_4 | TGGAGGCTTTGCTATCGGCG | DBT |
| Dck_1 | CAACGTGCAGAGCACTCAAG | DCK |
| Dck_2 | TGTATGAGAAACCTGAACGG | DCK |
| Dck_3 | AGTGGACCATATATCAAGAC | DCK |
| Dck_4 | TTTGAGCTTGCCATTGAGAG | DCK |
| Ddc_1 | CCTAGGTGGTCGCTACACTG | DDC |
| Ddc_2 | TAACCCAGCTCTTTCTACAG | DDC |
| Ddc_3 | CCGGTATCTTCTGAATGGTG | DDC |
| Ddc_4 | AGCCAGTAGGGCCACCAAGG | DDC |
| Dhfr_1 | GACATGGTTTGGATAGTCGG | DHFR |
| Dhfr_2 | AACCTCAGAGAACCACCACG | DHFR |
| Dhfr_3 | TCGCCGTGTCCCAAAATATG | DHFR |
| Dhfr_4 | CAGCCCGGCCAATACCTGAG | DHFR |
| Dhodh_1 | TTGATCCAGAGTCGGCGCAC | DHODH |
| Dhodh_2 | GTAGAAATGGTCGTCCCCCG | DHODH |
| Dhodh_3 | ATAAATTCCGAAATCCAGTA | DHODH |
| Dhodh_4 | GGTATGGATTCAACAGCCAC | DHODH |
| Dhtkd1_1 | CTGATGTTCCGTAAAATGCG | DHTKD1 |
| Dhtkd1_2 | GCCACGGTGAAGAGATACGG | DHTKD1 |
| Dhtkd1_3 | TCTTCTTTAGGAAAGCTCGT | DHTKD1 |
| Dhtkd1_4 | GATGGGGACTACTCCCCGAA | DHTKD1 |
| Dlat_1 | ACTACCGCAACGGACCGCAG | DLAT |
| Dlat_2 | CAGGCTCTCAAACCCAACAG | DLAT |
| Dlat_3 | CGACAAGGCCACCATAGGTG | DLAT |
| Dlat_4 | TTCAGAACCACACCTACCGG | DLAT |
| Dld_1 | GGTGGAACATGCTTGAACGT | DLD |
| Dld_2 | GCAGTAAAAGCATTAACAGG | DLD |
| Dld_3 | AGAGAAGCTGGTTGTTATTG | DLD |
| Dld_4 | CAAAAACATCCTTGTAGCTA | DLD |
| Dpyd_1 | CTATAACCTTCCGTGGCGAG | DPYD |
| Dpyd_2 | GAGGCACAGCTTATACTCGC | DPYD |
| Dpyd_3 | TTTGTCCAGGGCCATACAAG | DPYD |
| Dpyd_4 | AGAGCTAAGAGTCATTTCAT | DPYD |
| Dtymk_1 | TGGTGTAAACAACCGGACGT | DTYMK |
| Dtymk_2 | ATTAAGGCGAAGTTGAACCA | DTYMK |
| Dtymk_3 | CGTGCTGGCAAGACCACGCA | DTYMK |
| Dtymk_4 | CGTCCAGCAATTGTAACTGA | DTYMK |
| Ehhadh_1 | TTCATATGGATGCTTCACGG | EHHADH |
| Ehhadh_2 | CCACATCATGAGGTTACTAG | EHHADH |
| Ehhadh_3 | GTAAACCCATAGAACCCCGC | EHHADH |
| Ehhadh_4 | TCACTATGGCTCTAACCGTA | EHHADH |
| Fads2_1 | ACGTTACCAAATGGTCCCAG | FADS2 |
| Fads2_2 | AACACGATTCATACCTCAAG | FADS2 |
| Fads2_3 | CATGACAATGATCAGCCGCA | FADS2 |
| Fads2_4 | CGAGGACAAAGGCTGTGACG | FADS2 |
| Fasn_1 | CTACCAGGCCATCCGTAGTG | FASN |
| Fasn_2 | TGTCTCCGAAAAGAGCCGGG | FASN |
| Fasn_3 | TTGGTGGAGCCAATTAACAG | FASN |
| Fasn_4 | ACTGGCAATCTGATTGTGAG | FASN |
| Fbp1_1 | CATGGCAAGGACCAACATGG | FBP1 |
| Fbp1_2 | CACAAGAACACAGGTAGCGT | FBP1 |
| Fbp1_3 | TGGCTCAACCAATGTGACTG | FBP1 |
| Fbp1_4 | AACATCTACAGCCTTAATGA | FBP1 |
| Fbp2_1 | ACTGTATGGTAGTGCAACCC | FBP2 |
| Fbp2_2 | CCCGTTACGTTATGGAAAAG | FBP2 |
| Fbp2_3 | TTGTCTTCCACAGACCACGG | FBP2 |
| Fbp2_4 | TGTGATGTAGGGGAAATATG | FBP2 |
| Flad1_1 | ACAAACTCGCGGAGTCAGGT | FLAD1 |
| Flad1_2 | TCACTCACGTCCTCACCGCG | FLAD1 |
| Flad1_3 | GCTAAGCCTACGCCCAAAGT | FLAD1 |
| Flad1_4 | GAAGCTGATTCTAGACTCCG | FLAD1 |
| Folr1_1 | GACAATTTACACGACCAGGT | FOLR1 |
| Folr1_2 | AGTTCGGGGAACACTCATAG | FOLR1 |
| Folr1_3 | CCAGTTGAACCGGTACAGGT | FOLR1 |
| Folr1_4 | CTCCACCTACTCCTTACCCG | FOLR1 |
| G6pc_1 | GGTGTTTGAACGTCATCTTG | G6PC |
| G6pc_2 | GTTGTCCAAACAGAATCCTG | G6PC |
| G6pc_3 | GGTCAGCAATCACAGACACA | G6PC |
| G6pc_4 | ACAAGACTCCAGCCACGACC | G6PC |
| G6pc2_1 | GACCTGATGGGGGAAATGTG | G6PC2 |
| G6pc2_2 | GCAGGTCAATACCGAACAGT | G6PC2 |
| G6pc2_3 | GGGTAGCGGTCATAGGGGAC | G6PC2 |
| G6pc2_4 | AACACTCCACAGAAAGGACC | G6PC2 |
| G6pc3_1 | TAGGCCGACTGCCAATAGGA | G6PC3 |
| G6pc3_2 | TTCCCGGGCTAGAGAATATG | G6PC3 |
| G6pc3_3 | GGGGCTCATTAGCCAGCCAA | G6PC3 |
| G6pc3_4 | TAAAGAGAGTCCAATACATG | G6PC3 |
| G6pdx_1 | AGAGGTGGAAACTGACAACG | G6pdx |
| G6pdx_2 | TGCCCGCTCACGACTCACAG | G6pdx |
| G6pdx_3 | ATGACCCCACAGTACCCCAT | G6pdx |
| G6pdx_4 | AGAGATGGTCCAGAATCTCA | G6pdx |
| G6pd2_1 | GATGACCCCACAGTACCCCG | G6pd2 |
| G6pd2_2 | AGTGTTGAAACGTATCTCAG | G6pd2 |
| G6pd2_3 | CATCCACTGTGAGTTGTGAG | G6pd2 |
| G6pd2_4 | ATCATACTGGCCAACTACAT | G6pd2 |
| Gad1_1 | CTATTCCATAAAGAAAGCCG | GAD1 |
| Gad1_2 | GACATTTGATCGCTCCACCA | GAD1 |
| Gad1_3 | GCGGTTGCATTGACATAAAG | GAD1 |
| Gad1_4 | AGATGAGAGAGATCGTTGGA | GAD1 |
| Gad2_1 | TTCTGATTAAATGTGACGAG | GAD2 |
| Gad2_2 | GTATAAGATCTGGATGCACG | GAD2 |
| Gad2_3 | GGCATCGGAAACAAGCTGTG | GAD2 |
| Gad2_4 | GACAGCTGATTAAAATATCG | GAD2 |
| Gapdh_1 | GCTGTGGCGTGATGGCCGTG | GAPDH |
| Gapdh_2 | AAACAGGCCCACTTGAAGGG | GAPDH |
| Gapdh_3 | TGCCATTTGCAGTGGCAAAG | GAPDH |
| Gapdh_4 | GGCCGGTGCTGAGTATGTCG | GAPDH |
| Gart_1 | ACTCGTAGTTGTCGGACCAG | GART |
| Gart_2 | TGACGGCTTCAGTTGTACTG | GART |
| Gart_3 | GCTAGAAAGGATCACCGAAG | GART |
| Gart_4 | GGCAAAGTAGTGACCAGCGG | GART |
| Gatm_1 | ATCAAAGACTACTTCCATCG | GATM |
| Gatm_2 | ATTCGTTGTAAGAGGAGACA | GATM |
| Gatm_3 | ACTTGAGTGACCAGTCGATG | GATM |
| Gatm_4 | ACAACCATCAGGATGTCTCG | GATM |
| Gch1_1 | GCACCAATGGGTTCTCCGAG | GCH1 |
| Gch1_2 | CCCTTCCTACAAATGGAACA | GCH1 |
| Gch1_3 | GGTGAACCTCCCCAAACTGG | GCH1 |
| Gch1_4 | GTACTTCACCAAGGGATACC | GCH1 |
| Gck_1 | TTCTGGGGTGGAACGCACGT | GCK |
| Gck_2 | AAGGCACGAAGACATAGACA | GCK |
| Gck_3 | CCATCCGGTCGTACTCCAGG | GCK |
| Gck_4 | CCAGATGTATTCCATCCCCG | GCK |
| Gclc_1 | TGTGCCGGTCCTTGACTGCG | GCLC |
| Gclc_2 | CAATATGAGGAAACGCCGGA | GCLC |
| Gclc_3 | AGAAACATCCGGCATCGGAG | GCLC |
| Gclc_4 | TGTAGATGATAGAACACGGG | GCLC |
| Gclm_1 | TACTTACCCTGACTAAATCG | GCLM |
| Gclm_2 | GTGCCCGTCCACGCACAGCG | GCLM |
| Gclm_3 | TTAACTCCATCTTCAATCGG | GCLM |
| Gclm_4 | GCATTTACAGCCTTACTGGG | GCLM |
| Gpd1_1 | GATGGGGTACCACAAAAACC | Gpd1 |
| Gpd1_2 | ACCCAACTTTCGCATCACTG | Gpd1 |
| Gpd1_3 | GTCTTCCTCAAACACCCACA | Gpd1 |
| Gpd1_4 | GTTGGAGCTCACCACATTGG | Gpd1 |
| Gfpt1_1 | TGTGGCACAAGTTACCACGC | GFPT1 |
| Gfpt1_2 | AAGCTGCGGTCTTTCCCGTG | GFPT1 |
| Gfpt1_3 | GGAGAGAGGAGCCTTAACTG | GFPT1 |
| Gfpt1_4 | TCTGTTGTGAACACAATGAG | GFPT1 |
| Gfpt2_1 | CTGCGATACTGTAAGGATCG | GFPT2 |
| Gfpt2_2 | TACTGCACTTACAGCCACAG | GFPT2 |
| Gfpt2_3 | CACATGTCGGATATAAGACG | GFPT2 |
| Gfpt2_4 | AATTAGTGATGATCCCGTTG | GFPT2 |
| Ggt1_1 | ACTGACGTATCACCGTATCG | GGT1 |
| Ggt1_2 | CGTACAGGGTCGCATCACCG | GGT1 |
| Ggt1_3 | GAATTCAGGCTCATAGTAGG | GGT1 |
| Ggt1_4 | CGACCACGTGTACTCCAGGG | GGT1 |
| Ggt5_1 | GTGCGGAGGTCTTATACACA | GGT5 |
| Ggt5_2 | AGATCTGCTCGGATATTGGA | GGT5 |
| Ggt5_3 | TGCAGAGAAGTTGAACCCTG | GGT5 |
| Ggt5_4 | GCTGGGCTAGATCTTCCCGG | GGT5 |
| Ggt6_1 | TGACCTAAGCCTGTTGCATG | GGT6 |
| Ggt6_2 | CCATTGGTATGGCAGAACAA | GGT6 |
| Ggt6_3 | GGAGACTCCAAAATACCCCC | GGT6 |
| Ggt6_4 | GCAGCCACATGTTCCCAACT | GGT6 |
| Ggt7_1 | GGTCATGCAAATCTACTTCG | GGT7 |
| Ggt7_2 | CGCAGGCGGTGTCATAACCG | GGT7 |
| Ggt7_3 | GCGACGGCCTGATCTCGCTG | GGT7 |
| Ggt7_4 | TGTTGGTACATGATATCCGG | GGT7 |
| Gldc_1 | CCACTGAAATCCATTTCGTG | GLDC |
| Gldc_2 | TTTACTCAACTACCAGACCA | GLDC |
| Gldc_3 | CTTGATCCAGGGATGCCACA | GLDC |
| Gldc_4 | AGGCCACAAACAGCTACCAG | GLDC |
| Gls_1 | CGACGCGTTCGGCAACAGCG | GLS |
| Gls_2 | TGTACATCGCTATGTTGGGA | GLS |
| Gls_3 | GATTGCGAACATCTGATCCC | GLS |
| Gls_4 | ATATAACTCATCGATGTGTG | GLS |
| Gls2_1 | CGTCCGGTACTACCTCGGTG | GLS2 |
| Gls2_2 | GGGGATCGGAATTACGCCAT | GLS2 |
| Gls2_3 | AAAAGCAGGTCACCAAGTCG | GLS2 |
| Gls2_4 | TGAGTCAGGCAGTGTCATGG | GLS2 |
| Gmds_1 | CCTGTTGGAGAAAGGGTACG | GMDS |
| Gmds_2 | GAGAAGGCCTCACCTTGACA | GMDS |
| Gmds_3 | TCCTTACCATAGGGTGACCT | GMDS |
| Gmds_4 | CGAGATGGGCAAGCTCAGGA | GMDS |
| Gmps_1 | AGGGCTATTATCATATCTGG | GMPS |
| Gmps_2 | AGTGCGTCCCTTGTTGCCAG | GMPS |
| Gmps_3 | AAATATGACAACAAGTCCTG | GMPS |
| Gmps_4 | GAATGAGCAGCATTTATCAC | GMPS |
| Gne_1 | ACGTCCAACTCAAAGAACGC | GNE |
| Gne_2 | TTGCAGCTCAAAGATATATG | GNE |
| Gne_3 | GCTCCACACGATTGTTAGAG | GNE |
| Gne_4 | CCTCTTGTTAAACGAGATCA | GNE |
| Gpat2_1 | CTCCCGGAACCCTCCCCGAG | GPAT2 |
| Gpat2_2 | CAAGGAGAGGTTGACCTCTG | GPAT2 |
| Gpat2_3 | CTGAATGTACAACTTCACAA | GPAT2 |
| Gpat2_4 | AATGGCAACTGGTACCAGTG | GPAT2 |
| Gpd2_1 | CGTACCGTCATAGTAGACAA | GPD2 |
| Gpd2_2 | GGAGGTATCGCACACCACCG | GPD2 |
| Gpd2_3 | TGGCACCACTGATACGCCAA | GPD2 |
| Gpd2_4 | TGGGATCGGTAACAAGGGGA | GPD2 |
| Gpi1_1 | GTACACTGGCAAATCCATCA | GPI |
| Gpi1_2 | TTAGAGACAAACCAGACACG | GPI |
| Gpi1_3 | CGGCAAAGATGTGATGCCGG | GPI |
| Gpi1_4 | ACTTACCGTGTTCGTAGACA | GPI |
| Gpx4_1 | CGTGTGCATCGTCACCAACG | GPX4 |
| Gpx4_2 | CATGCCCGATATGCTGAGTG | GPX4 |
| Gpx4_3 | TGGTCTGGCAGGCACCATGG | GPX4 |
| Gpx4_4 | TAAGCCAGCACTGCTGTGCG | GPX4 |
| Gsta2_1 | CTTCACTACTTCAATGCCCG | GSTA2 |
| Gsta2_2 | AGGATTGACATGTATACAGA | GSTA2 |
| Gsta2_3 | AGACTGCCTTGGCAAAAGAC | GSTA2 |
| Gsta2_4 | GGAGATTGATGGGATGAAGC | GSTA2 |
| Gsta3_1 | GAAGAATGGAGCCTATCCGG | GSTA3 |
| Gsta3_2 | ATCCCGTCGATCTCTACCAT | GSTA3 |
| Gsta3_3 | GACCTGGCAAGGTTACGAAG | GSTA3 |
| Gsta3_4 | CCAAGATCAAGGAACAAACC | GSTA3 |
| Gsta4_1 | CAACGAGAAAAGCCTCTCCG | GSTA4 |
| Gsta4_2 | CATCCCATCGATTTCAACCA | GSTA4 |
| Gsta4_3 | CAAGTACAACTTGTATGGGA | GSTA4 |
| Gsta4_4 | CATGTATGCAGATGGCACCC | GSTA4 |
| Gstk1_1 | GGTTCCCCTCAACATACCCA | GSTK1 |
| Gstk1_2 | ATGATCCCAGCGATTAAAGT | GSTK1 |
| Gstk1_3 | CCTGCTATGGTTCCCCGAAA | GSTK1 |
| Gstk1_4 | GCTCCATGCTTACAGTGGTG | GSTK1 |
| Gstm1_1 | CTTGCCCGAAAGCACCACCT | GSTM1 |
| Gstm1_2 | ACTGGCCACTCACAAAGTCA | GSTM1 |
| Gstm1_3 | TGGACCCCTCTCTCACTCCG | GSTM1 |
| Gstm1_4 | GGTGGCATTACCGTCACCCA | GSTM1 |
| Gstm2_1 | AGGACAAGAAATACACCATG | GSTM2 |
| Gstm2_2 | AAGCGTTCTCACTCACCCCA | GSTM2 |
| Gstm2_3 | AGCTCCAACTCACAAAGTCA | GSTM2 |
| Gstm2_4 | AGTAGAGCTTCATCTTCTCA | GSTM2 |
| Gstm3_1 | GTATGCGGGTGTCCATAACT | GSTM3 |
| Gstm3_2 | GTATTCCAGGAGCAAGCGGA | GSTM3 |
| Gstm3_3 | GGAGAAGAGATATGTCATGG | GSTM3 |
| Gstm3_4 | CACTTACATTGGGAAAATCC | GSTM3 |
| Gstm4_1 | AGAAGAGCTTCACCATTCCA | GSTM4 |
| Gstm4_2 | TCAGGTGGTTGCTCACCCCA | GSTM4 |
| Gstm4_3 | GCTTGCGGGCAATGTAGCGC | GSTM4 |
| Gstm4_4 | ACAGACTCGAGCCAGCTGAT | GSTM4 |
| Gstm5_1 | CGTGGTTAGAATTGTAGCAG | GSTM5 |
| Gstm5_2 | CAAACCATGTGAATTTCCCC | GSTM5 |
| Gstm5_3 | GCGTGCGATGTATCTCAGGA | GSTM5 |
| Gstm5_4 | AAAGATACGAGTAGACATCA | GSTM5 |
| Gsto1_1 | ATACCTTAGAGAATGACTCA | GSTO1 |
| Gsto1_2 | ATTCGGTGACCAAGTGACCC | GSTO1 |
| Gsto1_3 | GGAAGACTCTCCGAACCTAA | GSTO1 |
| Gsto1_4 | TCAGCGTCCTCTGAGCGAAG | GSTO1 |
| Gsto2_1 | GTTGCACAACTCCTGACGCA | GSTO2 |
| Gsto2_2 | TGATCCGAATCTACAGCATG | GSTO2 |
| Gsto2_3 | TACCTGGATGACGTCTACCC | GSTO2 |
| Gsto2_4 | CTAGGTCCCGCCTTTAAGCA | GSTO2 |
| Gstp1_1 | ATTCACCATATCCATCTGGG | GSTP1 |
| Gstp1_2 | GGGTGACATATTTGCCGCGA | GSTP1 |
| Gstp1_3 | ACTCCATACAGGGCGGTGTG | GSTP1 |
| Gstp1_4 | GGTCTCCATCCTCAAACTTG | GSTP1 |
| Gstt1_1 | CCCGTTCCAGATGCACACGG | GSTT1 |
| Gstt1_2 | AACATCCAGTTCTGCCAACG | GSTT1 |
| Gstt1_3 | AAGATATAAATGGCGCGACA | GSTT1 |
| Gstt1_4 | GCTTGCTTACCTCTCACACA | GSTT1 |
| Gstt2_1 | TGGCATGCCGACAACATCCG | GSTT2 |
| Gstt2_2 | CATGAGCGAGCAATTCTCCC | GSTT2 |
| Gstt2_3 | CCCCTTCCAGACGCGTACCG | GSTT2 |
| Gstt2_4 | AAAGTTCCTGTACTCAAAGA | GSTT2 |
| Gstz1_1 | GTCAGAAATCATGCGCACGA | GSTZ1 |
| Gstz1_2 | AAGAGACTCGGCCTATCCCA | GSTZ1 |
| Gstz1_3 | ATTTCTGACCTCATCGCTAG | GSTZ1 |
| Gstz1_4 | TAATGACTCACCGTTAAAGC | GSTZ1 |
| Gys1_1 | TTGGGACACCTGCAACATCG | GYS1 |
| Gys1_2 | CAACGCCCAAAATACACCTG | GYS1 |
| Gys1_3 | TGTTGTGGGTGCACACTGGT | GYS1 |
| Gys1_4 | ATCCCAGATCACCGCAATCG | GYS1 |
| Gys2_1 | GCATAAGAGTAACGTCACCG | GYS2 |
| Gys2_2 | GTGGATGCGATGAATAAACA | GYS2 |
| Gys2_3 | GAGATAACGCCCAAGCAGTG | GYS2 |
| Gys2_4 | AAAACGACAGCCGATGAGTG | GYS2 |
| H6pd_1 | TAGGCGTAGTAATCAGACAG | H6PD |
| H6pd_2 | CTTGAAGGAGACCATAGATG | H6PD |
| H6pd_3 | AGTTCCACGCCCATACCCAG | H6PD |
| H6pd_4 | ATGTCTGCGTAGGCAAAGGG | H6PD |
| Hadh_1 | CCTTTCAACCAGCACCGATG | HADH |
| Hadh_2 | AGCAAATCGGTCTTGTCTGG | HADH |
| Hadh_3 | AAGCATGTGACCGTCATCGG | HADH |
| Hadh_4 | TGGCCATACAGTAGTATTGG | HADH |
| Hdc_1 | GAAGATCAGATTTCTACCTG | HDC |
| Hdc_2 | TTAACTGCTTAGGATTCACG | HDC |
| Hdc_3 | GTTTGTGATTCGGTCCTTCG | HDC |
| Hdc_4 | GACGGAAAACCACCAAACCA | HDC |
| Hk1_1 | CCGACAATCCAAAATAGACG | HK1 |
| Hk1_2 | CGTAGCCGCCATTGAAACGT | HK1 |
| Hk1_3 | GGATCTTTACCAGTAGGACT | HK1 |
| Hk1_4 | CTCCCGGGATTATAACCCAA | HK1 |
| Hk2_1 | ATTCCCGAGGACATCATGCG | HK2 |
| Hk2_2 | GGAGATGCGTCACATTGACA | HK2 |
| Hk2_3 | ATCCGGAGTTGACCTCACAA | HK2 |
| Hk2_4 | GGAGTGGCACACACATAAGT | HK2 |
| Hk3_1 | TGTTACCCACCGGTGCCGTG | HK3 |
| Hk3_2 | ATTCCTGGATGCATACCCCG | HK3 |
| Hk3_3 | TCCTATCCTCAGACTACCTG | HK3 |
| Hk3_4 | ACAATCAGCCCGACTTCACA | HK3 |
| Hmbs_1 | AAGATGAGGGTGATTCGAGT | HMBS |
| Hmbs_2 | CCTGGTCGTTCACTCCCTGA | HMBS |
| Hmbs_3 | AGAGAAGAGCCTGTTTACCA | HMBS |
| Hmbs_4 | CTCAGTTGCTATGTCCACCA | HMBS |
| Hmgcr_1 | TCATCATCCTGACGATAACG | HMGCR |
| Hmgcr_2 | CATTAGGTCGTGGCTCGATG | HMGCR |
| Hmgcr_3 | GCCAAATTGGACGACCCTCA | HMGCR |
| Hmgcr_4 | AACTGCCAGAGAGAAACACT | HMGCR |
| Hmgcs1_1 | GGAAATGCCAGACCTACAGG | HMGCS1 |
| Hmgcs1_2 | TACAACCAATGCATGCTATG | HMGCS1 |
| Hmgcs1_3 | GCTGAGTTGGAAAAATACGA | HMGCS1 |
| Hmgcs1_4 | ACAGTGCTACCTCAGCGCCC | HMGCS1 |
| Hmgcs2_1 | GACATTGCAGTCTACCCGAG | HMGCS2 |
| Hmgcs2_2 | TTTATTACGAACCTTGCTCG | HMGCS2 |
| Hmgcs2_3 | GAGGGATTGTAGAAAACCTG | HMGCS2 |
| Hmgcs2_4 | TCATATACTGCACATCATCG | HMGCS2 |
| Hmox1_1 | TCGTGCTCGAATGAACACTC | HMOX1 |
| Hmox1_2 | TCAGGACCTGACCCCCTGAG | HMOX1 |
| Hmox1_3 | ACGCTTTACATAGTGCTGTG | HMOX1 |
| Hmox1_4 | TTCCTTGTACCATATCTACA | HMOX1 |
| Hmox2_1 | TGCTTATACTCGTTACATGG | HMOX2 |
| Hmox2_2 | GTATGTGAAGTAAAGTGCAG | HMOX2 |
| Hmox2_3 | CCAAGGCATTCATTCTAGCG | HMOX2 |
| Hmox2_4 | CTCCGTGGGGAAATATAAGG | HMOX2 |
| Hs2st1_1 | GTAGGCTATATTGGTGAACG | HS2ST1 |
| Hs2st1_2 | GTCCCGGGCGAAGCTAGGTG | HS2ST1 |
| Hs2st1_3 | AATTTATATCAATGTCATCA | HS2ST1 |
| Hs2st1_4 | TTGTGAAGAACATAACCACC | HS2ST1 |
| Hsd17b10_1 | TGTTAATGATAACTCCACGT | HSD17B10 |
| Hsd17b10_2 | GCTACGGCCAAAAGACTGGT | HSD17B10 |
| Hsd17b10_3 | TTGGTCGCGGTAGTAACTGG | HSD17B10 |
| Hsd17b10_4 | GTCACGACTCACATTTGCTG | HSD17B10 |
| Hsd17b14_1 | TTTCCTGTGTCACGTCACCG | HSD17B14 |
| Hsd17b14_2 | CCAGGTGACCAAATCGAGAG | HSD17B14 |
| Hsd17b14_3 | GTCCCAGGCCCTTACCTACG | HSD17B14 |
| Hsd17b14_4 | GGGGACATACACCTTGATCA | HSD17B14 |
| Hsd17b4_1 | TCTAACATAGGCTCTTCACG | HSD17B4 |
| Hsd17b4_2 | ACCAAACCGTACCAGTCACG | HSD17B4 |
| Hsd17b4_3 | TGGGCGCCATCGTCAGAAAG | HSD17B4 |
| Hsd17b4_4 | TGCAGTGAACGACTTAGGAG | HSD17B4 |
| Hsd3b2_1 | CTTCAGACCAGAAACAAGGA | HSD3B2 |
| Hsd3b2_2 | GATTGTCTTGAATGGCCATG | HSD3B2 |
| Hsd3b2_3 | ATATGTGGGCAATGTAGCCT | HSD3B2 |
| Hsd3b2_4 | ATTATTATGTTAGAAATGAG | HSD3B2 |
| Idh2_1 | GGCCACCCAGAAGTACAGTG | IDH2 |
| Idh2_2 | TCGAGCTGGCACGTTCAAGT | IDH2 |
| Idh2_3 | TCACCGTCCATCTCCACTAC | IDH2 |
| Idh2_4 | ACATCGGCTCATCGACGACA | IDH2 |
| Ido1_1 | TAGGGAACAGCAATATTGCG | IDO1 |
| Ido1_2 | CATACCCAGACAGATATATG | IDO1 |
| Ido1_3 | CTACTGCACTGGATACAGTG | IDO1 |
| Ido1_4 | TCGGGGTCACATACCCATTG | IDO1 |
| Impa2_1 | CCTCGAAGCACTCGTCCCAG | IMPA2 |
| Impa2_2 | AGTTGCAGGTGCCGTCGATG | IMPA2 |
| Impa2_3 | AAACAATTAAATCTTCTACT | IMPA2 |
| Impa2_4 | GGAATACTTTCAGAGTATCG | IMPA2 |
| Impdh1_1 | CTCAGTGTATCAGATCGCCA | IMPDH1 |
| Impdh1_2 | GAAGAACAGAGACTACCCTC | IMPDH1 |
| Impdh1_3 | CCAGGCCAATGAAGTACGGA | IMPDH1 |
| Impdh1_4 | GTAGGTGAGGCCATCCGCGT | IMPDH1 |
| Impdh2_1 | TCCTTCCAGAAATACGAACA | IMPDH2 |
| Impdh2_2 | TTCGTCTTGTAGCTTACAGG | IMPDH2 |
| Impdh2_3 | AGAGCAGACGTCAAGTCCTG | IMPDH2 |
| Impdh2_4 | ATCCTAGATCATGACTAAGA | IMPDH2 |
| Isyna1_1 | ATCGTGAGCTATAACCACCT | ISYNA1 |
| Isyna1_2 | ATCAAACACCAGGTCGTTGG | ISYNA1 |
| Isyna1_3 | TTATGGATCGTTGACCCAGG | ISYNA1 |
| Isyna1_4 | CAATACGGAGCGCTTCTGCG | ISYNA1 |
| Itpr3_1 | CCACGAGGATTGTATCACGG | ITPR3 |
| Itpr3_2 | ACTGCAGATCCCTTACGACA | ITPR3 |
| Itpr3_3 | GCCGTGGAGAAACTCAACGA | ITPR3 |
| Itpr3_4 | GAGGCGGGCAAACTTGACAG | ITPR3 |
| Kcnk5_1 | GGCACAGCGGCACCCCGAAG | KCNK5 |
| Kcnk5_2 | GGAGTGTGGAGAGCTCTCAA | KCNK5 |
| Kcnk5_3 | ACAGCCATCTTCATCGTGTG | KCNK5 |
| Kcnk5_4 | GAAGGCAATGAAAACAAGTG | KCNK5 |
| Kctd5_1 | CCCCGACCTAGATTCGGACA | KCTD5 |
| Kctd5_2 | TATTAACAAAGACCTCGCAG | KCTD5 |
| Kctd5_3 | TGTCTCAAGTAGTTCAGTAC | KCTD5 |
| Kctd5_4 | CCCCGGTGGCGTGTCCAAGT | KCTD5 |
| Kmo_1 | GCTGTCGAAATGCATTCCGG | KMO |
| Kmo_2 | GATGTGTACGAAGCTAGGGA | KMO |
| Kmo_3 | GACAATTCCACCTAAGAATG | KMO |
| Kmo_4 | TTGCCCAAAAAATGGGACAA | KMO |
| Lalba_1 | CAAGCTGTTGTCAACGACAA | LALBA |
| Lalba_2 | GCAACGACAAAATACACACC | LALBA |
| Lalba_3 | ACTGGTATGAAATAAAACAC | LALBA |
| Lalba_4 | AAGTAGTGAGTTCCCCGAGT | LALBA |
| Ldhb_1 | AAATTGTGGCCGATAAAGGT | LDHB |
| Ldhb_2 | GTCTTCCAACACATCCACCA | LDHB |
| Ldhb_3 | GCTCGCCCAGGATCCATCCG | LDHB |
| Ldhb_4 | GGGCTGTACTTGACGATCTG | LDHB |
| Lpcat1_1 | AGGCCATCATGCGCACCATG | LPCAT1 |
| Lpcat1_2 | AGGACACCATCACATGGACG | LPCAT1 |
| Lpcat1_3 | AGTCGGTTACTGAGACACCC | LPCAT1 |
| Lpcat1_4 | CAGTGGAGGAGATCAAGCGA | LPCAT1 |
| Lpcat2_1 | TCATTCCGTGATACCAGCGA | LPCAT2 |
| Lpcat2_2 | TCGGGAGTCAGGGTCAACAC | LPCAT2 |
| Lpcat2_3 | CACGGTAACCGTAAACCCCA | LPCAT2 |
| Lpcat2_4 | CTGTCAACTCTTCACGAAGG | LPCAT2 |
| Lpl_1 | GAAAAACGTACCGTCTGCTG | LPL |
| Lpl_2 | CCATCCATGGATCACCACGA | LPL |
| Lpl_3 | TGGATTCCAATACTTCGACC | LPL |
| Lpl_4 | TGACACTGGATAATGTTGCT | LPL |
| Lta4h_1 | CCTAGTGACTAACAAGACGT | LTA4H |
| Lta4h_2 | GAAATTAACGTACACCGCAG | LTA4H |
| Lta4h_3 | TGGAGGAATAGGCTACATCG | LTA4H |
| Lta4h_4 | TTGGCCCAAGAACTCTGGTG | LTA4H |
| Ltc4s_1 | CTTGCAACAGAACTCCCACG | LTC4S |
| Ltc4s_2 | AGACGCGCTCGAACTCGGGA | LTC4S |
| Ltc4s_3 | CGTATCCCTGGAAATAGCGG | LTC4S |
| Ltc4s_4 | TACAGGTGATCTCTGCACGA | LTC4S |
| Lypla1_1 | TTCACGGATTGGGAGATACA | LYPLA1 |
| Lypla1_2 | CGCGGCGGTGGCCTTCCGGG | LYPLA1 |
| Lypla1_3 | TGGACAGATGTATTTGATGT | LYPLA1 |
| Lypla1_4 | GAGAGCAGTGTATAAAGACA | LYPLA1 |
| Man1a_1 | GGATCCGCGAAAACCACGAG | MAN1A1 |
| Man1a_2 | AGCCTACTATTTGTCCGGAG | MAN1A1 |
| Man1a_3 | AATTTCAAGAAGCTAAATCG | MAN1A1 |
| Man1a_4 | ATAATTATAAACGCTATGCG | MAN1A1 |
| Man1b1_1 | GAGCACCATTCGCATCTTAG | MAN1B1 |
| Man1b1_2 | GGTTACCAAAAGTTTGCTTG | MAN1B1 |
| Man1b1_3 | GTGGACTTCAGACAGCACGG | MAN1B1 |
| Man1b1_4 | ATGTTCATCAACACGAACAG | MAN1B1 |
| Man2a1_1 | TGCGTCGAAATAATCTGACA | MAN2A1 |
| Man2a1_2 | AGGAACCGCGAAAGACTGGG | MAN2A1 |
| Man2a1_3 | CAGCTGGAAATTGTGACCGG | MAN2A1 |
| Man2a1_4 | ACAATCCCTTTGAACAAGAA | MAN2A1 |
| Man2a2_1 | GGCAGTAGTGGTAGACTACG | MAN2A2 |
| Man2a2_2 | AAACGGCAAATCCTCAGACA | MAN2A2 |
| Man2a2_3 | CGGATGGGTGATGCCCGACG | MAN2A2 |
| Man2a2_4 | TTGGCTTCTGTAATAGCCCG | MAN2A2 |
| Man2b1_1 | ATGTTGTAAATGACTACGCG | MAN2B1 |
| Man2b1_2 | TGAGTTCAATGCAAAAACGT | MAN2B1 |
| Man2b1_3 | CTTGATAGTCAATGCGCCCA | MAN2B1 |
| Man2b1_4 | AGCTCCCAGAGGTAACACGT | MAN2B1 |
| Man2b2_1 | CAATGTCTACACTACCGTGG | MAN2B2 |
| Man2b2_2 | TGGTGGCCAACGTTAAACAG | MAN2B2 |
| Man2b2_3 | CAGGCCTTGTGTCTACCGAG | MAN2B2 |
| Man2b2_4 | TCAAGTCATGCACGCCCGCG | MAN2B2 |
| Man2c1_1 | AAGGGGCGGACCAACCACAG | MAN2C1 |
| Man2c1_2 | ATGTTGGCCAAAGAACTTGG | MAN2C1 |
| Man2c1_3 | TGGGTCAAGAGCCAGTACCC | MAN2C1 |
| Man2c1_4 | GCAGATTCTTCATGACGTGG | MAN2C1 |
| Mat1a_1 | ACAGGTATGGTGCTACTGTG | MAT1A |
| Mat1a_2 | CTATGCTACTGATGAGACCG | MAT1A |
| Mat1a_3 | GTGTTGCACAGAGATGACGA | MAT1A |
| Mat1a_4 | GACACATTGGGCAATATCTG | MAT1A |
| Mat2b_1 | TGTGACTGTTATGTTCGACA | MAT2B |
| Mat2b_2 | GCACAATGCACTATGACATG | MAT2B |
| Mat2b_3 | AAGTGAACATCCCTAGCCGG | MAT2B |
| Mat2b_4 | GCTGCTTCCCAGCTGAATGT | MAT2B |
| Mccc2_1 | TTACCAGTTATATGGCGACG | MCCC2 |
| Mccc2_2 | GAGGAGCTTTGATGTCCGAG | MCCC2 |
| Mccc2_3 | TGACTGTGAAAAAGCACGTG | MCCC2 |
| Mccc2_4 | GGTGGTCGTCCAAAGCATAG | MCCC2 |
| Mdh1_1 | GTCAGCGCCATCGATCCCCA | MDH1 |
| Mdh1_2 | GTCCATAGATGTCATTGCAA | MDH1 |
| Mdh1_3 | GACATTCTTTACATCATCAG | MDH1 |
| Mdh1_4 | GTCTTTGGGAAAGACCAGGT | MDH1 |
| Mdh2_1 | TCACGCACCTGAGAGATCAG | MDH2 |
| Mdh2_2 | CGATATCGTAGAGGGTCAGG | MDH2 |
| Mdh2_3 | GTTCGCTCTGACGATGTCAA | MDH2 |
| Mdh2_4 | GTTGGCAATGATGCAAACCA | MDH2 |
| Mgst1_1 | ATTCCAGAGGATAACCAACA | MGST1 |
| Mgst1_2 | GGTAAGAATGATCGTTGCAT | MGST1 |
| Mgst1_3 | GTTTGTTCGCACTGACGAGA | MGST1 |
| Mgst1_4 | TAGAGAGATCTGGTCCACTC | MGST1 |
| Mgst2_1 | TGTGACGGGCGTATATGTAC | MGST2 |
| Mgst2_2 | CTGTATTCATCGTAATGCTG | MGST2 |
| Mgst2_3 | GCTGGGTGGTACTTCAATCA | MGST2 |
| Mgst2_4 | GGTTATTTCGCTTGGCGGGT | MGST2 |
| Mgst3_1 | GGTGGGAGGTGTTTACCACC | MGST3 |
| Mgst3_2 | CGACACTCACGTGTTCTGGT | MGST3 |
| Mgst3_3 | TGTAGTAGCCATATGCGTAA | MGST3 |
| Mgst3_4 | TACAGCACAGATCCTGAGAA | MGST3 |
| Mthfd2_1 | TCGATGAGATATTGTGACTG | MTHFD2 |
| Mthfd2_2 | AGATAATTAAGCGAACAGGT | MTHFD2 |
| Mthfd2_3 | GCTTTCATGTCATTAACGTG | MTHFD2 |
| Mthfd2_4 | CTATGTTCTCAACAAAACCA | MTHFD2 |
| Mthfr_1 | AGACCCTGTAGGTGACCACT | MTHFR |
| Mthfr_2 | GACTGGGATGAGTTTCCTAA | MTHFR |
| Mthfr_3 | AGGTAACATCTACGAAGAGG | MTHFR |
| Mthfr_4 | CGAAGCTCTCTGCATCGGGG | MTHFR |
| Nat1_1 | GAGAATCTTAACATGCATTG | NAT1 |
| Nat1_2 | TGGTCACCTGTACTAGAAGG | NAT1 |
| Nat1_3 | ATTGTGGATTCCGCCTATGG | NAT1 |
| Nat1_4 | AACATATCCTCCCAACATTG | NAT1 |
| Nat2_1 | GTTCCATGGATTCCCCACAA | NAT2 |
| Nat2_2 | CCATCTTTGATCAAATTGTG | NAT2 |
| Nat2_3 | TGGACGTTCCTACCAGATGT | NAT2 |
| Nat2_4 | CCAGCCAATAAGTACAGCAG | NAT2 |
| Ndufa4l2_1 | AGATGGCAGGAACTAGTCTA | NDUFA4L2 |
| Ndufa4l2_2 | GTCTAGGGACCCGCTTCTAC | NDUFA4L2 |
| Ndufa4l2_3 | GGCAGATAAAAAGACACCCT | NDUFA4L2 |
| Ndufa4l2_4 | CCAGACATCAGGACTGCGCA | NDUFA4L2 |
| Nme1_1 | GCAGAAACTTCAGACCAACA | NME1 |
| Nme1_2 | TGGTGGGCGAGATCATCAAG | NME1 |
| Nme1_3 | CTGGTGAAATACATGCACTC | NME1 |
| Nme1_4 | TGGTCTCTCCAAGCATCACG | NME1 |
| Nme2_1 | CGAAGGGCGTTGTTACCCGA | NME2 |
| Nme2_2 | TCAAACGGTTCGAGCAGAAG | NME2 |
| Nme2_3 | CATGAACTCGGGGCCCGTGG | NME2 |
| Nme2_4 | AAGCCAGATGGCGTGCAGCG | NME2 |
| Nmnat1_1 | AAAGGCATTATCTCACCGGT | NMNAT1 |
| Nmnat1_2 | GAACAGCCTGAGGTGCATGT | NMNAT1 |
| Nmnat1_3 | GTTCTGCCATGATGATTCGG | NMNAT1 |
| Nmnat1_4 | GGAGGACATCACGCAAATCG | NMNAT1 |
| Nmnat2_1 | CATGGAAGGTGTGTTGACGT | NMNAT2 |
| Nmnat2_2 | GTGATGCGGTATGAGGAGAT | NMNAT2 |
| Nmnat2_3 | TGAATGTGCCCTTTAGTGAT | NMNAT2 |
| Nmnat2_4 | GACGGTGCCGACTTGACACA | NMNAT2 |
| Nmnat3_1 | GGTGGCTTCCCATCACCGAG | NMNAT3 |
| Nmnat3_2 | GCAGGTGCATATTCGTGATG | NMNAT3 |
| Nmnat3_3 | GACATCTGACTGGATTCGGG | NMNAT3 |
| Nmnat3_4 | CCAGAGTTGAAACTCCTCTG | NMNAT3 |
| Nos1_1 | CTGGACCGCTGGCCAATGTG | NOS1 |
| Nos1_2 | GATGTAGTTGAACATCCCAT | NOS1 |
| Nos1_3 | GCATGTTGGACACAGCGGGG | NOS1 |
| Nos1_4 | GTCAGAAGATGTCCGCACCA | NOS1 |
| Nos2_1 | GAGCCTTTAGACCTCAACAG | NOS2 |
| Nos2_2 | ATCTCCATTCTACTACTACC | NOS2 |
| Nos2_3 | TCTTCACCACAAGGCCACAT | NOS2 |
| Nos2_4 | TGTTTGCCGTCACTCCGCTG | NOS2 |
| Nos3_1 | TGTGTGGGCCGGATCCAGTG | NOS3 |
| Nos3_2 | AACATGCTGCTAGAAATCGG | NOS3 |
| Nos3_3 | GCGAGGGGACCCCGCCAACG | NOS3 |
| Nos3_4 | GCCGCCAAGAGGATACCAGT | NOS3 |
| Pnp_1 | AGATGCTGTGTGATGATGCA | NP |
| Pnp_2 | TCAGTGCCTGGAAACAAATG | NP |
| Pnp_3 | TGTGGCCAGAACCCTCTCCG | NP |
| Pnp_4 | CCTCAAGTGGCAGTGATCTG | NP |
| Npl_1 | TGAACGTCGCCAGGTCGCGG | NPL |
| Npl_2 | CGTGGGAGCACTAAACGTGA | NPL |
| Npl_3 | CACATTGGCCGAAATCCAAG | NPL |
| Npl_4 | ACCATACTTACTCTTTACCC | NPL |
| Nt5c_1 | CTGAAGTACGACCACTGTGT | NT5C |
| Nt5c_2 | ACGAGCAGTACGGAGCTCTG | NT5C |
| Nt5c_3 | GCCTGGAGATTCATACACAC | NT5C |
| Nt5c_4 | GTGCCGCTGGAACAGCGCCG | NT5C |
| Nt5c1a_1 | ACAGCCAAAGCCTCAGAATG | NT5C1A |
| Nt5c1a_2 | CGCCGCGGTACCGCGCTGGG | NT5C1A |
| Nt5c1a_3 | GCTTCGAGTGGCCTTCGATG | NT5C1A |
| Nt5c1a_4 | ATCGGCTGACAAGTAGAGGT | NT5C1A |
| Nt5c1b_1 | AGAACGTCATCCTGACGCCA | NT5C1B |
| Nt5c1b_2 | GATTTCGTACAACGTAACCT | NT5C1B |
| Nt5c1b_3 | GTATACGGGCAGTGATCCGT | NT5C1B |
| Nt5c1b_4 | GACTAATAACCATGCCCAAG | NT5C1B |
| Nt5c2_1 | GTCCTGGAACATGCTACGGT | NT5C2 |
| Nt5c2_2 | ACCGTCGAGAAGCCTATCAC | NT5C2 |
| Nt5c2_3 | GCACGGCATTGTCTACTCAG | NT5C2 |
| Nt5c2_4 | TGAAGTCAAAGAAACGGCAA | NT5C2 |
| Nt5c3_1 | AGAGTAGAAGAAATTATCTG | Nt5c3 |
| Nt5c3_2 | CTCACCACTCTACCATGTAA | Nt5c3 |
| Nt5c3_3 | CCAGCAGAAAATATGAACAC | Nt5c3 |
| Nt5c3_4 | TCTCACTTACTATGACACGT | Nt5c3 |
| Nt5c3b_1 | CCATGTAGTTGGACACAATG | NT5C3B |
| Nt5c3b_2 | CCATTGAGATCGACCCACAC | NT5C3B |
| Nt5c3b_3 | ATTGTAAGCAAATCTGCTCA | NT5C3B |
| Nt5c3b_4 | ATTGTGGGCGCCCTTCGCAG | NT5C3B |
| Nt5dc1_1 | AACCGGGCTGGTACTCCCAA | NT5DC1 |
| Nt5dc1_2 | CCTTTGGCAAGAAGGAGTGG | NT5DC1 |
| Nt5dc1_3 | TTCTTACAGACTCAGGACAG | NT5DC1 |
| Nt5dc1_4 | AGTGGTGGACTCTTTAACAA | NT5DC1 |
| Nt5dc2_1 | CAGAAAGTAGTCCACCACAC | NT5DC2 |
| Nt5dc2_2 | GAAGTGATCGACCTGTACGG | NT5DC2 |
| Nt5dc2_3 | CAGTGGATCGAGCAAGACAT | NT5DC2 |
| Nt5dc2_4 | GGCACATGGTAGGTCCCGAT | NT5DC2 |
| Nt5dc3_1 | AATGTACCGAGCAATCGAAG | NT5DC3 |
| Nt5dc3_2 | CAAAGGGATGCGGTACATCG | NT5DC3 |
| Nt5dc3_3 | GTGCTTGGAATAGAATACCA | NT5DC3 |
| Nt5dc3_4 | GAAGTCATAGACATGTATGA | NT5DC3 |
| Nt5e_1 | TCATGAATTTGATAACGGTG | NT5E |
| Nt5e_2 | TGAATAAGATCATCGCCCTG | NT5E |
| Nt5e_3 | TATGCCTTTGGCAAATACCT | NT5E |
| Nt5e_4 | CCTGAAGCGGCACGTCTGAG | NT5E |
| Nt5m_1 | GGGTGTCGGAGCAGTACGGA | NT5M |
| Nt5m_2 | AGTACTTGAACATCTTGATG | NT5M |
| Nt5m_3 | GTGCCCCGCTACGGGCCGAG | NT5M |
| Nt5m_4 | GTTACTCCAGTATGCCTGGG | NT5M |
| Nudt1_1 | GAGCGTGGATACACTGCACA | NUDT1 |
| Nudt1_2 | AGGCTGTAGCACTAGCACAA | NUDT1 |
| Nudt1_3 | GGGACGCCCACAGAAAGTGA | NUDT1 |
| Nudt1_4 | AAGGAGAGACCATTGAGGAT | NUDT1 |
| Nudt21_1 | ACTAAAACGCTTAATGACAG | NUDT21 |
| Nudt21_2 | CCGAACTGGTTGACCCCCCG | NUDT21 |
| Nudt21_3 | AGCCAGATTTCAGCGCATGA | NUDT21 |
| Nudt21_4 | CCTGGTGGGGAACTTAACCC | NUDT21 |
| Nudt5_1 | AAACAGTGAAACTTACAACC | NUDT5 |
| Nudt5_2 | ACAACTTATATGGATCCCAC | NUDT5 |
| Nudt5_3 | CAAGCACCTTGTTACCTCAG | NUDT5 |
| Nudt5_4 | CCTCCAGGGTTCATCGAAGA | NUDT5 |
| Oas2_1 | AGAGTCGTAACTCTCCAGCG | OAS2 |
| Oas2_2 | ACTGACCCAACCAATAATGT | OAS2 |
| Oas2_3 | GAAGACTTCGACATGGCACA | OAS2 |
| Oas2_4 | TCAAGTAAGAGCCTCACCAG | OAS2 |
| Odc1_1 | ATATTGACGTCATTGGTGTG | ODC1 |
| Odc1_2 | CTGCGTAAAAGGGAGTGACG | ODC1 |
| Odc1_3 | GCAATCCGTAGAACCAACCT | ODC1 |
| Odc1_4 | TCATTGCCAAAAAAACCGTG | ODC1 |
| Ogdh_1 | TTGGCCCACTCATAGATACG | OGDH |
| Ogdh_2 | GACTAGTTCGAACTATGTGG | OGDH |
| Ogdh_3 | GTAAGTGGAAGACCTTGTCA | OGDH |
| Ogdh_4 | AAAGCTGAACAGTTCTACTG | OGDH |
| Pah_1 | CCTCTTCTGGAAAAGTACTG | PAH |
| Pah_2 | CACTTACCTCAAATAAGCGC | PAH |
| Pah_3 | GCAGCATCATCAAGAGCCTG | PAH |
| Pah_4 | TCCTCGGGTGGAATACACAG | PAH |
| Paics_1 | TTCGGCACACCCACTCAATG | PAICS |
| Paics_2 | ATGCCAATAATGATCCACAG | PAICS |
| Paics_3 | TCAACCAGCGTACAGTCCTG | PAICS |
| Paics_4 | GTGGGTTGCAGACCGAGTGG | PAICS |
| Pank1_1 | ACACTGCCTACGGCAAAACT | PANK1 |
| Pank1_2 | AAAGTAAACCAGCTTAACCA | PANK1 |
| Pank1_3 | CCATGTTAACCAGCAACATA | PANK1 |
| Pank1_4 | GGACAACTACAAAAGAGTGA | PANK1 |
| Pank2_1 | GTCCGCAGACTCGCCGACCG | PANK2 |
| Pank2_2 | GGATCCGTAAGCCACATTGG | PANK2 |
| Pank2_3 | AAAGCAGGCATGTCATGGGT | PANK2 |
| Pank2_4 | CGAGTAGCGCGGCGCCGTCG | PANK2 |
| Pank3_1 | CTTTATCAGATTTCCAACCC | PANK3 |
| Pank3_2 | CCAATATTCACTACCAGCAG | PANK3 |
| Pank3_3 | GCTCTGGAAATGGCATCCAA | PANK3 |
| Pank3_4 | ACCTATTGACATCACAGCAG | PANK3 |
| Pck1_1 | ACTGACAGACTCGCCCTATG | PCK1 |
| Pck1_2 | GTGGCCGAGACTAGCGATGG | PCK1 |
| Pck1_3 | CCTTTGGAAGCGGATATGGT | PCK1 |
| Pck1_4 | TCGCAGATGTGGATATACTC | PCK1 |
| Pck2_1 | TGCGTATTATGACCCGCCTG | PCK2 |
| Pck2_2 | TGATTGTAACTCCTTCGCAG | PCK2 |
| Pck2_3 | AGGGTTTGGATGCTACGGCA | PCK2 |
| Pck2_4 | ATGGAAGCACATACATAATG | PCK2 |
| Pcyt1a_1 | AATACGTATCTAATTGTGGG | PCYT1A |
| Pcyt1a_2 | ATGCAAGACGGAACCTACAG | PCYT1A |
| Pcyt1a_3 | GCACCACCTCGTCCACGTAG | PCYT1A |
| Pcyt1a_4 | CTTTAGTAAGCCCTATGTCA | PCYT1A |
| Pcyt1b_1 | GTGCCCTTGCGTGACCTGAG | PCYT1B |
| Pcyt1b_2 | CCTCACCTGGTGTTCCTAAG | PCYT1B |
| Pcyt1b_3 | GCTGGATTCATCAGCAAATG | PCYT1B |
| Pcyt1b_4 | GCCTCCCTCAGAAACCATGG | PCYT1B |
| Pcyt2_1 | CCGTATGCACACCCACGATG | PCYT2 |
| Pcyt2_2 | GAGGATACGGAACAGGTCAA | PCYT2 |
| Pcyt2_3 | AGACGGCCGAGACACCTATG | PCYT2 |
| Pcyt2_4 | GTGTAGGCCGGCGATGACGT | PCYT2 |
| Pde9a_1 | TAAGTGGGGACATCACGTCG | PDE9A |
| Pde9a_2 | ATACCAGATCAATGCCCGCA | PDE9A |
| Pde9a_3 | GACACAGATGATGTACAGTA | PDE9A |
| Pde9a_4 | CTGGCTTTGGGAACCCAACG | PDE9A |
| Pdha1_1 | AGAACAACCGCTATGGCATG | PDHA1 |
| Pdha1_2 | ATCACTGCCTATCGAGCACA | PDHA1 |
| Pdha1_3 | GCGCCGGATGGAGCTAAAGG | PDHA1 |
| Pdha1_4 | TGTTTGACATTATACGGCGA | PDHA1 |
| Pdha2_1 | TTCTCACAGATGAAAACACA | PDHA2 |
| Pdha2_2 | CAAGTACTACCGGACCATGC | PDHA2 |
| Pdha2_3 | GGACGTGATGACGTGATCCG | PDHA2 |
| Pdha2_4 | TACGGCAAGAACTTCTACGG | PDHA2 |
| Pdhb_1 | TCTTAATTGTAGGTTAGCAG | PDHB |
| Pdhb_2 | GGGCACAGGCTGAAGGCCAG | PDHB |
| Pdhb_3 | TGAAGCTATTAATCAAGGTA | PDHB |
| Pdhb_4 | GCCATTCGTGATAATAACCC | PDHB |
| Pdhx_1 | CTTTAGTGAAGATCCCGCGA | PDHX |
| Pdhx_2 | GTTAGACGAGATCTGGTCAA | PDHX |
| Pdhx_3 | TGTCTCCTACGATGGAGCAA | PDHX |
| Pdhx_4 | TCCTGGGCAACCGAATGCAG | PDHX |
| Pdk1_1 | TTGTCGCAGAAACATAAACG | PDK1 |
| Pdk1_2 | ATGGCTATGAGAACGCTAGG | PDK1 |
| Pdk1_3 | TTGATAGCCTTATTGTTCGG | PDK1 |
| Pdk1_4 | AAACACCATGTGATAGAGAT | PDK1 |
| Pdk2_1 | CAACTGCAGCGTGTCTGATG | PDK2 |
| Pdk2_2 | CATAGACCATGTGAATGGGC | PDK2 |
| Pdk2_3 | GCAGTTTCTAGACTTCGGTA | PDK2 |
| Pdk2_4 | GGTGCTGCCATCAAAGATGA | PDK2 |
| Pdk3_1 | CTTACTTATCCAAAACTCGT | PDK3 |
| Pdk3_2 | CCAGCTCTGAACTAATCCCA | PDK3 |
| Pdk3_3 | TAAGCATGCGGAAAGAAATG | PDK3 |
| Pdk3_4 | GTTGTTCCTACAATGGCCCA | PDK3 |
| Pdk4_1 | TATCGACCCAAACTGTGATG | PDK4 |
| Pdk4_2 | GCTTAGGTTTGTAGACACGC | PDK4 |
| Pdk4_3 | TGAGCATCCGAGTAGAAATG | PDK4 |
| Pdk4_4 | GGAACGTACACAATGTGGAT | PDK4 |
| Pdp1_1 | GAGGTCAACACCATCTACAT | PDP1 |
| Pdp1_2 | GTGTACCTCAGACGATTCTG | PDP1 |
| Pdp1_3 | ACTGGAACACCCAAAAAATG | PDP1 |
| Pdp1_4 | TGCTTGTGCCACTGAAGGAT | PDP1 |
| Pdp2_1 | CCTTATAAATACTATCTCCG | PDP2 |
| Pdp2_2 | CTACAGACACACATCAACCG | PDP2 |
| Pdp2_3 | CTTCAGCAGGTTCCTACTCG | PDP2 |
| Pdp2_4 | GCAGGTGAACTCCATTGACG | PDP2 |
| Pdpr_1 | CCAGTCTGGAAATCCCAACG | PDPR |
| Pdpr_2 | GCATGCCAGTATGAACGATG | PDPR |
| Pdpr_3 | AGTGATCCCAGTCTTCACGA | PDPR |
| Pdpr_4 | GGCGAGTCTCCTGTAGTACA | PDPR |
| Pdss1_1 | AGCCTTTAGACCGATTATTG | PDSS1 |
| Pdss1_2 | GAGGCGGCACGTGTTCCGAG | PDSS1 |
| Pdss1_3 | TATACTATGCATAGTTGGGG | PDSS1 |
| Pdss1_4 | GTTATTGAAGATTTGGTGCG | PDSS1 |
| Pdxk_1 | AGGGGTACTAACCATACACA | PDXK |
| Pdxk_2 | CTCTGTTACCCACGTAGCCA | PDXK |
| Pdxk_3 | CCACTTTGTCCCTGTACACG | PDXK |
| Pdxk_4 | GGAACTTCATGAGCTGTACG | PDXK |
| Pfkl_1 | GGAGATACCAGTGCGCACTG | PFKL |
| Pfkl_2 | CTGTAAGGCCTTCACTACGA | PFKL |
| Pfkl_3 | AGCCCTGCACCGCATTATGG | PFKL |
| Pfkl_4 | CCGTATGGGCATATATGTGG | PFKL |
| Pfkm_1 | CCTCACGGTAGAGCGAACAG | PFKM |
| Pfkm_2 | GCGCCTTGGATATGACACCC | PFKM |
| Pfkm_3 | TTAGACCAAAGACGTGACCA | PFKM |
| Pfkm_4 | CATAGACACGCTCTCCCACG | PFKM |
| Pfkp_1 | CAATCTGTGCGTGATCGGCG | PFKP |
| Pfkp_2 | GAAGTATTCCTACCTCAACG | PFKP |
| Pfkp_3 | TTAGATCAAAGAAATCGGCT | PFKP |
| Pfkp_4 | AAGGAAGCCGTGAAACTCCG | PFKP |
| Pfkfb2_1 | TGATGCCACCAATACCACTC | Pfkfb2 |
| Pfkfb2_2 | TGGGTACTCCAATCCAGTTG | Pfkfb2 |
| Pfkfb2_3 | TTTAAAAAAAAACTCACTTG | Pfkfb2 |
| Pfkfb2_4 | CGAGCCCTGACTACCCCGAA | Pfkfb2 |
| Pfkfb3_1 | TGTGGGAGAGTATCGGCGTG | Pfkfb3 |
| Pfkfb3_2 | AAGTACCATTCAGACCGCGG | Pfkfb3 |
| Pfkfb3_3 | TGTCAAAAGCTACCTGACAA | Pfkfb3 |
| Pfkfb3_4 | CATTCTCGCCATGCCGACAC | Pfkfb3 |
| Pfkfb4_1 | CAACCAATACGACCCGAGAG | Pfkfb4 |
| Pfkfb4_2 | TCCCCCCCAGAATTCAACGT | Pfkfb4 |
| Pfkfb4_3 | CCCCGGTCTTACCACCACGC | Pfkfb4 |
| Pfkfb4_4 | TGTCCCGGTTGACATAGTCA | Pfkfb4 |
| Pgk1_1 | TAAGGTGCTCAACAACATGG | PGK1 |
| Pgk1_2 | TCAAGAACAGAACATCCCTG | PGK1 |
| Pgk1_3 | GGACTGCACACCGAGCCCAT | PGK1 |
| Pgk1_4 | CTTCCTCTACATGAAAGCGG | PGK1 |
| Phgdh_1 | CCACACTGAGAGCCTACCTG | PHGDH |
| Phgdh_2 | CTCACTTCTGACCAGACTGT | PHGDH |
| Phgdh_3 | TAGTAGCAGACCGGACAATG | PHGDH |
| Phgdh_4 | CTGCGATTTCCTCCCCACAG | PHGDH |
| Pik3c3_1 | AGCCTGTAAGAACTCAACAC | PIK3C3 |
| Pik3c3_2 | ATACACATCCCATATAGTCA | PIK3C3 |
| Pik3c3_3 | CTCACCAAGGCTCATCGGCA | PIK3C3 |
| Pik3c3_4 | ATGGACCAGGCGATCTACAA | PIK3C3 |
| Pip4k2a_1 | GGAAATACGACTTAAAGGTG | PIP4K2A |
| Pip4k2a_2 | GTATATAGTGGAATGTCATG | PIP4K2A |
| Pip4k2a_3 | GGGGGGTCAACCATTCGGTA | PIP4K2A |
| Pip4k2a_4 | CATCAAGACCATTACCAGTG | PIP4K2A |
| Pip4k2b_1 | CAGTCTTGATGACGAAACGC | PIP4K2B |
| Pip4k2b_2 | GCATCGCAAATACGACCTGA | PIP4K2B |
| Pip4k2b_3 | AGCTTACAGCAAGATCAAGG | PIP4K2B |
| Pip4k2b_4 | GGGCATGTACCGCCTGACCG | PIP4K2B |
| Pip4k2c_1 | TTTCGTACAGCAGAAAGTGA | PIP4K2C |
| Pip4k2c_2 | AGTGTCGAGCGAAGATATTG | PIP4K2C |
| Pip4k2c_3 | TCTGGCAGCAACATCACTGG | PIP4K2C |
| Pip4k2c_4 | TCCTTGAACTTGAAATGACT | PIP4K2C |
| Pip5k1a_1 | GTGAGTGATGCCTAACTGGA | PIP5K1A |
| Pip5k1a_2 | ACAACATGGACCATGCACAG | PIP5K1A |
| Pip5k1a_3 | TGAAGGGTTCAACTTACAAG | PIP5K1A |
| Pip5k1a_4 | ACAGCTATGGAATCCATCCA | PIP5K1A |
| Pip5k1c_1 | TGGTGGCAAGAACATCCGCG | PIP5K1C |
| Pip5k1c_2 | CCGACGCATCCACGCCTCGG | PIP5K1C |
| Pip5k1c_3 | TTACCAAATAGTCATCTGGA | PIP5K1C |
| Pip5k1c_4 | GCCATGGAGTCTATCCAGGG | PIP5K1C |
| Pipox_1 | TCCAAACATTCGGTTCACCA | PIPOX |
| Pipox_2 | ACAGAGCCGCATAATCAGAA | PIPOX |
| Pipox_3 | AAGCCGGAACCCAGTTACAC | PIPOX |
| Pipox_4 | GGCTTGGTAGCTCTTCAAGG | PIPOX |
| Pklr_1 | TGTACGAAAAGCCAGTGATG | PKLR |
| Pklr_2 | GGGTTCACTCCAGACCTGTG | PKLR |
| Pklr_3 | GGGCGATGCAAAGACAGTGT | PKLR |
| Pklr_4 | CTGGTGACCGAAGTGGAACA | PKLR |
| Pkm_1 | TTTCTCTCATGGAACCCATG | PKM |
| Pkm_2 | TGAAATAGCACATGCCTGTG | PKM |
| Pkm_3 | GGGCAGAGTCAATGTCCAGG | PKM |
| Pkm_4 | CTTCCTGACTTCATGCACGT | PKM |
| Pla2g10_1 | AGGTCACATGTATACAAGCG | PLA2G10 |
| Pla2g10_2 | CCATAGTTCATGTAAGCCAT | PLA2G10 |
| Pla2g10_3 | TCGAGGCCCAACACAATCCA | PLA2G10 |
| Pla2g10_4 | CACCAGTCAATGGCGTCACG | PLA2G10 |
| Pla2g12a_1 | TGCATCTCACAGCTGAACAT | PLA2G12A |
| Pla2g12a_2 | GATGGTTTATATCCATAGCG | PLA2G12A |
| Pla2g12a_3 | TGAGGTACGTGTCTATCTTG | PLA2G12A |
| Pla2g12a_4 | AACAGGACCAGACCACCGAC | PLA2G12A |
| Pla2g12b_1 | CGAGTCTGTCAACAGCTACG | PLA2G12B |
| Pla2g12b_2 | TCCACTCACCATATCGACAC | PLA2G12B |
| Pla2g12b_3 | TTCCTGGTACCTTGATACCC | PLA2G12B |
| Pla2g12b_4 | GACGTCTGCTACGACACCTG | PLA2G12B |
| Pla2g1b_1 | CCTTCAGGGGATCACTCCCG | PLA2G1B |
| Pla2g1b_2 | CCTGTCTAAGTCGTCCACTG | PLA2G1B |
| Pla2g1b_3 | GCATGAGTAGGAGTAAGTGT | PLA2G1B |
| Pla2g1b_4 | TGGTGCACTTGATCATATTG | PLA2G1B |
| Pla2g2d_1 | AGATGGTCACACACATGACG | PLA2G2D |
| Pla2g2d_2 | GGAGGGGATTTACCAGTCTG | PLA2G2D |
| Pla2g2d_3 | GGTATAACTGCAACCCAGGG | PLA2G2D |
| Pla2g2d_4 | TGGCCCTACGGCTGTCACTG | PLA2G2D |
| Pla2g2e_1 | CCACTGGCCAGTGGACGAGA | PLA2G2E |
| Pla2g2e_2 | GCCATAGTCATTGTACTGCA | PLA2G2E |
| Pla2g2e_3 | TGTCTCGAGTGATAGAGAAG | PLA2G2E |
| Pla2g2e_4 | TATGGCTGCTATTGCGGTGT | PLA2G2E |
| Pla2g2f_1 | GCTTCTCATAGCAACAGTCG | PLA2G2F |
| Pla2g2f_2 | GGTCATAGTGGTCCACGTAG | PLA2G2F |
| Pla2g2f_3 | CCTTCTTGCAGTGGTAACCA | PLA2G2F |
| Pla2g2f_4 | TCCCTCTAAAACCTCCAGGT | PLA2G2F |
| Pla2g3_1 | CTGACAGAGAGCACCGTCGA | PLA2G3 |
| Pla2g3_2 | GATGCCATAGTTATACTGCA | PLA2G3 |
| Pla2g3_3 | GCCAGTGGGAGGCATGTCAA | PLA2G3 |
| Pla2g3_4 | GGGAGCCCAGAGGGTTACCA | PLA2G3 |
| Pla2g4b_1 | CTCCGAGCCAAAGAGCTCGG | PLA2G4B |
| Pla2g4b_2 | GTACCGCCAAAGTTGGCTAG | PLA2G4B |
| Pla2g4b_3 | GGGCACAATAGATAGGCAGA | PLA2G4B |
| Pla2g4b_4 | ACTGCATCTCCTATATCACG | PLA2G4B |
| Pla2g5_1 | CCGTTGTTATGGGCAACTGG | PLA2G5 |
| Pla2g5_2 | GGTCGCGGGACACCCAAGGA | PLA2G5 |
| Pla2g5_3 | GTCATAGGACTGGGTCCGAA | PLA2G5 |
| Pla2g5_4 | TGAGTTCTAGCAAGCCCCCT | PLA2G5 |
| Pla2g6_1 | AAATCCATGGCCTATATGCG | PLA2G6 |
| Pla2g6_2 | GGCACCTGCATTAGCCCCGT | PLA2G6 |
| Pla2g6_3 | AGTCTCCCCAAAGTCGTTTG | PLA2G6 |
| Pla2g6_4 | CCATTGGGCCAAGAACGCCG | PLA2G6 |
| Pla2g7_1 | GGAAATGGATCGTACCCCGT | PLA2G7 |
| Pla2g7_2 | TTGGGATCCAAACAGTGTCG | PLA2G7 |
| Pla2g7_3 | TAGTGGCCACTGTCGAACAC | PLA2G7 |
| Pla2g7_4 | GCGATTCTTGACATTGAACA | PLA2G7 |
| Pnpo_1 | AATGCGCAAGAGTTACCGCG | PNPO |
| Pnpo_2 | ACTAACTACGAGAGCCGGAA | PNPO |
| Pnpo_3 | GGTAGCCACACACATAGCAT | PNPO |
| Pnpo_4 | CTGAACAGCCTCGTCAAACC | PNPO |
| Pon2_1 | AGGTGAAATATTGATCCCGT | PON2 |
| Pon2_2 | GACATACTTGAACCTACACT | PON2 |
| Pon2_3 | GCGGTTCCTAGCGCTCAGGT | PON2 |
| Pon2_4 | ATCTCAAAGACGAGAGACCG | PON2 |
| Ppap2a_1 | AAAGTATATCCATTTCAGAG | Plpp1 |
| Ppap2a_2 | GTATGGCGAGTATACGGGAC | Plpp1 |
| Ppap2a_3 | CCCGAGTAGAAAGACAACCT | Plpp1 |
| Ppap2a_4 | ATACTTAGCGATGTCAGTCA | Plpp1 |
| Ppap2c_1 | CATGCAATACATGCCAAAGG | Plpp2 |
| Ppap2c_2 | AGTTGGATCGCGAATAAAGA | Plpp2 |
| Ppap2c_3 | CAGCCCTGCTAATGTCACGG | Plpp2 |
| Ppap2c_4 | CATAGCCAGAACAGTTGACC | Plpp2 |
| Ppap2b_1 | CATTGCAGTAAAACCCTCGA | Plpp3 |
| Ppap2b_2 | GCAGCTCTCTATAAGCAAGT | Plpp3 |
| Ppap2b_3 | GTCCCTGAGAGTAAGAACGG | Plpp3 |
| Ppap2b_4 | CAGTCAGATCAATTGCTCCG | Plpp3 |
| Ppat_1 | ACCTTGGAATCGGACATACG | PPAT |
| Ppat_2 | ATAAGACGCCCGATGCAGAG | PPAT |
| Ppat_3 | TGATCACTCTGGGACTCGTG | PPAT |
| Ppat_4 | AGGGGTGTATGCGAGTAACT | PPAT |
| Prdx4_1 | ATCTAAGCAAAGCCAAGAGT | PRDX4 |
| Prdx4_2 | GTATTACTTACAAATCCAGT | PRDX4 |
| Prdx4_3 | TGGACAAGTGTACCCCGGGG | PRDX4 |
| Prdx4_4 | AATACCCCTCGAAGACAAGG | PRDX4 |
| Prps1_1 | GATGACTGCAGTAACCCGGC | PRPS1 |
| Prps1_2 | TGCAGATCATATTATCACCA | PRPS1 |
| Prps1_3 | CATCTCCCACAAGTACCATG | PRPS1 |
| Prps1_4 | ATGTCTACATTGTTCAAAGT | PRPS1 |
| Prps2_1 | GATCGGTGAAAGTGTGAGAG | PRPS2 |
| Prps2_2 | GTGGACCGGATGGTTCTCGT | PRPS2 |
| Prps2_3 | GGAGATTAATGACAACCTGA | PRPS2 |
| Prps2_4 | GGCTGATCACATCATTACCA | PRPS2 |
| Psat1_1 | TGGAAGGAGTGCTGACTACG | Psat1 |
| Psat1_2 | TGCAAACGAGACTGTGCACG | Psat1 |
| Psat1_3 | CAATACAGAGAATCTTGTGA | Psat1 |
| Psat1_4 | CTTTGTAGTCAAGGACTGAT | Psat1 |
| Psph_1 | TTCGTACATTTCAGACACTG | PSPH |
| Psph_2 | CTTATGCCAGGAGTCAGATG | PSPH |
| Psph_3 | CCTGGACATTACGCTCCTGG | PSPH |
| Psph_4 | CAGCCTATTGGCAAACACAT | PSPH |
| Ptgs1_1 | TGGCACGGATAGTAACAACA | PTGS1 |
| Ptgs1_2 | AGCTGCTCATCATCCCACGT | PTGS1 |
| Ptgs1_3 | TGGTGGGTAGCGCATCAACA | PTGS1 |
| Ptgs1_4 | GTGGATGCCAGTGATAGAGA | PTGS1 |
| Ptgs2_1 | AACATCATATTTGAGCCTTG | PTGS2 |
| Ptgs2_2 | TAGTGCACATTGTAAGTAGG | PTGS2 |
| Ptgs2_3 | GACTACGTGCAACACCTGAG | PTGS2 |
| Ptgs2_4 | CGTGGGGAATGTATGAGCAC | PTGS2 |
| Pts_1 | GGGAAATGCAACAATCCGAA | PTS |
| Pts_2 | CGAGGCGCGACAGTCGCGCG | PTS |
| Pts_3 | CGGGCACAACTATAAAGGTG | PTS |
| Pts_4 | GTGATCAAGAGGCTTCATGA | PTS |
| Pycr1_1 | CCGGGGTGTTAGTCATACAT | PYCR1 |
| Pycr1_2 | TTGGGGCGAACATTGAGGAC | PYCR1 |
| Pycr1_3 | CACAGGTACTCACGCTCAGG | PYCR1 |
| Pycr1_4 | GGAGGGCCGAGACCGTAGCT | PYCR1 |
| Pygb_1 | CCTCTTAACCCAGAGAATCG | PYGB |
| Pygb_2 | AGAGCGAAGAAGTAGTCGCG | PYGB |
| Pygb_3 | TCTTTCCTTAGTCAACGTTG | PYGB |
| Pygb_4 | GGACTGCAAGAGAATCAACA | PYGB |
| Pygl_1 | AAACCAACGGGATTACCCCG | PYGL |
| Pygl_2 | GGGCGAAGTAGTAGTCGCGG | PYGL |
| Pygl_3 | CTTACAAAATGCCTGCGATG | PYGL |
| Pygl_4 | AGGAGGCAAACGGATCAACA | PYGL |
| Pygm_1 | GTAGCCGCCAACATTGACTG | PYGM |
| Pygm_2 | ACTTGGAGGACTTGAAACGT | PYGM |
| Pygm_3 | GGATCCAGCGTCCCACGAGG | PYGM |
| Pygm_4 | GCAGCCTATCTACGTCCCCA | PYGM |
| Rrm1_1 | AACTACATAAATCCGCACAA | RRM1 |
| Rrm1_2 | CCCAGCGATGTAGCTACCAG | RRM1 |
| Rrm1_3 | AATCCAAGGCCTATATAGTG | RRM1 |
| Rrm1_4 | AAACTACTACCCTATTCCAG | RRM1 |
| Rrm2_1 | CTTGCTAGGAGAACGCGTTG | RRM2 |
| Rrm2_2 | TATTCTGGCTCAAGAAACGG | RRM2 |
| Rrm2_3 | GGAAAGACAACGAAGCGGCG | RRM2 |
| Rrm2_4 | ACATTAAAGATCCCAAGGAA | RRM2 |
| Rrm2b_1 | GAGAATGTACAAGCAAGCAC | RRM2B |
| Rrm2b_2 | TTGGAAAGATGACGAACCGT | RRM2B |
| Rrm2b_3 | GGAGATAAAATACTTCTCGT | RRM2B |
| Rrm2b_4 | TCTCCTAGGGGAAAGAGTGG | RRM2B |
| Sat1_1 | ATCCTGCGACTGATCAAGGT | SAT1 |
| Sat1_2 | CTGTCTTACAGATCTCCAAG | SAT1 |
| Sat1_3 | ACAGCAACTTGCCAATCCAT | SAT1 |
| Sat1_4 | GGATCAGGGGTTACCTTCAG | SAT1 |
| Sat2_1 | AGAGATTATCCCAGCACCTG | SAT2 |
| Sat2_2 | CAGCCAGTACCCCGATATTG | SAT2 |
| Sat2_3 | TTCATCTATAGCACATGGAC | SAT2 |
| Sat2_4 | AGACATTATGAGGATGATCC | SAT2 |
| Scd1_1 | ATGATAAGGAAGATCCGCAG | Scd1 |
| Scd1_2 | AGGGGCGCTGCTCACCGAAG | Scd1 |
| Scd1_3 | TCTCGTTCATTTCCGGAGGG | Scd1 |
| Scd1_4 | GGATGAAGCACATCAGCAGG | Scd1 |
| Sephs1_1 | ATGGTGTACATACCAAGTCG | SEPHS1 |
| Sephs1_2 | CACCCAAGAAGACGTAGAGT | SEPHS1 |
| Sephs1_3 | ATATACGGGTCGTCGACGAT | SEPHS1 |
| Sephs1_4 | CCTCAGTGACCTTTATGCAA | SEPHS1 |
| Shmt2_1 | CGGCAGATACTACGGAGGAG | SHMT2 |
| Shmt2_2 | AACATCCGCGTACTTGAAAG | SHMT2 |
| Shmt2_3 | AGCCTCATGATCGAATCATG | SHMT2 |
| Shmt2_4 | TAGTCGATGAGGCCAGTTTG | SHMT2 |
| Slc11a1_1 | GGGTGTGCTACCACATACTG | SLC11A1 |
| Slc11a1_2 | GAGAAGTAGACAGAACCCGC | SLC11A1 |
| Slc11a1_3 | CCTAGCATGATACCGTCCAG | SLC11A1 |
| Slc11a1_4 | GAATGGGGATCTTCTCACTC | SLC11A1 |
| Slc12a7_1 | AATGAAAGATGTAGTCACGA | SLC12A7 |
| Slc12a7_2 | AACAACGTTACTGAGATACA | SLC12A7 |
| Slc12a7_3 | TGGTCATGGAAAGGCCAACG | SLC12A7 |
| Slc12a7_4 | AGCAGATGAACATCAGCGCG | SLC12A7 |
| Slc12a8_1 | CATCTGCCCACCAAGCACCG | SLC12A8 |
| Slc12a8_2 | CAGTGAAGAATGACTCCCCG | SLC12A8 |
| Slc12a8_3 | GAAAGGGCCAAATAAAACAC | SLC12A8 |
| Slc12a8_4 | CCAGTTCCTCTATTGGCTGG | SLC12A8 |
| Slc15a1_1 | TTGTCCAATCGTGTAGACGA | SLC15A1 |
| Slc15a1_2 | GAAGTAAGGCATATCCCAAG | SLC15A1 |
| Slc15a1_3 | CCACCAAACGCAGACACACA | SLC15A1 |
| Slc15a1_4 | CTGGGACGACAATCTCTCCA | SLC15A1 |
| Slc16a1_1 | ACTACTAAGAAAGACCAAAG | SLC16A1 |
| Slc16a1_2 | CACCAGCGATCATTACTGGA | SLC16A1 |
| Slc16a1_3 | GACTTGCAGCCAACACCAAG | SLC16A1 |
| Slc16a1_4 | AGGCCCTATTGGTCTCATCA | SLC16A1 |
| Slc16a3_1 | TATGGGTGTACCCGACACAA | SLC16A3 |
| Slc16a3_2 | CTGAGTGTCTTCCGAGACCG | SLC16A3 |
| Slc16a3_3 | AGTATCGATTGAGCATGATG | SLC16A3 |
| Slc16a3_4 | AAAAGACGCTGACCGCCTTG | SLC16A3 |
| Slc25a1_1 | ATGAACGAGCGAACCCACCG | SLC25A1 |
| Slc25a1_2 | GAATAATCTCTCTAACCCCG | SLC25A1 |
| Slc25a1_3 | ACTGCGACTGTACTGAAGCA | SLC25A1 |
| Slc25a1_4 | CTTCACGTATTCGGTCGGGA | SLC25A1 |
| Slc25a13_1 | GGGGCGACTCCCAGTAACTG | SLC25A13 |
| Slc25a13_2 | CAGATTTATATGAGCCGAGG | SLC25A13 |
| Slc25a13_3 | ACAAGGCATCCGGAGCACAC | SLC25A13 |
| Slc25a13_4 | TACAAGATCGATAGGATACA | SLC25A13 |
| Slc25a36_1 | TTGTAGATGTGGTGGCACAG | SLC25A36 |
| Slc25a36_2 | AGGGCTATCCTGGAAAAAGA | SLC25A36 |
| Slc25a36_3 | GAGTCTTTATAAGCCAAATG | SLC25A36 |
| Slc25a36_4 | TATCAGACAGACGGACTGCG | SLC25A36 |
| Slc25a37_1 | CTGGATTCATTACTGCATCG | SLC25A37 |
| Slc25a37_2 | CACTGTCCGGATACAACTGA | SLC25A37 |
| Slc25a37_3 | GATGAAGTGAATTGACTGGA | SLC25A37 |
| Slc25a37_4 | GCATCTATGGCGCCCTCAAG | SLC25A37 |
| Slc25a5_1 | CAAGGGCATCATAGACTGCG | SLC25A5 |
| Slc25a5_2 | CGCAGCCATCTCCAAGACAG | SLC25A5 |
| Slc25a5_3 | TTAAGATCTACAAATCTGAT | SLC25A5 |
| Slc25a5_4 | GCACAAGGATGTAGCCCCAG | SLC25A5 |
| Slc26a9_1 | TTGTGGGGGAGATCCGACAA | SLC26A9 |
| Slc26a9_2 | GGGTCTGGAAGATCACAACC | SLC26A9 |
| Slc26a9_3 | AGAGAGCGCAGCAAATGACG | SLC26A9 |
| Slc26a9_4 | TTCTAGACTCACAAAGACGA | SLC26A9 |
| Slc27a5_1 | GTGGGCTTAATGAACTATGT | SLC27A5 |
| Slc27a5_2 | TACCTCTGTACCATACGATA | SLC27A5 |
| Slc27a5_3 | GTAACAGTGATCTTGTATGT | SLC27A5 |
| Slc27a5_4 | CCTTTGTGGATGCTTTAGAG | SLC27A5 |
| Slc2a1_1 | CCTGCTCATCAATCGTAACG | SLC2A1 |
| Slc2a1_2 | TCAGCATGGAGTTCCGCCTG | SLC2A1 |
| Slc2a1_3 | GTGTCACCTACAGCTCTACG | SLC2A1 |
| Slc2a1_4 | CAAACATGGAACCACCGCTA | SLC2A1 |
| Slc2a3_1 | TTAGAAGACCTACCAAGTGA | SLC2A3 |
| Slc2a3_2 | GACCACGCCTGCTCCAATCG | SLC2A3 |
| Slc2a3_3 | CTGGAATGATGGTTAAGCCA | SLC2A3 |
| Slc2a3_4 | TGTGCCTATGTACATTGGAG | SLC2A3 |
| Slc2a5_1 | CTGGCCCCGAAAAACCTACG | SLC2A5 |
| Slc2a5_2 | TGCGCTGAGGTAGATCTGAT | SLC2A5 |
| Slc2a5_3 | CCTGCCCAGTTTATTCACCA | SLC2A5 |
| Slc2a5_4 | CGGGGACTCCAGTTAGACCC | SLC2A5 |
| Slc34a2_1 | TCATAGAGGAGCATCCCGAG | SLC34A2 |
| Slc34a2_2 | CTCCATCACCAACACGATCG | SLC34A2 |
| Slc34a2_3 | AGAGGTGCAGTTATCAGTCG | SLC34A2 |
| Slc34a2_4 | CTCACCGATGAGTGGAGTCA | SLC34A2 |
| Slc35a2_1 | TCACCCGCTGTAGTGGACCC | SLC35A2 |
| Slc35a2_2 | CTGCTCTTCGCACAAAAGAG | SLC35A2 |
| Slc35a2_3 | CTGCAAGGTATAGATGAGAG | SLC35A2 |
| Slc35a2_4 | GGCCACTGGATCAGAACCCG | SLC35A2 |
| Slc35f2_1 | GGCCGTGCCGCAAATACACA | SLC35F2 |
| Slc35f2_2 | TTGGTGCAGACATATTAGCT | SLC35F2 |
| Slc35f2_3 | TCTTGCCCGGAGAATAAACC | SLC35F2 |
| Slc35f2_4 | CCAGCATCAACGTGTAAACC | SLC35F2 |
| Slc38a1_1 | ATACTTTGGTGTGCACGCGT | SLC38A1 |
| Slc38a1_2 | TGCATGGTGTATGAGAAGCT | SLC38A1 |
| Slc38a1_3 | TCACCATCACCACCAACACT | SLC38A1 |
| Slc38a1_4 | AGATTGGCAGGACGGACGGG | SLC38A1 |
| Slc43a3_1 | TGACCGCTTCAAGACTACTG | SLC43A3 |
| Slc43a3_2 | GATTCATCTTGCACGTGGTG | SLC43A3 |
| Slc43a3_3 | TGCTCCAGAGCAATGTAACA | SLC43A3 |
| Slc43a3_4 | TACCCATAGCTGTAGTTGGG | SLC43A3 |
| Slc44a1_1 | CTGGAAGCAATACCGAACAG | SLC44A1 |
| Slc44a1_2 | GTACATGTGGTGGTACCACG | SLC44A1 |
| Slc44a1_3 | GTGGCACGGGTGTATTATGG | SLC44A1 |
| Slc44a1_4 | CACCATCGCCTTGTTCCACG | SLC44A1 |
| Slc44a2_1 | ATCAACAACCTTGTACACGG | SLC44A2 |
| Slc44a2_2 | GAACATTACAGATCTAGTGG | SLC44A2 |
| Slc44a2_3 | GGTGAAACGCATTACCTGAG | SLC44A2 |
| Slc44a2_4 | TCAAGTGCAGGTACACTCGG | SLC44A2 |
| Slc4a4_1 | AGCCTGCTGTAGGCGAACAA | SLC4A4 |
| Slc4a4_2 | GCCTCCAAAAGTGATGGCGT | SLC4A4 |
| Slc4a4_3 | ACTTTGAAGTCAGAGTTGAT | SLC4A4 |
| Slc4a4_4 | AGAGAATGTTCAGATGAATG | SLC4A4 |
| Slc5a6_1 | CTTACTTAATCCCAGAGATG | SLC5A6 |
| Slc5a6_2 | CAAACATGACCAGACCAATG | SLC5A6 |
| Slc5a6_3 | GCAGTTCACCAACGGTATGG | SLC5A6 |
| Slc5a6_4 | ATCCTATAGGTGATATACAT | SLC5A6 |
| Slc6a8_1 | GGAAGCGCCACACGTTACCG | SLC6A8 |
| Slc6a8_2 | GCATCAGTGTGACAGCCCGT | SLC6A8 |
| Slc6a8_3 | GCACAACGAGGACCACGTAG | SLC6A8 |
| Slc6a8_4 | ACTCGATGACAGGGGACCGG | SLC6A8 |
| Slc7a1_1 | GCCATGGCATAGATAACTCG | SLC7A1 |
| Slc7a1_2 | CACAAACGTGAAATACGGTG | SLC7A1 |
| Slc7a1_3 | TGACGTGAGAACTCTCCGAT | SLC7A1 |
| Slc7a1_4 | CCAGGTCCTTCAGTTCAAAG | SLC7A1 |
| Slc7a5_1 | GCCCTCCTCGCAGTACATCG | SLC7A5 |
| Slc7a5_2 | ACCCCTACTTACGCACGCAG | SLC7A5 |
| Slc7a5_3 | AGCGGCCTCTTCGCCTACGG | SLC7A5 |
| Slc7a5_4 | GTAGCAGAGTGCGCCCACGA | SLC7A5 |
| Slco4a1_1 | CATCCATCTCTACCTCATAG | SLCO4A1 |
| Slco4a1_2 | GCGCTACGTTGTTATGAGAG | SLCO4A1 |
| Slco4a1_3 | TGACCACTGACAGCCCACTG | SLCO4A1 |
| Slco4a1_4 | AGGTGCAGTATGTGTCCTCG | SLCO4A1 |
| Sms_1 | ACTTACTAACATCCCCGCTG | SMS |
| Sms_2 | TTACCACCCATAGTTCGCGG | SMS |
| Sms_3 | CTTACACGAACAAGAATGGC | SMS |
| Sms_4 | AAACAAGAAACTGACAGCGT | SMS |
| Soat1_1 | TGTGGTAAATTGTTCTGATG | SOAT1 |
| Soat1_2 | CAGTATCAGAATGAACCGGG | SOAT1 |
| Soat1_3 | CCCACCATTGTCCAGCGATG | SOAT1 |
| Soat1_4 | ACGAGTACTAAATGCAGCCA | SOAT1 |
| Soat2_1 | GCAATGGACTCGACATATGG | SOAT2 |
| Soat2_2 | ATAGGCCCGCTATGAACATG | SOAT2 |
| Soat2_3 | TCTACAGAGAGACATACCCC | SOAT2 |
| Soat2_4 | AGGACCCAAGAGTTACACCC | SOAT2 |
| Sod2_1 | GGCGTTGAGATTGTTCACGT | SOD2 |
| Sod2_2 | ATGATCTGCGCGTTAATGTG | SOD2 |
| Sod2_3 | ACAAACCTGAGCCCTAAGGG | SOD2 |
| Sod2_4 | CCTGCACTGAAGTTCAATGG | SOD2 |
| Srm_1 | TCTGCCCGCAGTAAAACCTA | Srm |
| Srm_2 | GGTCCAGTGCGAGATTGATG | Srm |
| Srm_3 | CGTGCCCTGGCTCACCCATG | Srm |
| Srm_4 | GCCCAGGTGCTGATCATCGG | Srm |
| Sptlc1_1 | AATGTGCCATAGAACCCTCG | SPTLC1 |
| Sptlc1_2 | CCCTCCAACCCACAACATCG | SPTLC1 |
| Sptlc1_3 | TCCTGCGTACTCTAAGAGAG | SPTLC1 |
| Sptlc1_4 | TTTGTCGTAGAATCCTCGCA | SPTLC1 |
| Sptlc2_1 | GTTGTGTTTGAAGATTCGAA | SPTLC2 |
| Sptlc2_2 | TGAGAGCAATCACTTCAGGA | SPTLC2 |
| Sptlc2_3 | AATCTCGAAGATATCCAAAG | SPTLC2 |
| Sptlc2_4 | ACAACTATCTTGGATTTGCG | SPTLC2 |
| Sptlc3_1 | AGGGAAATATGATGATTCGA | SPTLC3 |
| Sptlc3_2 | GAAGATATTGACGTATACAT | SPTLC3 |
| Sptlc3_3 | TTCTCAAAGTCCTGATACAG | SPTLC3 |
| Sptlc3_4 | CTGAGAGAAGCTATCATCCG | SPTLC3 |
| Sptssa_1 | CAGGTACTGGTAGTAGAACC | SPTSSA |
| Sptssa_2 | CTGAACACGGTTCGCTCCCA | SPTSSA |
| Sptssa_3 | CCATCACGCAGATTCGATGC | SPTSSA |
| Sptssa_4 | GTACAGGGCCATCCCCACCA | SPTSSA |
| Sptssb_1 | CATGGATTTCAAGCGCGTGA | SPTSSB |
| Sptssb_2 | CTGGTATTGATAATAGAGCC | SPTSSB |
| Sptssb_3 | CACCATCATACTGACCATTG | SPTSSB |
| Sptssb_4 | TTGAGCATCGATTGTTCCCA | SPTSSB |
| Sqle_1 | GGACCCGGAAGTGATCATCG | SQLE |
| Sqle_2 | AGGGTACTTGTTGATATTCG | SQLE |
| Sqle_3 | GAAACCAACCAAGTGCAGAG | SQLE |
| Sqle_4 | CGCTGTCGCCATCGACACGG | SQLE |
| Srd5a1_1 | TCGGCGGCCGGACCACTGCG | SRD5A1 |
| Srd5a1_2 | TGCGGTGTATGCTGAAGACT | SRD5A1 |
| Srd5a1_3 | GAAACATAGCTAGCAGGACG | SRD5A1 |
| Srd5a1_4 | TCACCATGCCCACTAACCAC | SRD5A1 |
| Sucla2_1 | CAGAGCGTAACATACTGTCA | SUCLA2 |
| Sucla2_2 | TGTGCACTTCCTATCAGTAC | SUCLA2 |
| Sucla2_3 | GTAGAAGATTCTGACGGAAA | SUCLA2 |
| Sucla2_4 | TCAAGTATTCATGCAGCGAA | SUCLA2 |
| Sulf1_1 | CAGTTGACGACTCTGTCGAG | SULF1 |
| Sulf1_2 | AGAAATACGTACATCGACCC | SULF1 |
| Sulf1_3 | GCACGCAACCTCTACTCTCG | SULF1 |
| Sulf1_4 | CTGGCGTGATACATTCCTAG | SULF1 |
| Sulf2_1 | GACGGTGAGATATACCACGT | SULF2 |
| Sulf2_2 | ACAAATGTAAAGGCCCCATG | SULF2 |
| Sulf2_3 | ACATGGCTCAGACTACTCCA | SULF2 |
| Sulf2_4 | GCACATTGGTGTAGTCACGA | SULF2 |
| Taldo1_1 | GCGCATCCTTGATTGGCATG | TALDO1 |
| Taldo1_2 | CTTCTTTGTAAAGCTCGATG | TALDO1 |
| Taldo1_3 | TTCTGAATTCAGGCCTCAAG | TALDO1 |
| Taldo1_4 | GCTTGTATTCATCGATGGCT | TALDO1 |
| Tap1_1 | ACTAATGGACTCGCACACGT | TAP1 |
| Tap1_2 | GTCTCTAGCAAAGTCCACGC | TAP1 |
| Tap1_3 | TGCCACATAACTGATAGCGA | TAP1 |
| Tap1_4 | TGGACATGAGCCATATGTTG | TAP1 |
| Tat_1 | ATGTTCGCGTCAATATTGGC | TAT |
| Tat_2 | GGCAAGCTTACCGATAGATG | TAT |
| Tat_3 | AACAACCCGTCCAATCCCTG | TAT |
| Tat_4 | TGAGCTGTGTCTAGCCGTGT | TAT |
| Tdo2_1 | GATAGCTCGGATGCATCGTG | TDO2 |
| Tdo2_2 | TGATGAATAGGTGCTCGTCA | TDO2 |
| Tdo2_3 | AATCCATTTGGCTCTAAACC | TDO2 |
| Tdo2_4 | CACTATCGTGATAACTTTGG | TDO2 |
| Th_1 | GGGTGAGCCAATTCCCCACG | TH |
| Th_2 | ACTGTGTGCACTGAAACACA | TH |
| Th_3 | CCCCAAGGTTCATTGGACGG | TH |
| Th_4 | GTGCGCTTCGAGGTGCCCAG | TH |
| Tk1_1 | TAGGACTGACCGATCATGTG | TK1 |
| Tk1_2 | CAGGCCCAGCCTCTTCGTGT | TK1 |
| Tk1_3 | AGGACTCCTGGGTCACATCG | TK1 |
| Tk1_4 | GTAATTGTGGCAGCGCTGGA | TK1 |
| Tk2_1 | CTCCAATACAACAGACGTCG | TK2 |
| Tk2_2 | TTGAGGGCAATATTGCAAGT | TK2 |
| Tk2_3 | AGAATCGCGTAGTCAACCTC | TK2 |
| Tk2_4 | TACCATGATGCCAGCCGATG | TK2 |
| Tkt_1 | CGTGGACGGACACAGCGTGG | TKT |
| Tkt_2 | CAGCGCTGCAGCATGATGTG | TKT |
| Tkt_3 | CCCACAGATAGCCACCCGGA | TKT |
| Tkt_4 | CTCCGAGGGCTCCGTCTGGG | TKT |
| Tph1_1 | AGAGGTAACCAGCCACAGGA | TPH1 |
| Tph1_2 | CGGAAGAAGAGATTAAGACC | TPH1 |
| Tph1_3 | GTGGCGTGCGACTTCAGCAG | TPH1 |
| Tph1_4 | TGGGAGAATTGAGCAAAACT | TPH1 |
| Tph2_1 | AGGACAATGTCTATCGACAG | TPH2 |
| Tph2_2 | TGCTTCATATCGAATCCAGG | TPH2 |
| Tph2_3 | CCCCTGCTGACCAAGTACTG | TPH2 |
| Tph2_4 | GCCTGAGAGCATTTGGACGG | TPH2 |
| Tpi1_1 | TGAAGGTCAGTACAAACGCA | TPI1 |
| Tpi1_2 | AAGTCGATGTAAGCGGTGGG | TPI1 |
| Tpi1_3 | AGTGAGCCACGCCCTAGCAG | TPI1 |
| Tpi1_4 | CCAACGAAGAACTTCCTGGT | TPI1 |
| Trpm2_1 | CGTGGCCTGTCAGTGTACCG | TRPM2 |
| Trpm2_2 | GTTCATCTCAGAGCAAACGA | TRPM2 |
| Trpm2_3 | AGGAAGGTAGTGTGTGCGTG | TRPM2 |
| Trpm2_4 | AGTAGGAGAGGATATTCATG | TRPM2 |
| Tsta3_1 | TCGGGTACTCATACGCCAAG | TSTA3 |
| Tsta3_2 | CATCTCGCTGCAATGGTAGG | TSTA3 |
| Tsta3_3 | TGTCTCGTCAATAGGATAGG | TSTA3 |
| Tsta3_4 | GAAGATGGCCACGTGCTACC | TSTA3 |
| Ttyh3_1 | GGAAATGCAAGATGTTGTCG | TTYH3 |
| Ttyh3_2 | GAGCACTCAGTACTGAGTGG | TTYH3 |
| Ttyh3_3 | GAACCCCGGAGTATCGCTGG | TTYH3 |
| Ttyh3_4 | ACGCCATGCCAACCGCACAG | TTYH3 |
| Tyms_1 | CAGGCACGATACAGCCTGAG | TYMS |
| Tyms_2 | GACAATTCTACAGATTACTC | TYMS |
| Tyms_3 | TCACCACATAGAACTGACAG | TYMS |
| Tyms_4 | TTCAAGAAGGAGGACCGCAC | TYMS |
| Tyr_1 | AGAAATTCGAGAACTAACTG | TYR |
| Tyr_2 | TTTATGCGATGGAACACCTG | TYR |
| Tyr_3 | ATGTTGATATCATTAAACAT | TYR |
| Tyr_4 | ACCCCTTTGAAGGGGAACTG | TYR |
| Uck1_1 | ACCACCTGCTTACCTTGAGT | UCK1 |
| Uck1_2 | GATGTTAGGCTGTCTCGAAG | UCK1 |
| Uck1_3 | ACATTCTCCAGGTACCTGGG | UCK1 |
| Uck1_4 | TGCCGCCGCTCACGCCGATG | UCK1 |
| Uck2_1 | GGGAGACAAAGTCGTACACG | UCK2 |
| Uck2_2 | GATACCCGTCTGTCTCGCCG | UCK2 |
| Uck2_3 | CCCTTCGAAGAGCACCACGT | UCK2 |
| Uck2_4 | CAATTCAACTTTGATCACCC | UCK2 |
| Uggt1_1 | CAAGTGGGTCAACAACCTAG | Uggt1 |
| Uggt1_2 | GGAGAAGAAGTACCCGTACG | Uggt1 |
| Uggt1_3 | CATTGCGGAATTCTCTGTCG | Uggt1 |
| Uggt1_4 | TCGTGTGACAGGGTCAACAA | Uggt1 |
| Ugdh_1 | GAAGTAGTCGAATCCTGTCG | UGDH |
| Ugdh_2 | CATTTGAGTTCTGCACAATG | UGDH |
| Ugdh_3 | TGAAGCAACAGGCGCCGATG | UGDH |
| Ugdh_4 | TGTTGGCATCAAATATGCGG | UGDH |
| Ugt1a1_1 | AGACAAACTCTTGGGCACGT | UGT1A1 |
| Ugt1a1_2 | CAGTGGCAAGTGACCCATAG | UGT1A1 |
| Ugt1a1_3 | ACTTTGTGAAAGATTACCCC | UGT1A1 |
| Ugt1a1_4 | ACACTAACAGCCTCCCAGCG | UGT1A1 |
| Upp1_1 | GCTACGCCATGTATAAAGCC | UPP1 |
| Upp1_2 | GGGCCTTGACCATCCAGGGA | UPP1 |
| Upp1_3 | TCCCGCCTGAAGTGCCAATG | UPP1 |
| Upp1_4 | GGTGATTCGCAACACCAACC | UPP1 |
| Uqcrh_1 | TGGATCTGGAGACCCCAAAG | UQCRH |
| Uqcrh_2 | GACGAACGAAAGATGCTCAC | UQCRH |
| Uqcrh_3 | AATCCTCTTCTGTCTGTGAC | UQCRH |
| Uqcrh_4 | GCTCTCTCACTGTTGTTAGG | UQCRH |
| Uros_1 | TCTCCTTTGATAGTTCCACA | UROS |
| Uros_2 | TGACAGCACAGGAATCAGTG | UROS |
| Uros_3 | GCCAAGTCTGTGTACGTGGT | UROS |
| Uros_4 | AGACATGCATGCTTTCCATG | UROS |
| Vdac1_1 | AAGAGGGAGCACATCAACCT | VDAC1 |
| Vdac1_2 | CGGCGAGAATGACGAATCAA | VDAC1 |
| Vdac1_3 | TGGAACACAGACAACACCCT | VDAC1 |
| Vdac1_4 | CTTCGCAGTTGGCTATAAGA | VDAC1 |
| Xdh_1 | CCAGCATGCAACGTACAGGG | XDH |
| Xdh_2 | CATACTCATGACAATACCAG | XDH |
| Xdh_3 | TCAAAACGCAACGTCTTCCG | XDH |
| Xdh_4 | GGAATTCCACCATCCCACCA | XDH |
| Ppara_1 | CATCGAGTGTCGAATATGTG | Ppara |
| Ppara_2 | GATTTCTCAGTCCATCGGTG | Ppara |
| Ppara_3 | TCTGTCGGGATGTCACACAA | Ppara |
| Ppara_4 | CTGCTGCCAGTGCATGTCCG | Ppara |
| Ppard_1 | CCTCCGGCATCCGTCCAAAG | Ppard |
| Ppard_2 | GGCCTCGGGCTTCCACTACG | Ppard |
| Ppard_3 | TCGAGTATGAGAAGTGCGAT | Ppard |
| Ppard_4 | CCTCAAGTATGGCGTGCACG | Ppard |
| Pparg_1 | AATGCTGGAGAAATCAACTG | Pparg |
| Pparg_2 | AGAACCTTCTAACTCCCTCA | Pparg |
| Pparg_3 | GCACCCTTGAAAAATTCGGA | Pparg |
| Pparg_4 | CTGCCTATGAGCACTTCACA | Pparg |
| Ppargc1a_1 | GAATGAGGCAAACTTGCTAG | Ppargc1a |
| Ppargc1a_2 | TATTGAGCGAACCTTAAGTG | Ppargc1a |
| Ppargc1a_3 | TTGCATGCGCACCTTAACAA | Ppargc1a |
| Ppargc1a_4 | AAGACCAGTGAACTAAGGGA | Ppargc1a |
| Ppargc1b_1 | AGGGCTTGCTAACATCACAG | Ppargc1b |
| Ppargc1b_2 | TGGACGAGCTTTCACTGGTG | Ppargc1b |
| Ppargc1b_3 | GGCCTTGACTACTGTCTGTG | Ppargc1b |
| Ppargc1b_4 | TCCTGTGCTAGGGAGTCTTG | Ppargc1b |
| Slc1a5_1 | AATCCCTATCGATTCCTGTG | Slc1a5 |
| Slc1a5_2 | TACAACAGAGTCGTTGATGG | Slc1a5 |
| Slc1a5_3 | GCGGGAGATCAATTCAACCA | Slc1a5 |
| Slc1a5_4 | GTGGTGTGCAGCCTGATCGG | Slc1a5 |
| Prkaa1_1 | GAAGATTCGGAGCCTTGACG | Prkaa1 |
| Prkaa1_2 | GATCGGCCACTACATCCTGG | Prkaa1 |
| Prkaa1_3 | ATCACCATGAAAATATCAGA | Prkaa1 |
| Prkaa1_4 | CCTGTGACAATAATCCACAC | Prkaa1 |
| Me1_1 | ATGGCCAGAGGATGTCGTCA | Me1 |
| Me1_2 | TGGTTATTCGAAGAGCTGCA | Me1 |
| Me1_3 | GTTCCCACATCCAAAGTGAT | Me1 |
| Me1_4 | CCAGGAAGGCGTCATACTCA | Me1 |
| Me2_1 | CCAGGAACCATATGCCCATG | Me2 |
| Me2_2 | ATTGTGTACACGCCAACAGT | Me2 |
| Me2_3 | GTGCCCACGTCGATGCACAC | Me2 |
| Me2_4 | AGGTCATGTTAGATCAATTG | Me2 |
| Me3_1 | AGTTCAGCATTGTTGCAATG | Me3 |
| Me3_2 | GAAACACCAGCGTGTCCGTG | Me3 |
| Me3_3 | AGCGCGGATACGATGTCACC | Me3 |
| Me3_4 | GGAAGGCATACCCAAGACAG | Me3 |
| Bcat1_1 | TAGAAAAATAAGGTCCCACG | Bcat1 |
| Bcat1_2 | CCCCACATTTATCGGAACTG | Bcat1 |
| Bcat1_3 | CACAGCGTAGTGCAAAACAG | Bcat1 |
| Bcat1_4 | CAAGATCCGATTGTTCCGGC | Bcat1 |
| Bcat2_1 | ACGGAACGAGCCTCTACGTG | Bcat2 |
| Bcat2_2 | CAGGAACTATGGACCCACTG | Bcat2 |
| Bcat2_3 | GTGGAGTGGAATAACAAGGC | Bcat2 |
| Bcat2_4 | GCACAGAATGACGTACAGGA | Bcat2 |
| Cd36_1 | AAATATAACTCAGGACCCCG | CD36 |
| Cd36_2 | CCAAAACTGTCTGTACACAG | CD36 |
| Cd36_3 | TAGGATATGGAACCAAACTG | CD36 |
| Cd36_4 | TTAATCATGTCGCAATAGCT | CD36 |
| Mpc1_1 | ACTTCCGGGACTATCTCATG | Mpc1 |
| Mpc1_2 | GGCGGACTATGTCCGGAGCA | Mpc1 |
| Mpc1_3 | AAATCTCCAGAGATTATCAG | Mpc1 |
| Mpc1_4 | TGGGGCCCAGTTGCCAACTG | Mpc1 |
| Mpc2_1 | CCTGCCGGGTGGTTGTAAAG | Mpc2 |
| Mpc2_2 | CCACTTTATCCATGAGTCGG | Mpc2 |
| Mpc2_3 | CCCAGAAAAAAACTGTTCTG | Mpc2 |
| Mpc2_4 | ATTGGTGTGTGCTGGACTAG | Mpc2 |
| Atg5_1 | AAGAGTCAGCTATTTGACGT | Atg5 |
| Atg5_2 | AAATGTACTGTGATGTTCCA | Atg5 |
| Atg5_3 | CCTTCTACACTGTCCATCCA | Atg5 |
| Atg5_4 | AAGAAAAACTCACCATTTCA | Atg5 |
| Nnmt_1 | GCCTTCAAGATCACACACAT | Nnmt |
| Nnmt_2 | GTTGAGGCGTGCAATCAAGC | Nnmt |
| Nnmt_3 | GGAGAACTCCTGATTGACAT | Nnmt |
| Nnmt_4 | TGCGATAGGCCGGGAGGTCA | Nnmt |
| Glud1_1 | CAGTAGCGGAGATGCGCCCG | Glud1 |
| Glud1_2 | CCGCGGCGCCAGCATCGTAG | Glud1 |
| Glud1_3 | ACATACAAGTGCGCTGTGGT | Glud1 |
| Glud1_4 | GGAAGGAGAGGCTCAACACA | Glud1 |
| Got1_1 | GATCCCCGCAAGGTTAACCT | Got1 |
| Got1_2 | GTTGGTGATGATACGTAGAT | Got1 |
| Got1_3 | AGACCTAGAGAAAGATGCGT | Got1 |
| Got1_4 | CATTCGGCCCTATTGCTACT | Got1 |
| Got2_1 | TGGAGGTCCCATTTCAACAT | Got2 |
| Got2_2 | TTTCTGCCCAAACCATCCTG | Got2 |
| Got2_3 | CATCCTCCTCACCTTCACCA | Got2 |
| Got2_4 | AGCTCACCTTCCGGACACTG | Got2 |
| Gpt2_1 | GCGGTGGAGTACGCTGTGCG | Gpt2 |
| Gpt2_2 | ACGCTAAGAAACGAGCGCGG | Gpt2 |
| Gpt2_3 | GTTCTCTGCATTATCAACCC | Gpt2 |
| Gpt2_4 | GGGGATGGGAATCATCACGC | Gpt2 |
| Mtap_1 | TGTGGATACTCCATTCGGCA | Mtap |
| Mtap_2 | GGACAATAGTCACAATTGAG | Mtap |
| Mtap_3 | GGCAAAACGGTTCAGCCATG | Mtap |
| Mtap_4 | GCCTTCAAAAGTCAACTACC | Mtap |
| Aldh9a1_1 | AGCAAGTGTCAACATCCAGG | ALDH9A1 |
| Aldh9a1_2 | GCACAGAACCGACTGCACCG | ALDH9A1 |
| Aldh9a1_3 | GCCCCGCCGCGGTAGTTGAG | ALDH9A1 |
| Aldh9a1_4 | GCGCTGCCAAGTCCTCCTAG | ALDH9A1 |
| Bckdhb_1 | AGCCTACCTGATCAAAGGCA | BCKDHB |
| Bckdhb_2 | AGTTGAAAAGATCACCTGAG | BCKDHB |
| Bckdhb_3 | CCATGGCCCACACAACCCCA | BCKDHB |
| Bckdhb_4 | GAGAATGGTAGAGAGCCCCA | BCKDHB |
| Eno1_1 | AAGGCTTTCAATGTGATCAA | ENO1 |
| Eno1_2 | CCTTACCCTTCCCCATGAAG | ENO1 |
| Eno1_3 | GTACAGATCGACCTCAACAG | ENO1 |
| Eno1_4 | GTTCCTTCTAGGTGTCTCAC | ENO1 |
| Gpam_1 | GAAATACCTGATCATCCCTG | GPAM |
| Gpam_2 | GACATCCTCGTCATACCCGT | GPAM |
| Gpam_3 | GAGGCGTTATCAGAATGCTG | GPAM |
| Gpam_4 | GCAGTCCAAAGCCATCCAGA | GPAM |
| Gsta1_1 | CTTCACTACTTCAATGCCCG | GSTA1 |
| Gsta1_2 | GATGCACTCCATTCTGCCCC | GSTA1 |
| Gsta1_3 | GATGTTTGACCAAGTGCCCA | GSTA1 |
| Gsta1_4 | GCTTCACTACTTCAATGCCC | GSTA1 |
| Hsd3b1_1 | AGGTACCCAGAACCTATTGG | HSD3B1 |
| Hsd3b1_2 | ATTCCGACCAGAAACCAAGG | HSD3B1 |
| Hsd3b1_3 | ATTCCTCCTTGGTTTCTGGT | HSD3B1 |
| Hsd3b1_4 | CTTGAACACAGGCCTCCAAT | HSD3B1 |
| Pank4_1 | ACTTGCCTACCTTTCCCGGG | PANK4 |
| Pank4_2 | CTATGAGATCTCAGTCCAGG | PANK4 |
| Pank4_3 | GGCTCTGCTCACCAAAACAA | PANK4 |
| Pank4_4 | GTGGACTCTTACCTTCACGA | PANK4 |
| Pfkfb1_1 | AAGCCTCTAAGAGAGCAGGT | Pfkfb1 |
| Pfkfb1_2 | CCATAAGTATCTCAGCCGTG | Pfkfb1 |
| Pfkfb1_3 | CCGCAACGTGACCTTCCTCA | Pfkfb1 |
| Pfkfb1_4 | CTTTCGCCCAGACAACATGG | Pfkfb1 |
| Pip5k1b_1 | AAGAGCATCCCGAAAAGAGA | PIP5K1B |
| Pip5k1b_2 | AGTCCAGGTCCTTAAACGTG | PIP5K1B |
| Pip5k1b_3 | GAGTTCTCGGAAGTACCGGA | PIP5K1B |
| Pip5k1b_4 | GCTGGGAATAGGATACACAG | PIP5K1B |
| Pikfyve_1 | ATAATCTTGATGCTTCAAAG | Pikfyve |
| Pikfyve_2 | CAGAAGACAGGGCCTCACGA | Pikfyve |
| Pikfyve_3 | CTATATGCTGACTTGAAGGG | Pikfyve |
| Pikfyve_4 | GATGTGTTCTCAACCAGTAA | Pikfyve |
| Pla2g4a_1 | ATAATGATATAAACCCCGTG | PLA2G4A |
| Pla2g4a_2 | CCACTTTCATTGAAGATACA | PLA2G4A |
| Pla2g4a_3 | CGTGCCACCAAAGTAACCAA | PLA2G4A |
| Pla2g4a_4 | GTGCCACCAAAGTAACCAAG | PLA2G4A |
| Tymp_1 | AATCCTCAGTAAGAAGGCCG | TYMP |
| Tymp_2 | CACCAGGTGCCAATGATCAG | TYMP |
| Tymp_3 | CTAATGGCCATTCGGCTGCA | TYMP |
| Tymp_4 | GCAGATGCTGCACATCCTTG | TYMP |
| Pla2g2a_1 | CAAGGGAACATTGCGCAGTT | PLA2G2A |
| Pla2g2a_2 | CTTACCAAAGGCCATGATCG | PLA2G2A |
| Pla2g2a_3 | GCTGCTAGCAGCCTCGATCA | PLA2G2A |
| Pla2g2a_4 | GGGGGAATCCTTTGCCACCC | PLA2G2A |
| Ppp1r8_1 | GATGACGAGATCATCAACCC | Ppp1r8 |
| Ppp1r8_2 | AATGAGACCGTAGAATCGAT | Ppp1r8 |
| Ppp1r8_3 | GGACTCGAGAGCATGACTGG | Ppp1r8 |
| Ppp1r8_4 | TCAACACTGCCCACAACAAG | Ppp1r8 |
| Gtf2b_1 | TATTACCTTGCCAATCATGG | Gtf2b |
| Gtf2b_2 | TTACAACTATATTGCGAGGG | Gtf2b |
| Gtf2b_3 | ATAGCTACTTACCTACAACT | Gtf2b |
| Gtf2b_4 | AGTTGTGATCAGATCCACGC | Gtf2b |
| Rps3a1_1 | CATCATTCTGTAGATCAGCA | Rps3a1 |
| Rps3a1_2 | GATGTTCCTAATATTGAACA | Rps3a1 |
| Rps3a1_3 | CTCCATGGTCAAGAAGTGGC | Rps3a1 |
| Rps3a1_4 | AAAATTCAAGCTAATCACTG | Rps3a1 |
| Cstf1_1 | CCGAACTCTTTACGACCACG | Cstf1 |
| Cstf1_2 | GATGTCATACAGGCGAAGCG | Cstf1 |
| Cstf1_3 | CCACAAAGGCCCGTGCCGCG | Cstf1 |
| Cstf1_4 | GGTTACATTAGTATCGCCAA | Cstf1 |
| Tuba1c_1 | GAGCCGCTCCATCAGCAGGG | Tuba1c |
| Tuba1c_2 | ACACAATCTGGCTAATAAGG | Tuba1c |
| Tuba1c_3 | CACAGCAGTGGAAACCTGGG | Tuba1c |
| Tuba1c_4 | CCAGAGGGAAGTGGATGCGA | Tuba1c |
| Cdc45_1 | TGACCACGTCCAGTATACGC | Cdc45 |
| Cdc45_2 | GCAAAGATACTCATACTCGA | Cdc45 |
| Cdc45_3 | CATAAACTGTGGAGCCAACG | Cdc45 |
| Cdc45_4 | ATCTCTAAAGATGTCGTCGT | Cdc45 |
| Rps15a_1 | CTCACCGTGCTTCATCATCA | Rps15a |
| Rps15a_2 | CCTCACAGGAAGGTTGAACA | Rps15a |
| Rps15a_3 | CTTTAGAACATGGCCTGATG | Rps15a |
| Rps15a_4 | ATCAACAACGCTGAGAAGAG | Rps15a |
| Cul1_1 | TCGCTTCATAAACAACAATG | Cul1 |
| Cul1_2 | GCACACAAGATGAACTAGCA | Cul1 |
| Cul1_3 | GAAGTTCTACACTCAGCAGT | Cul1 |
| Cul1_4 | CTTGCAGCAATTGAGAAGTG | Cul1 |
| Rfc4_1 | AGAATTACAATCTTAAAGGG | Rfc4 |
| Rfc4_2 | GGCAGCTTTAAGGCGTACCA | Rfc4 |
| Rfc4_3 | CTCCAGCCGTCCAAAGTGTG | Rfc4 |
| Rfc4_4 | TTAAATGCATCTGATGAACG | Rfc4 |
| Cenpl_1 | ATATAGTCCACTGTTTGTGG | Cenpl |
| Cenpl_2 | ATTCCAGTCGTTTCTGCAGA | Cenpl |
| Cenpl_3 | ATTGTTGCCGAAAAGCAAAA | Cenpl |
| Cenpl_4 | CATTTCTCCTCCACAAACAG | Cenpl |
