## Supplemental Table 5 for "Recurrent Breast Cancer Cells Depend on *De novo* Pyrimidine Biosynthesis to Suppress Ferroptosis"

Table S5. Reagents and chemicals.

| **Antibodies** | | |  |
| --- | --- | --- | --- |
| **Protein** | **Supplier** | **Catalog #** | **Concentration** |
| Phospho-histone H2A.X (ser139) | Cell Signaling Technology | 2577 | 1:1,000 |
| Histone H2A | Cell Signaling Technology | 3636 | 1:1,000 |
| Phospho-histone H3 (ser10) | Cell Signaling Technology | 3377 | 1:1,000 |
| α-tubulin | Cell Signaling Technology | 3873 | 1:1,000 |
| HER2/ErbB2 | Cell Signaling Technology | 4290 | 1:1,000 |
| Nrf2 | Cell Signaling Technology | 12721 | 1:1,000 |
| Goat anti-Rabbit Alexa Fluor 680 | Invitrogen | A21109 | 1:15,000 |
| Goat anti-Mouse IRDye^®^ 800CW | Li-Cor | 926-32210 | 1:15,000 |

| **Taqman^®^ gene expression probes** | | |
| --- | --- | --- |
| Taqman^®^ Gene Expression Master Mix | Applied Biosystems #4369016 | |
| Taqman^®^ Universal PCR Master Mix | Applied Biosystems #4304437 | |
| **Gene** | **Species** | **Catalog #** |
| Her2/Erbb2 | rat | Rn00566561_m1 |
| Slc28a1 | mouse | Mm013153355_m1 |
| Slc28a1 | human | Hs00984391_m1 |
| Slc28a2 | mouse | Mm04212034_m1 |
| Slc28a2 | human | Hs01035846_m1 |
| Slc28a3 | mouse | Mm00491586_m1 |
| Slc28a3 | human | Hs00223220_m1 |
| Slc29a1 | mouse | Mm01270577_m1 |
| Slc29a1 | human | Hs01085706_m1 |
| Slc29a2 | mouse | Mm00432817_m1 |
| Slc29a2 | human | Hs01546959_g1 |
| Slc29a3 | mouse | Mm00469915_m1 |
| Slc29a3 | human | Hs00983219_m1 |
| Slc29a4 | mouse | Mm00525575_m1 |
| Slc39a4 | human | Hs00928283_m1 |
| Tbp | mouse | Mm00446971_m1 |
| Tbp | human | Hs00427620_m1 |

| **Cell treatment compounds** | | |
| --- | --- | --- |
| **Compound** | **Supplier** | **Catalog #** |
| DMSO | Sigma | D2650 |
| BAY 2402234 | Selleckchem | S9947 |
| Uridine | Sigma | U3750 |
| Lapatinib | Selleckchem | S1028 |

| **Plasmids** | | | |
| --- | --- | --- | --- |
| **Name** | **Supplier** | **Catalog #** | **Additional Notes** |
| pLV[Exp]-Puro-EF1A>mSlc28a3[NM_022317.3]*/HA | VectorBuilder | VB220816-1457bvq |  |
| pLV[Exp]-Neo-EF1A>mSlc29a4[NM_146257.2]* | VectorBuilder | VB220901-1344eup |  |
| lentiGuide-Puro | Addgene | 52963 | Gift from Feng Zhang |
| pLenti-PGK-Neo-PIP-FUCCI | Addgene | 118616 | Gift from Jean Cook |
| psPAX2 | Addgene | 12260 | Gift from Didier Trono |
| pMD2.G | Addgene | 12259 | Gift from Didier Trono |
